# Unique Microglial Transcriptomic Signature within the Hippocampal Neurogenic Niche

**DOI:** 10.1101/2021.08.29.457876

**Authors:** Sana Chintamen, Pallavi Gaur, Nicole Vo, Elizabeth M. Bradshaw, Vilas Menon, Steven G. Kernie

**Author notes:** Correspondence should be addressed to Steven G. Kernie. Authors contributed equally (Sana Chintamen, Pallavi Gaur).

## Abstract

Microglia, the resident immune cells of the brain, are crucial in the development of the nervous system. Recent evidence demonstrates that microglia modulate adult hippocampal neurogenesis by inhibiting cell proliferation of neural precursors and survival both *in vitro* and *in vivo*, thus maintaining a balance between cell division and cell death in the neural stem cell pool. There are increasing reports suggesting these microglia found in neurogenic niches differ from their counterparts in non-neurogenic areas. Here, we present evidence that microglia in the hippocampal neurogenic niche are a specialized population that express genes known to regulate neurogenesis. By comprehensively profiling myeloid lineage cells in the hippocampus using single cell RNA-sequencing, we resolve transcriptomic differences in microglia originating from the subgranular zone. These cells have lower expression of genes associated with homeostatic microglia and increased expression of genes associated with phagocytosis. Intriguingly, this small yet distinct population expresses a gene signature with substantial overlap with previously characterized phenotypes, including disease associated microglia (DAM), a particularly unique and compelling microglial state.

## Introduction

The hippocampus is important for memory consolidation as well as declarative and spatial memory and learning^1–3^. This region is also known to be affected early or more severely in a variety of neurodegenerative and psychiatric diseases. These include Alzheimer’s disease (AD), epilepsy, and major depressive disorder, which all are known to exhibit alterations in immune activity and each manifest hallmark traits of inflammation^4–7^. Subsets of microglia show proclivity towards disease progression in both the rodent and human brain ^8–10^. Thus, characterizing various immune subsets in the hippocampus is crucial for uncovering mechanisms of disease development and progression.

During early postnatal development, the brain is highly plastic and microglia exhibit a great degree of heterogeneity. In contrast, previous studies in the adult rodent brain have shown limited heterogeneity, corresponding to a time point when the brain is less plastic^10–12^. However, since neurogenic niches undergo life-long development, immune cells show phenotypic differences that correlate with a specialized need to support these regions ^13–15^. Increasing evidence demonstrates that microglia and other immune cells actively regulate adult hippocampal neurogenesis^16, 17^, and in fact, immune input has been shown to alter neurogenesis during injury, stroke, and aging^18^. Importantly, adult hippocampal neurogenesis (AHN) is key in certain forms of spatial memory and learning, memory consolidation, and recovery from injury^19^. Deficits in AHN in the murine and human brain have been found in a host of neurodegenerative diseases such as depression, Alzheimer’s Disease, and age-associated cognitive deficits^20–23^. This suggests that attenuating reductions in neurogenesis may prevent the cognitive decline associated with aging or neurodegeneration^24^.

Bulk sequencing experiments show subtle differences in various genes between subregions in the hippocampus^13^. Single cell transcriptomic profiling of cells in the dentate gyrus has demonstrated that immune cells minimally express common microglia markers and more highly express some genes associated with microglial activation^25^. This necessitates a direct comparison between various populations within the hippocampus at the single-cell level to provide relative information on how immune cells are specialized to support the neurogenic niche. In this study, we leverage transcriptomic profiles from myeloid lineage cells in the hippocampus at the level of single cells to resolve heterogeneity previously obscured in bulk sequencing/profiling.

Our experimental paradigm profiles over 18,000 cells from twelve murine hippocampi to resolve heterogeneity in the myeloid landscape of the adult hippocampus. In doing so, we have a substantially higher number and resolution of hippocampal myeloid cells than previously reported^11, 13, 25^. Consequently, we uncovered rare populations that reside in the hippocampus and have previously not been identified. Here, we identify a unique subset or population of cells that correspond to myeloid cells in the subgranular zone, which shape, regulate and/or support the pool of hippocampal neural progenitor cells. By examining these cells within the myeloid cell pool in the hippocampus, not only are we able to make a direct comparison to other subsets of microglia, but we also can examine other populations that may influence the neurogenic niche, even when not in direct contact with stem/progenitor cells in this region. This novel and comprehensive transcriptomic study, with single cell resolution, uniquely highlights genes involved in immune activation and neuronal development and support that provides insight into how the neurogenic niche is regulated in development and disease.

## Materials and Methods

### Animals

All experimental procedures were in accordance with the Guide for the Care and Use of Laboratory Animals of the National Institutes of Health and approved by the Institutional Animal Care and Use Committee at Columbia. Experimental animals were humanely housed and cared for under the supervision of the Institute of Comparative Medicine at Columbia University. For generation of the dual reporter mice, Cx3Cr1CreERT2+/+ (Jackson stock no. 021160) males were bred with Rosa26-loxp-stop-tdtomato+/+ females, resulting in progeny (F1) that were heterozygous for both the Cre recombinase and the flox-stop tdTomato reporter (Jackson stock 007914). Mice from F1 were crossed mice from F2 that were homozygous for the CreERT2 and tdTomato were selected as breeders. Finally, these mice were crossed with Nestin-GFP mice developed by us (Jackson stock no. 02967), resulting in progeny (F3) that were heterozygous for each of the 3 alleles of interest, the Cx3Cr1CreERT2, Rosa26-tdtomato, and TK-Nestin-eGFP. See Supplemental Figure 1^26^.

#### Microglial Isolation

Seven-week old dual reporter mice mice were injected with Tamoxifen (100mg/kg) once a day for four consecutive days. At week eight, 2 females and 2 males were sacrificed. Mice were perfused with approximately 25-30 mL of ice-cold sterile PBS (Corning Cellgro REF 21-040-CV). Each brain was extracted whole and placed in 5 mL of homogenization buffer (see buffers list) at 4% while other mice were being perfused. After all brains were extracted, hippocampi were dissected on a sterile petri dish placed atop a cold metal platform on top of ice brick to ensure brains remain cold throughout. Each brain was hemisected along the midline using a sterile scalpel. Using curved forceps with sharpened ends, bilateral hippocampi were dissected from each hemisphere of each mouse and pooled together in a 2 mL dounce with 1 ml of sample buffer (see buffer list). Cortical tissue was also dissected to be used for setting up sample gates during FACS.

Following homogenization, we adapted the isolation protocol from Bohlen et. al. 2018 ^27^. In brief, cell suspensions were filtered by passing through a 70um filter. Samples were transferred to 2 mL eppendorf tubes coated with 10% sterile filtered FBS in PBS (to prevent cell adhesion on tubes) and centrifuged. Pellets were suspended in 1.8 mL myelin removal buffer. Myelin removal beads were briefly vortexed. 200 µL of myelin removal beads were added to each sample and incubated over ice for 15 minutes with gentle flicking every 5 minutes to mix settled beads. The reaction was stopped after incubation period by diluting with 2 mL of myelin buffer per sample. Samples were transferred to 2 mL Eppendorf tubes and centrifuged. Pellets were resuspended in MACS buffer (1ml buffer/pellet). After LS columns were washed twice with flow through discarded, cell suspension was applied to columns (1 tube/LS column). LS columns were washed to elute remaining cells adhering to columns. Flow through containing demyelinated cells were transferred to 2 mL eppendorf tubes and centrifuged. Pellets were resuspended in 1 mL Sterile PBS and incubated with 1 µl Live/Dead Violet per sample for 5 minutes covered from light over ice. Samples were centrifuged, resuspended in flow buffer (containing RNAsin and DNase), and transferred to 5 mL polypropylene tubes for FACS.

#### Flow cytometry

Samples were sorted on BD Influx at the Columbia Center for Translational Immunology Flow Cytometry core. Between 113,00-132,000 cells were retrieved per sort sample. Samples were of high viability and yield. Gates were established as illustrated in Figure 1B. Cells were first selected by size and granularity (FSC and SSC, respectively). Next cells were gated to exclude doublets. Subsequently, cells were gated for viability and finally gated for td-Tomato expression. TdTomato+ cells were validated using cd11b (1:100 BD biosciences 557396) and cd45 (1:50 BD Biosciences 59864) and Tmem119 (1:100 abcam ab225495).

**Figure 1:**
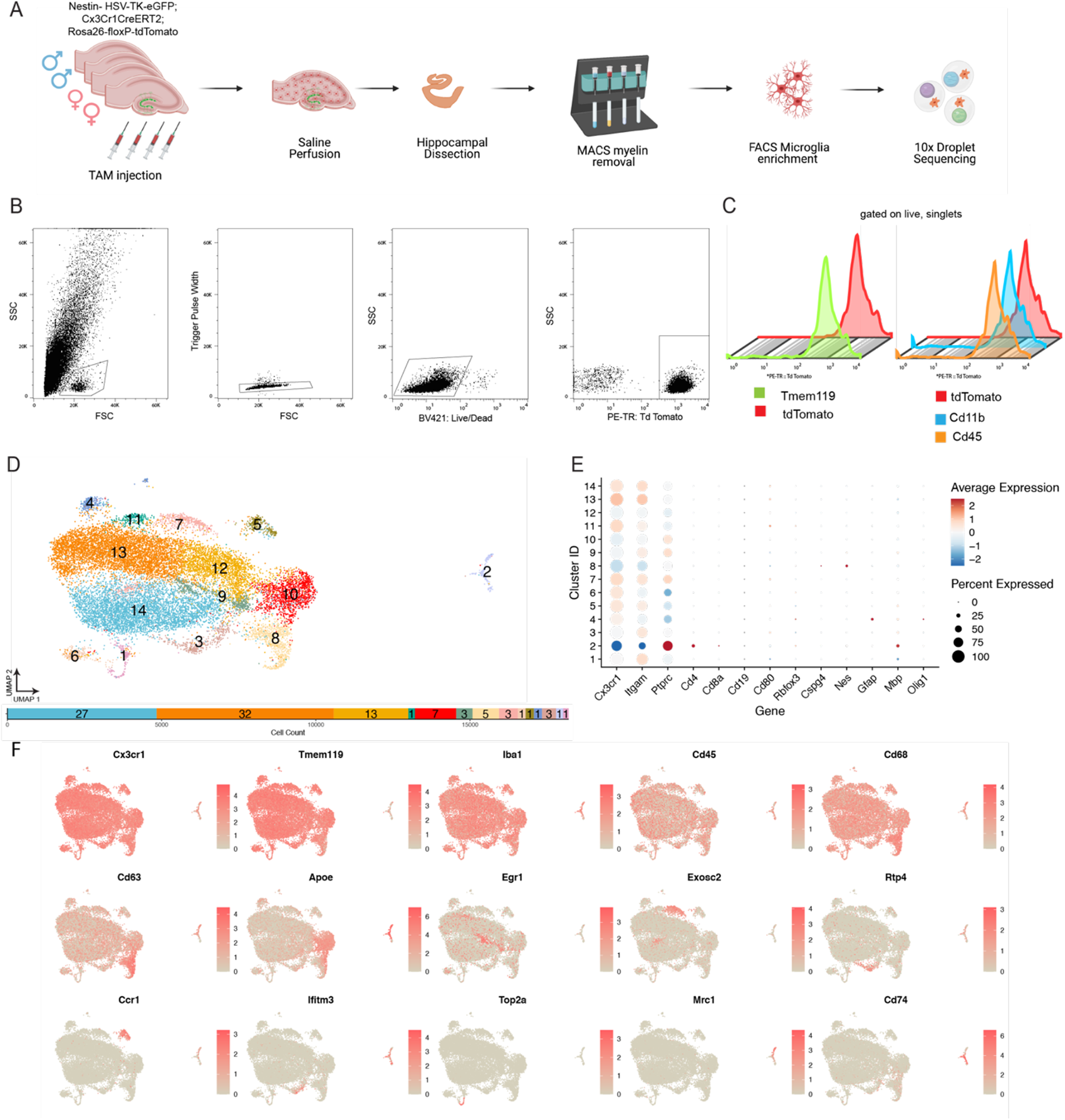
Transcriptomic Heterogeneity of myeloid population in the hippocampus. (A) Schematic illustrating experimental workflow and double reporter mouse model used in single cell sequencing experiments. (B) Fluorescent Activated Cell Sorting (FACS) gating scheme to isolate tdTomato+ cells. (C) Flow cytometry analysis of cells expressing Tdtomato with Tmem119 antibody(left) or CD11b/CD45 (right). (D) Top: UMAP Plot showing dimensionality reduction, colored by Seurat clusters. Each dot represents a cell. Bottom: Bar Plot showing distribution of cells across clusters to show cell count in each cluster (right to left starting with cluster 1). Percentage of cells in each cluster labeled within bar. (E) Dot Plot displaying expression of marker genes for cell types found in the brain across clusters. Each row represents a cluster while each column represents the level of expression of a selected gene marker. The fraction of cells in a given cluster is represented by the size of the dot. The color of the dot represents the average expression of the cells within a given cluster. (F)UMAP projection displaying scaled expression of previously identified gene markers to investigate heterogeneity in hippocampal myeloid cells.

#### Sequencing

##### Single-cell 3′ library construction and Library pooling

After samples were sorted, suspensions were spun down and resuspended at a final concentration of 1000 cells /µL with between 90,000-13,000 cells. Cell viability was confirmed with trypan blue after resuspension. Between roughly 30,000 – 40,000 cells were loaded onto 10x Chromium Controller. Libraries were constructed as per manufacturer’s (10x Genomics) instructions. Samples were sequenced at the Columbia Sulzberger genomics core. Chemistry Single Cell 3’ v3. Cell ranger v 3.0.2 for Run 1, 3.1 for Run 2 and 3. Live cell suspensions were loaded onto GEM droplets. Pooled 3’-end libraries were sequenced on NovaSeq 6000.

#### Analysis

##### Single-cell RNA-seq Preprocessing and Alignment

Read alignment to reference transcriptome mm10 was performed using Cell Ranger v 3.0. At least 94% of reads were mapped to genome. Cells were filtered using default parameters for UMI counts. Counts matrices were generated and further preprocessed as outlined below.

##### Normalization and Integration

After preprocessing of the transcriptomic data using CellRanger 3.0.2 (Run 1) and 3.1.0 (Runs 2 and 3) using CellRanger default parameters, count matrices were exported to Seurat version 4.0.0 to create a list of Seurat objects for each sample. To attain this, the raw gene-UMI matrix from each sample was converted to corresponding Seurat object in R 4. 0.3 using Read10X function from Seurat package and a list of all Seurat objects was created. To avoid confounding effects due to low quality cells, we used a standard criterion of excluding cells with higher than 20% of total UMIs from mitochondrial genes before proceeding further for normalization and integration (Supplementary table 1). To remove the potential influence of technical effects in the analyses, we normalized the raw data using function SCTransform separately for each dataset within the Seurat pipeline; this uses Pearson residuals to harmonize the data instead of regular log-normalized expression values.

Next, by running PrepSCTIntegration function that calculates all Pearson residuals, we proceeded to identify anchors and integrate the datasets using FindIntegrationAnchors And IntegrateData function respectively. The anchor as well as integration dimension was set to 30, using 3000 features total.

##### Dimensionality reduction and clustering

After count normalization and integration, we used the RunPCA function in Seurat for Principal Component Analysis and proceeded with 20 principal components for further clustering and visualization; this value was selected based on the elbow plot cutoff for significant components.

The function FindClusters from the Seurat package was used for *K*-nearest neighbor clustering with a resolution parameter of 0.5. For the visualization of integration and clustering onto a 2D space, we used uniform manifold approximation and projection (UMAP) by implementing as implemented in the Seurat functions RunUMAP, DimPlot, and UMAPPlot.

##### Identification of cluster markers via differential expression analysis

We used Seurat’s FindAllMarkers function, which implements the non-parametric Wilcoxon rank sum test, to inspect differentially expressed genes by comparing a single cluster of interest with all others. We only tested the genes that were observed to be positively (up) regulated in minimum fraction of at least 70% cells in the cluster of interested showed at least ∼1.5(logfc=0.5) fold change between cluster of interest and all other groups, with an FDR-adjusted p-value <0.01.

##### Gene enrichment analysis of differentially expressed genes

We used TopGO to find enriched Gene Ontology (GO) terms for high variance genes obtained in differential expression analysis. We ran TopGO with the Kolmogorov-Smirnov test and considered terms with a statistical significance of <0.05 (adjusted p-value).

##### Comparison of novel SGZ localized cluster 8 and homeostatic clusters (13 and 1. 14) with Keren-Shaul et al. Disease-Associated Microglia data set

We compared the transcriptomic profiles of SGZ (cluster 8) and homeostatic clusters (cluster 13 and 14) to microglia subtypes signatures found in Keren-Shaul et al’s study (GEO: GSE98969). The primary goal of the integration was to compare the unique signature found in SGZ-enriched cluster with the <u>disease-associated microglia</u> (DAM) signature identified by Keren-Shaul et al.

We used Harmony to integrate our data with the Keren-Shaul et al. data by passing a merged Seurat object consisting of all selected microglial cells from both datasets and following the standard pipeline through PCA. We then ran the *RunHarmony* function on the normalized data, where we ran 10 rounds of iteration with default values of theta and lambda to achieve the corrected harmony coordinates. We used the first 10 dimensions from the Harmony integration to generate the UMAP and run nearest neighbor analyses, using a resolution of 0.7 with the harmony embeddings rather than PCs. We investigated the expression of genes known to be downregulated in DAM profile to map the cluster associated with unique signature found in SGZ-enriched cluster in the integrated dataset.

##### Data and Code availability

Single cell sequencing analysis was done primarily using Seurat version 4.0.0. Scripts and code are available on GitHub at https://github.com/sanachintamen/HC_myeloid. Raw BAM files and Cell Ranger processed gene expression matrices are available in the NCBI GEO databank with accession number GSE182289.

#### Brain Sectioning and Immunohistochemistry

Mice were perfused with PBS followed by 4% paraformaldehyde. Brains were dissected and post-fixed over night at 4%C. Free floating sections were cut with 50 µm thickness on Leica 1000S vibrating blade microtome. Sections were permeabilized in 0.3% PBST followed by 1 hour incubation with 5% Normal Donkey Serum at RT. Samples were then incubated overnight in primary antibody at the following concentrations 1:500 Iba1 (Wako 019-19741), 1:100 Cd68 (Biorad MCA1957), 1:200 Cd9 (BioLegend 124802) at 4%C. The next day, sections were washed thrice with PBST for 5 minutes at room temperature. Afterwards, they were incubated with secondary antibody staining solution containing secondary antibodies at a final concentration of 1:200 (Jackson immunoresearch). Sections were incubated in secondary antibody solution for two hours at room temperature after which they were washed thrice in PBST and twice in PBS. Sections were then mounted using LifeTechnologies Prolong mounting medium containing DAPI or NucBlue.

#### Imaging and Analysis

Images for Sholl Analysis were obtained using a Laser Scan confocal microscope (TCS SP8, Leica). Z-stack images were acquired at 0.5μm intervals. Images were projected across z-planes for representative images. Cells were traced in a semi-automated manner in 3D using Neurolucida. Dendritic complexity was measured using intersections at 10 µm radii were used to determine microglial process ramification. These intersections were summed for each cell to obtain the total number of intersections. For confocal imaging of sections from dual reporter mice, Nikon Ti Eclipse inverted with Yokogawa CSU-X1 spinning disk was used to minimize photobleaching. Images were obtained using 25x water of 40x air objectives. Images were stitched together using NIS-Elements. Epifluorescent images were acquired with a Zeiss microscope (Axio Imager M2, Zeiss) equipped with a Hamamatsu camera (Orca-R2, Hamamatsu).

## RESULTS

### Single cell sequencing of myeloid cells in the hippocampus

The brain immune compartment consists primarily of innate immune cells which include microglia (the predominant cell type) as well as other CNS-associated macrophages. To dissect cellular interactions between immune cells and neural progenitor/precursor cells in the neurogenic niche, we utilized a double reporter mouse model expressing eGFP under the control of the Nestin promoter and tdTomato conditionally active in cells expressing the fractalkine receptor (CX3CR1) and CreERT2^28^ (Supplementary Figures 2 & 3). These double reporter mice were used for both transcriptomic and histology experiments (Figure 1A). We isolated tdTomato+ myeloid lineage cells for single cell RNA-sequencing (Figure 1B). We validated the reporter line using flow cytometry analysis using antibodies common for microglia and showed near complete overlap with Cd11b^hi^/Cd45^lo^ cells as well as Tmem119^+^ cells (Figure 1C).

Single cell RNA-sequencing experiments were conducted to determine which cells compose the myeloid landscape of the hippocampus. For each experiment, four hippocampi of tdTomato positive mice (two male, two female) were pooled and enriched for tdTomato-positive myeloid lineage cells, which were then sequenced with three technical triplicates, yielding cells from the hippocampi of twelve mice in total.

The pre-processing of the data yielded 20,376 combined cells from all samples. After filtering out the cells with higher than 20% mitochondrial gene expression, the remaining 18,198 cells were included in downstream normalization, analysis, and integration (Supplemental Figure 2). We applied Canonical Component Analysis (CCA)-based integration on our integrated dataset to account for batch effects, and found that cells from different batches mixed together across all major cell types after applying CCA. Subsequently, we conducted principal component analysis for dimensionality reduction in order to cluster myeloid cells based on transcriptome profiles; this approach (see Methods) yielded 14 clusters. These are illustrated in Figure 1D and we refer to these clusters as such for downstream analysis. Cells from all clusters expressed common marker genes that were mostly specific to microglia and other macrophages in the CNS (Figure 1E, Supplemental Figures 3C & 4).

We next determined whether there were sex-specific differences in the transcriptome profiles of cells originating from male versus female samples. To do this, we plotted *Xist* expression, a gene expressed specifically by the inactivated X chromosome in female cells (Supplemental Figure 5). We found that cluster 14 is the only cluster primarily comprising female cells and that *Xist* expression separates cluster 14 from clusters 12 and 13 (Supplemental Figure 4). Based on the expression of known genes, we designated these three clusters as homeostatic clusters. We then tested whether there were significant differences in gene expression related to immune function in these putative homeostatic clusters (Table 1). We found that none of the differentially expressed genes reflect significant changes in immune function between cells of male and female origin in these homeostatic populations.

**Table 1:**
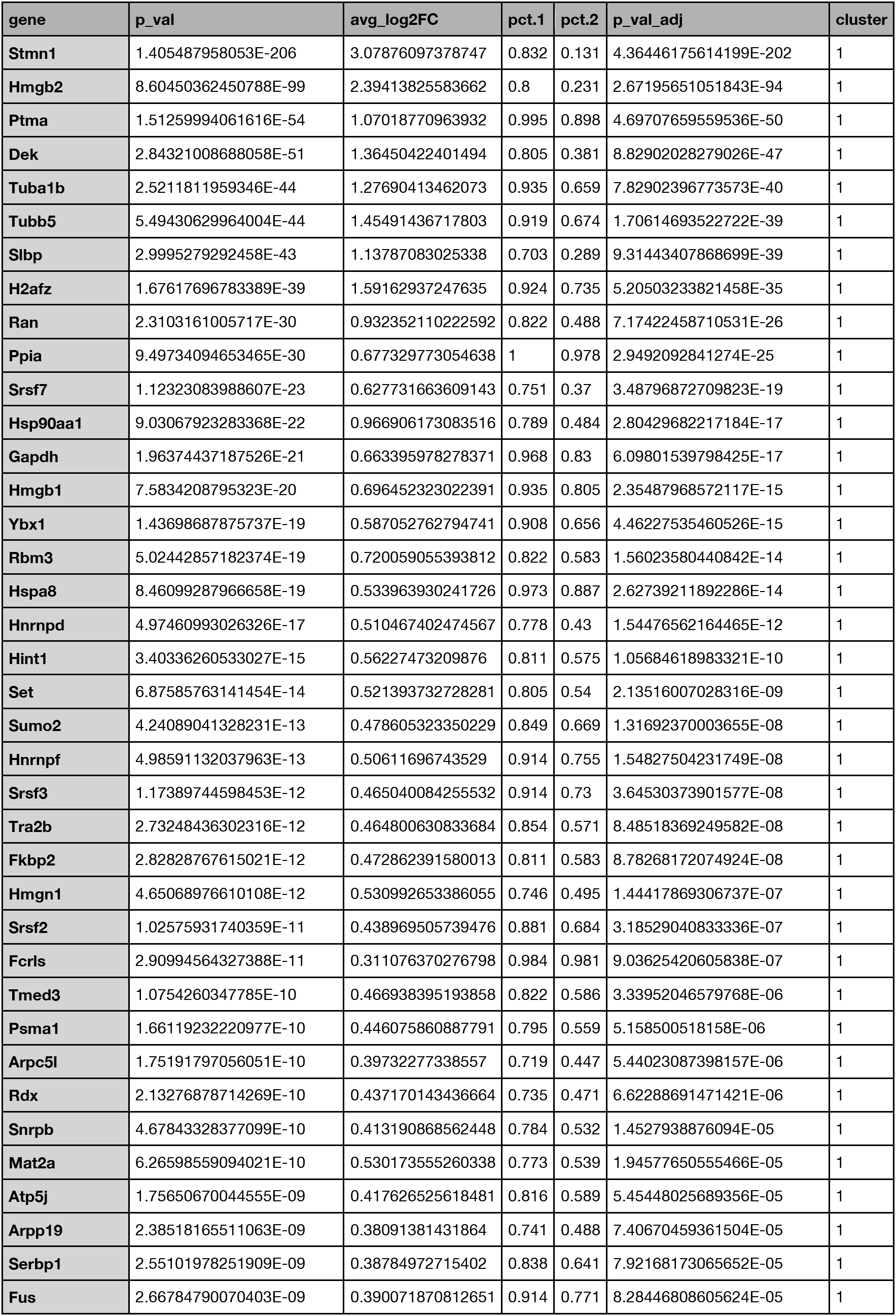
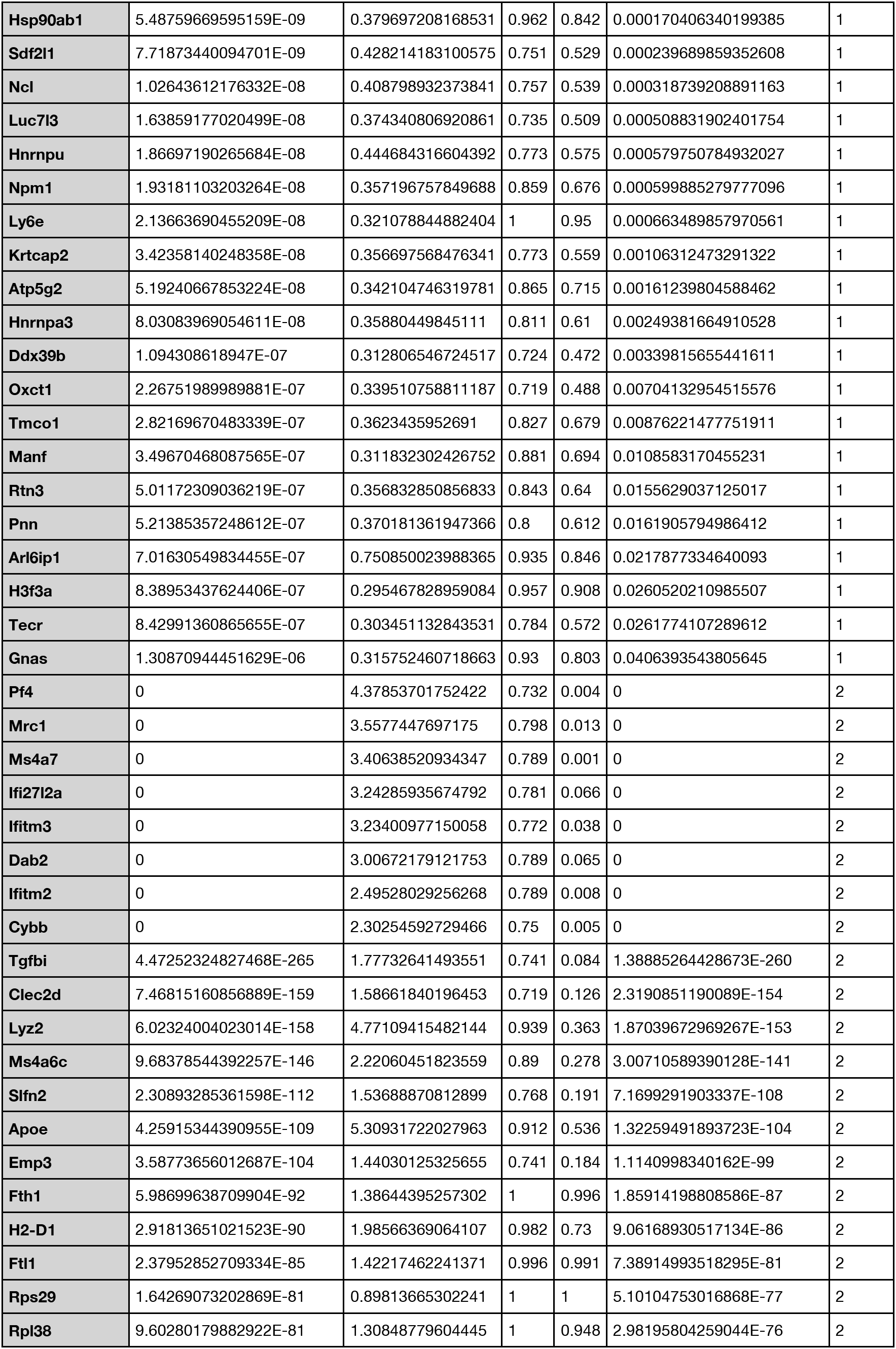
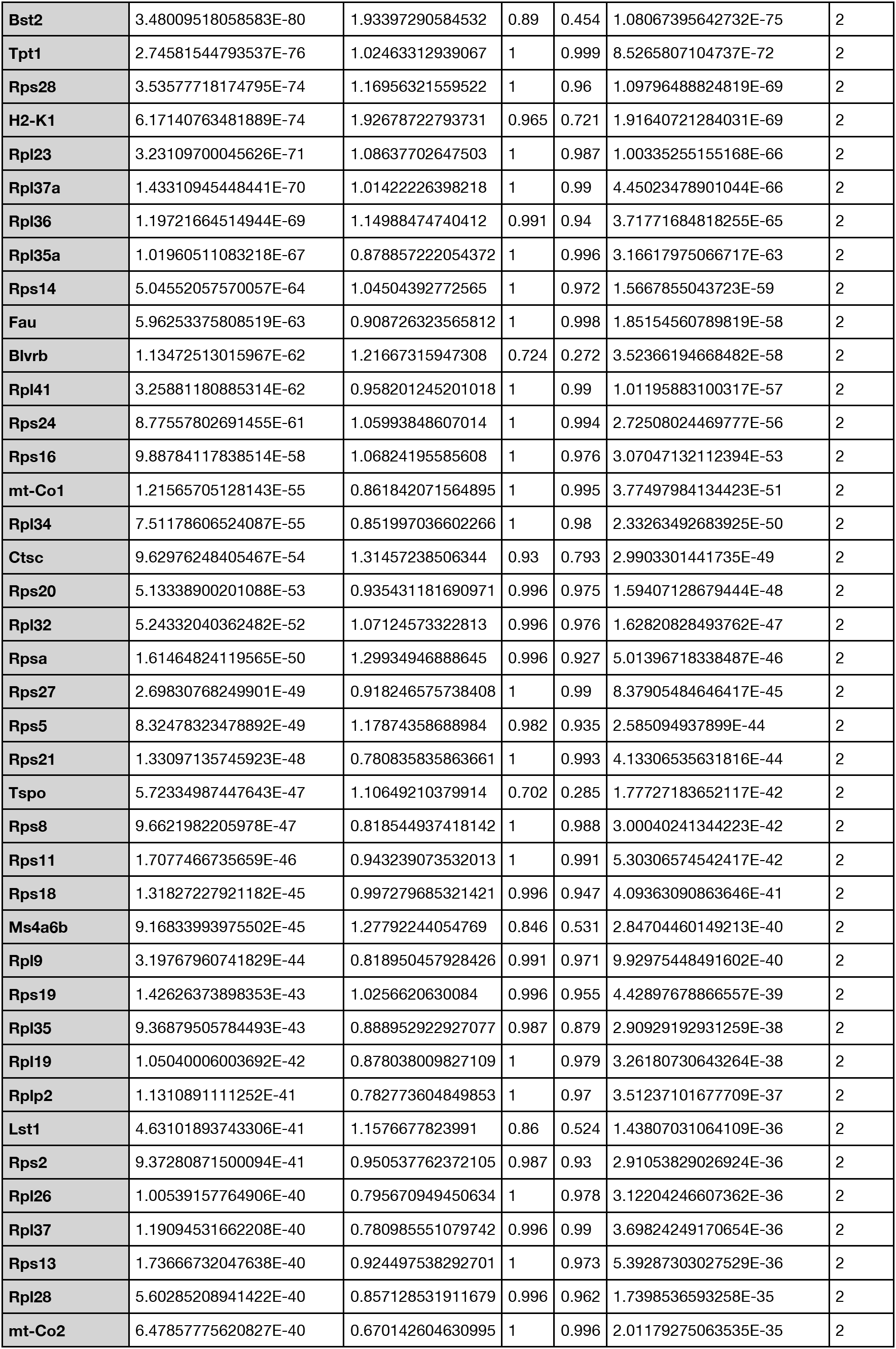
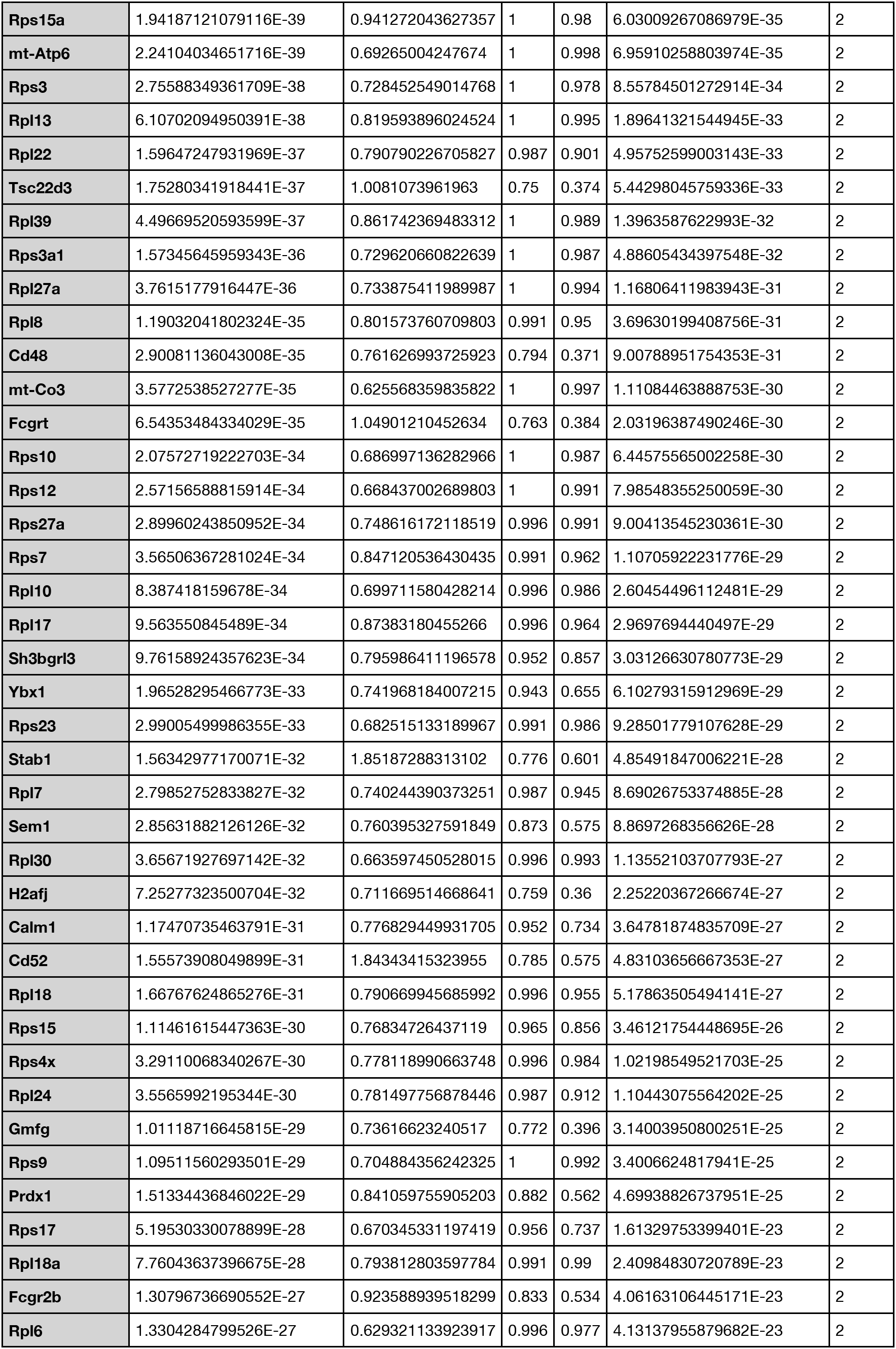
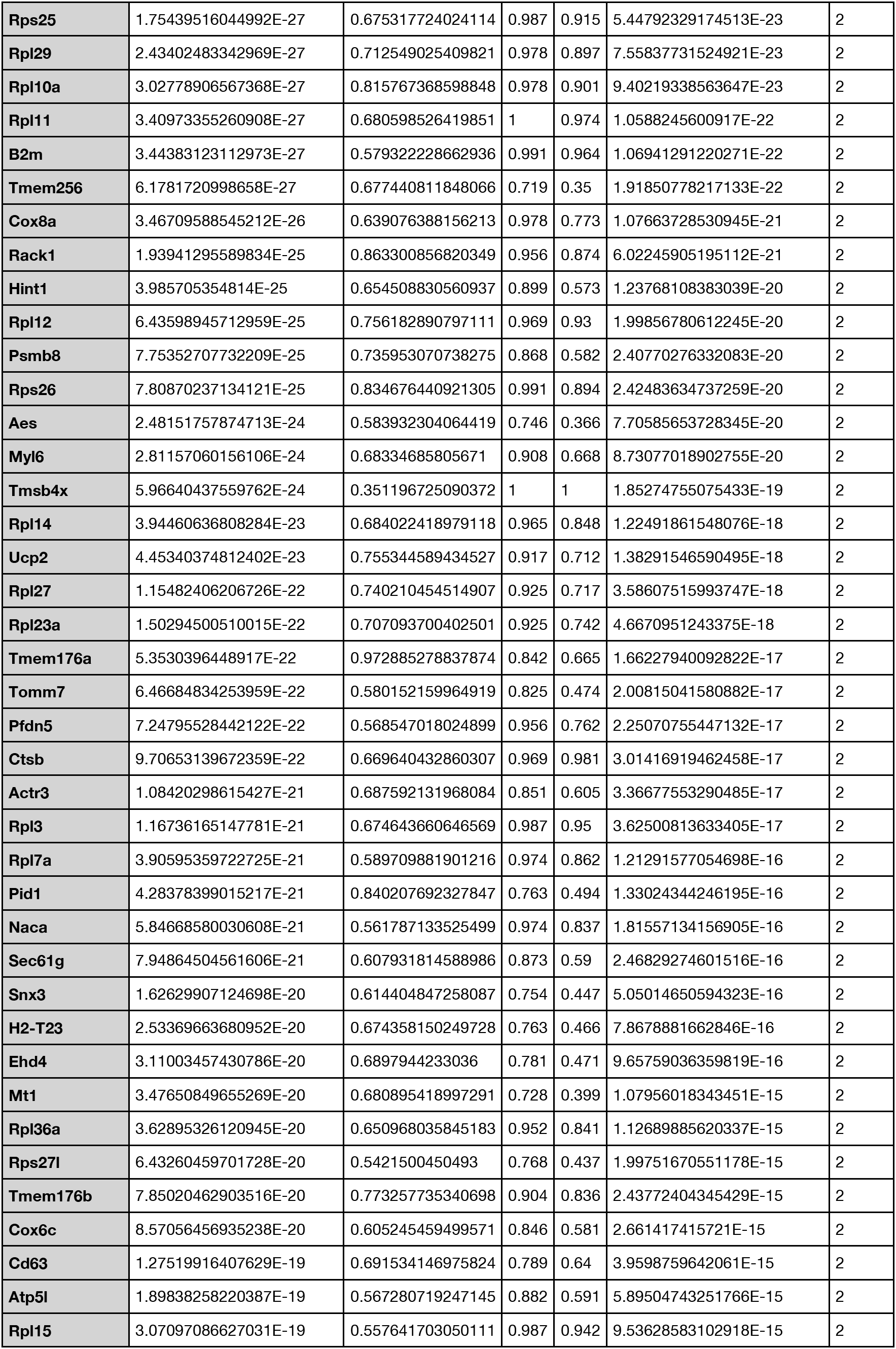
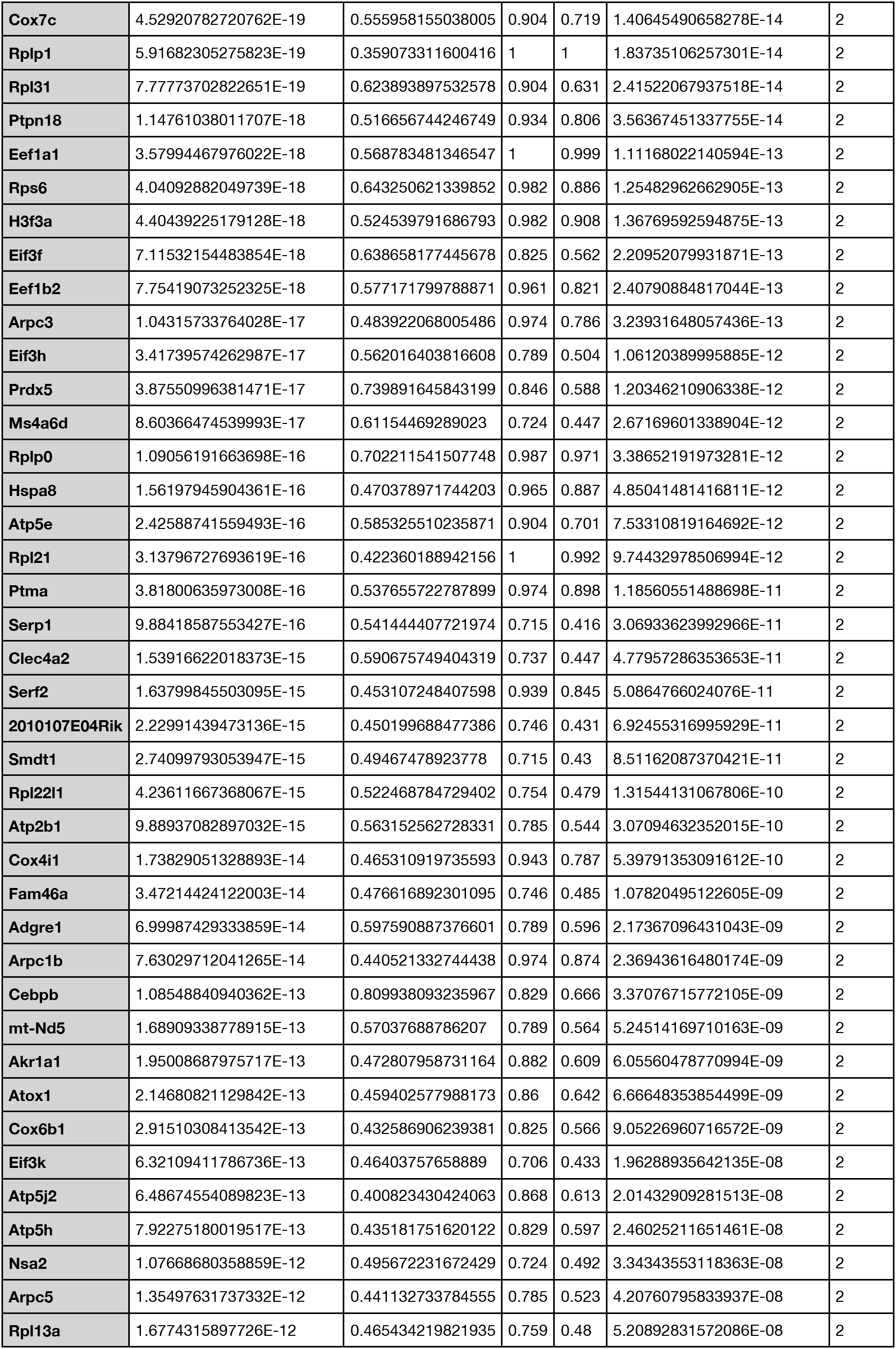
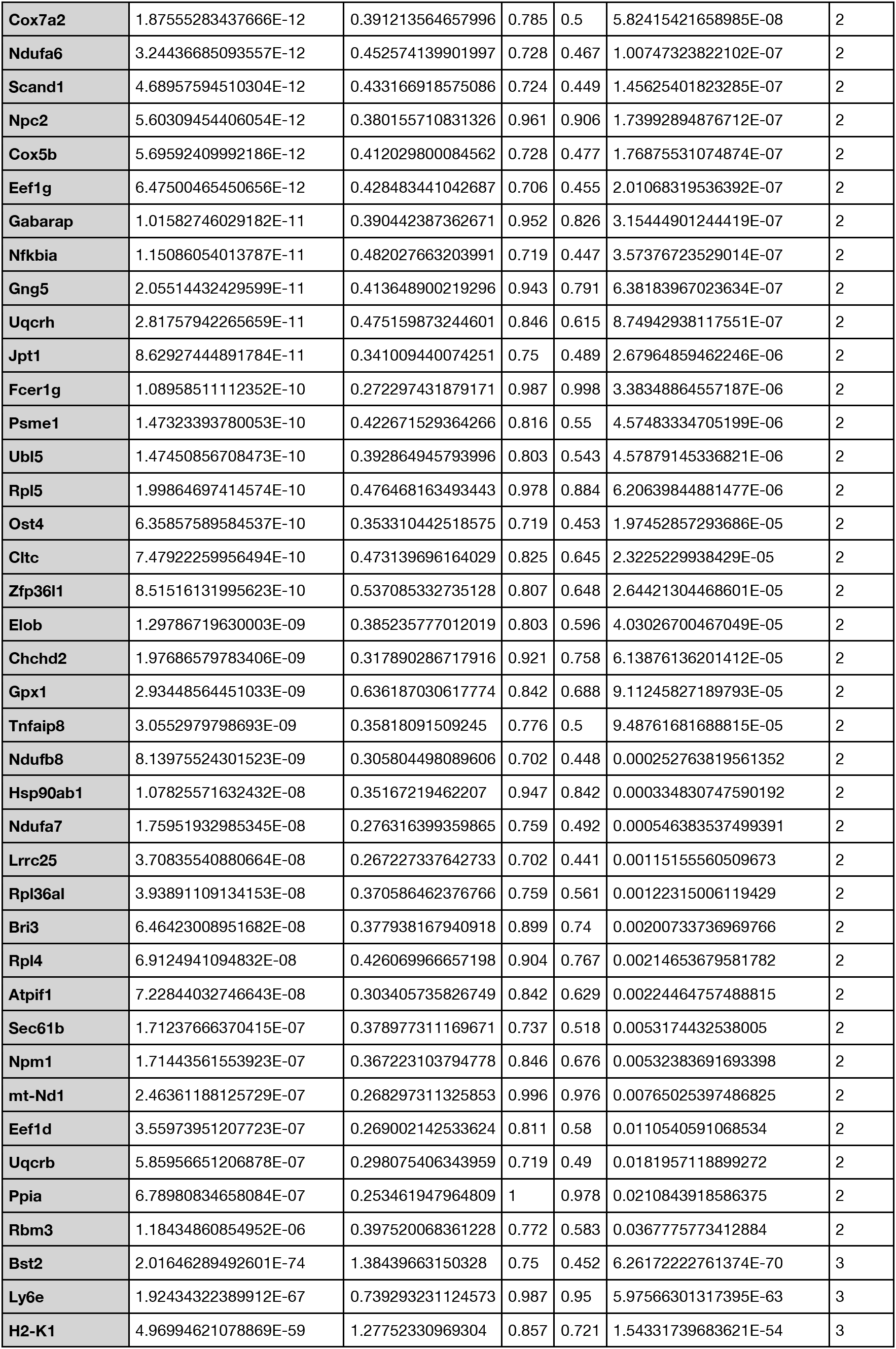
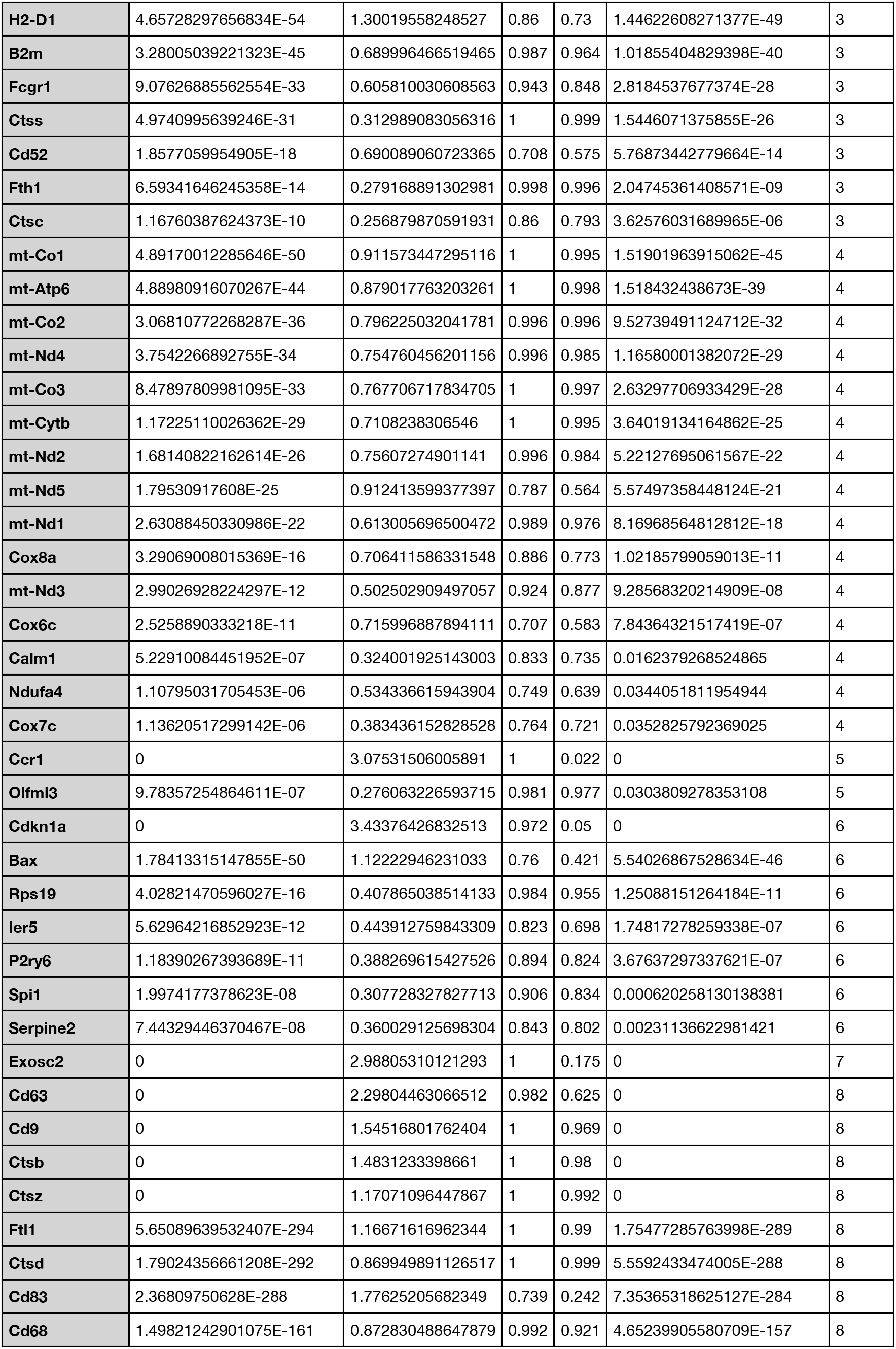
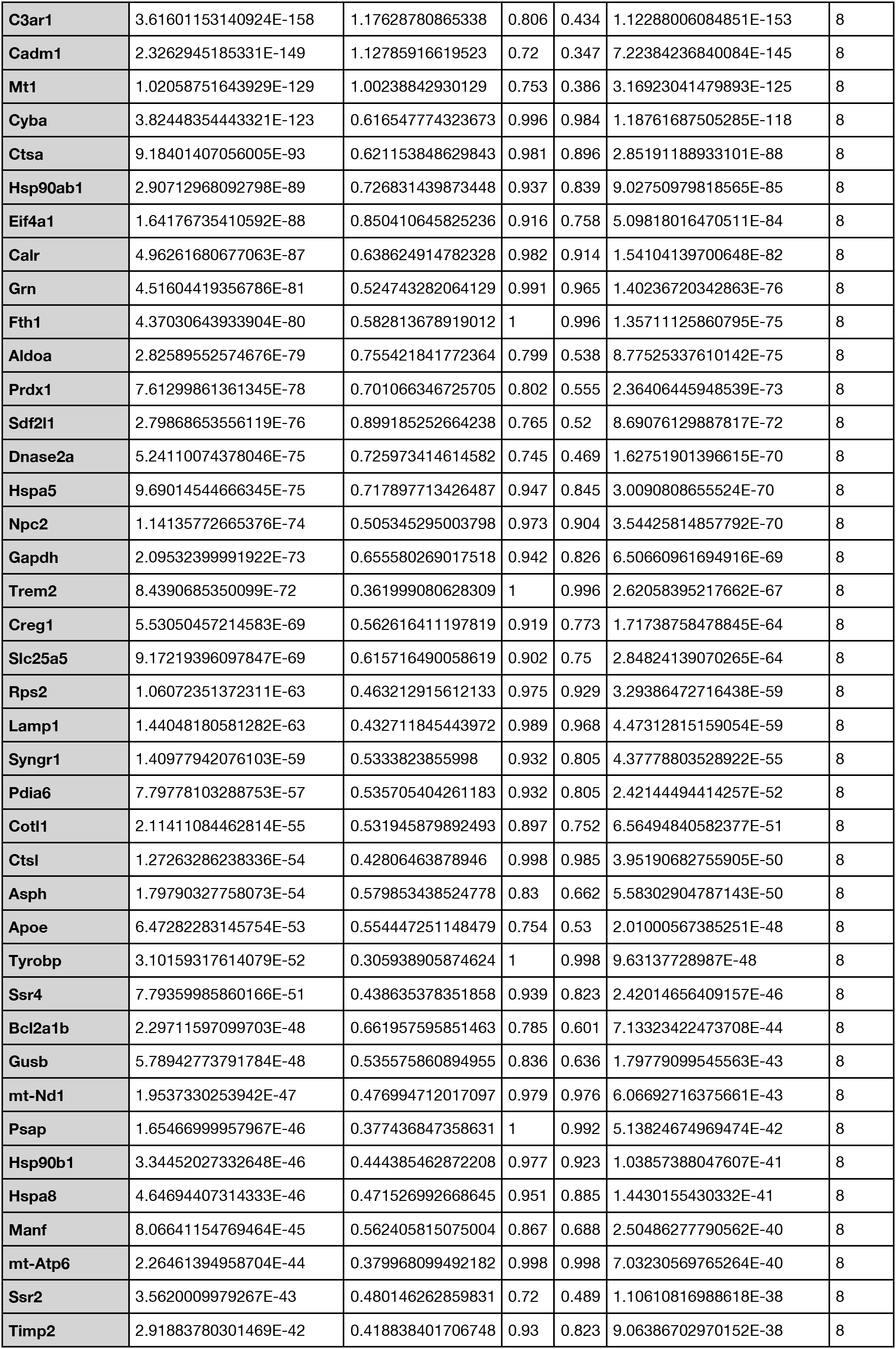
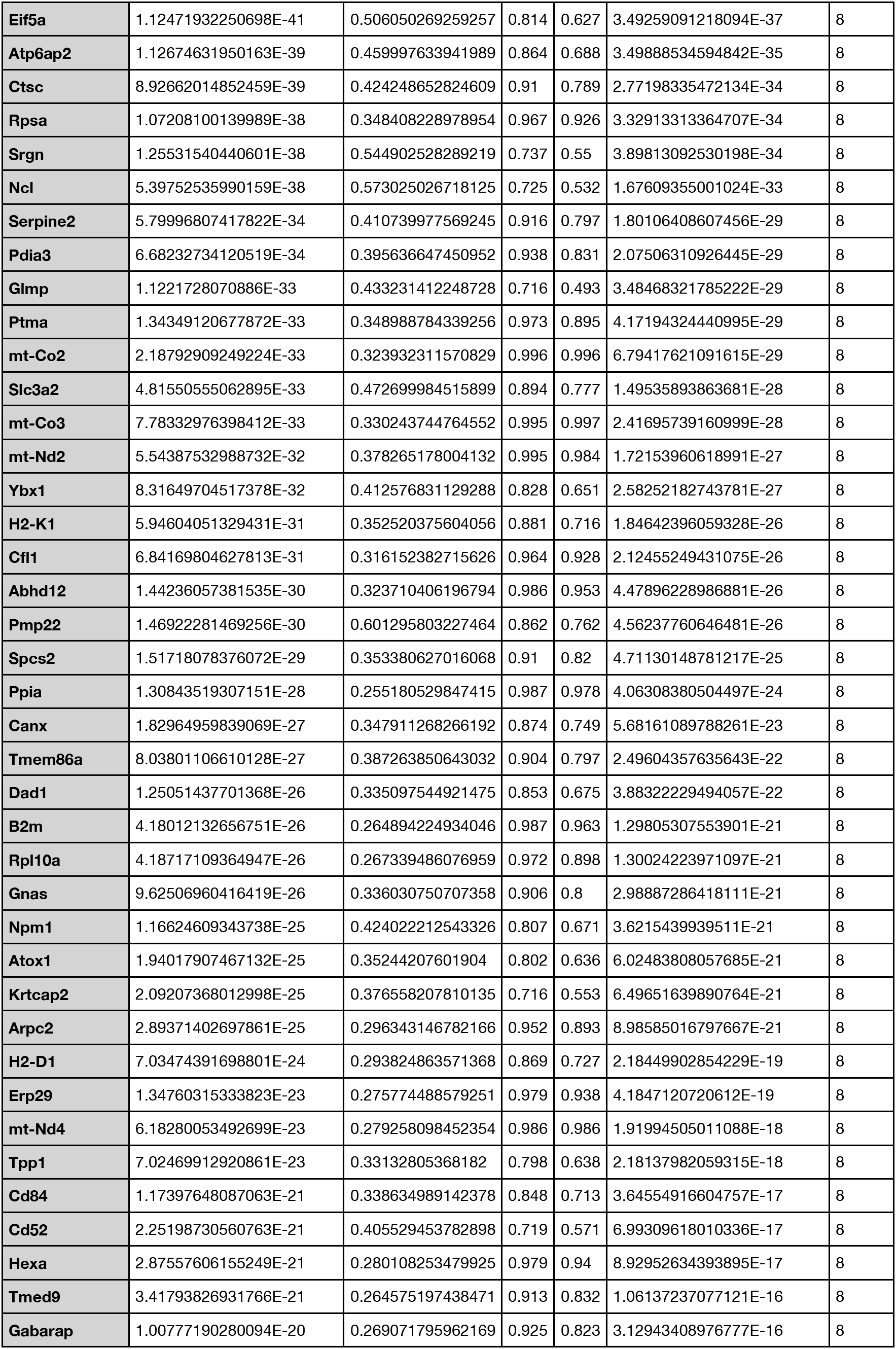
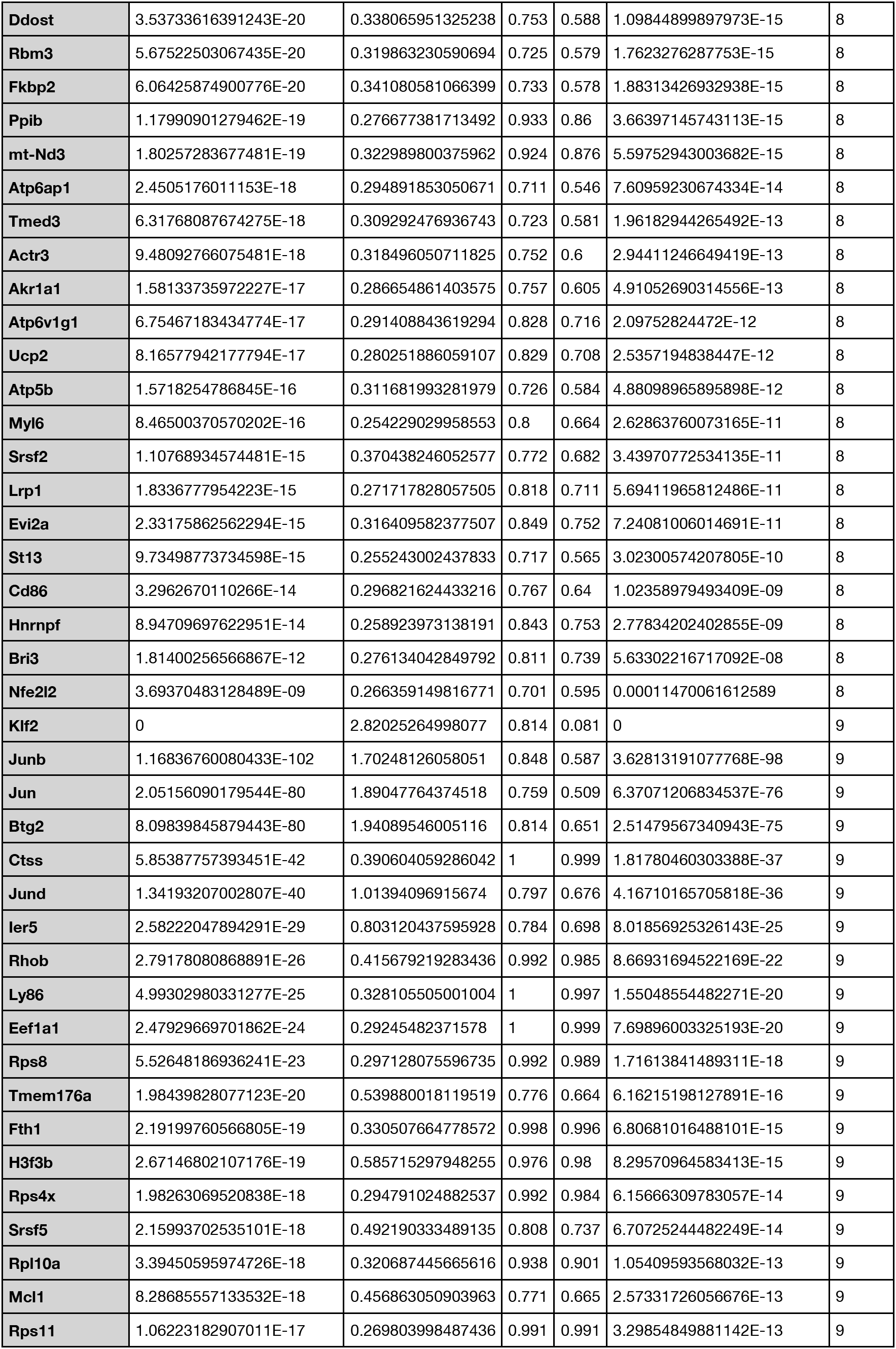
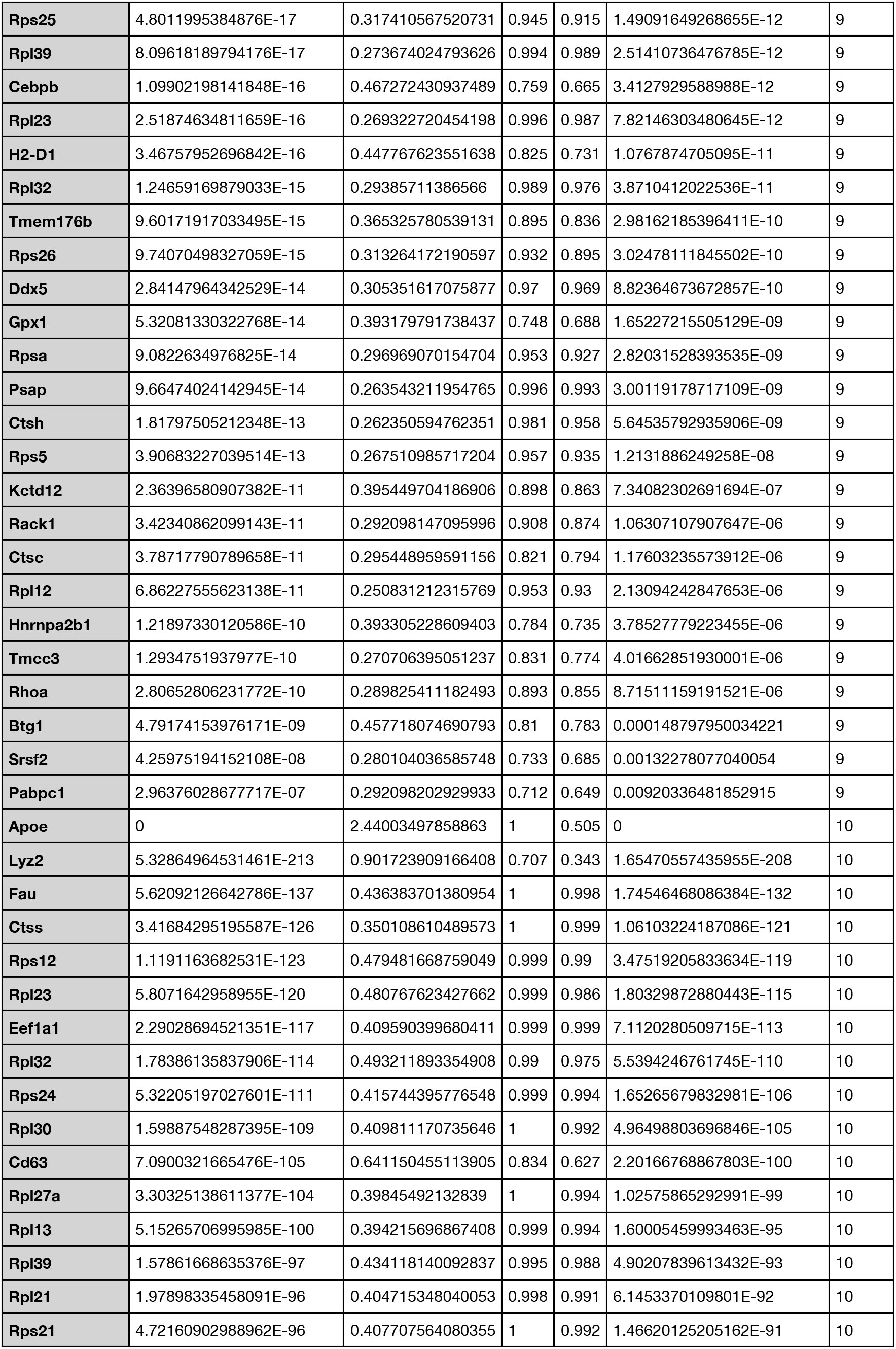
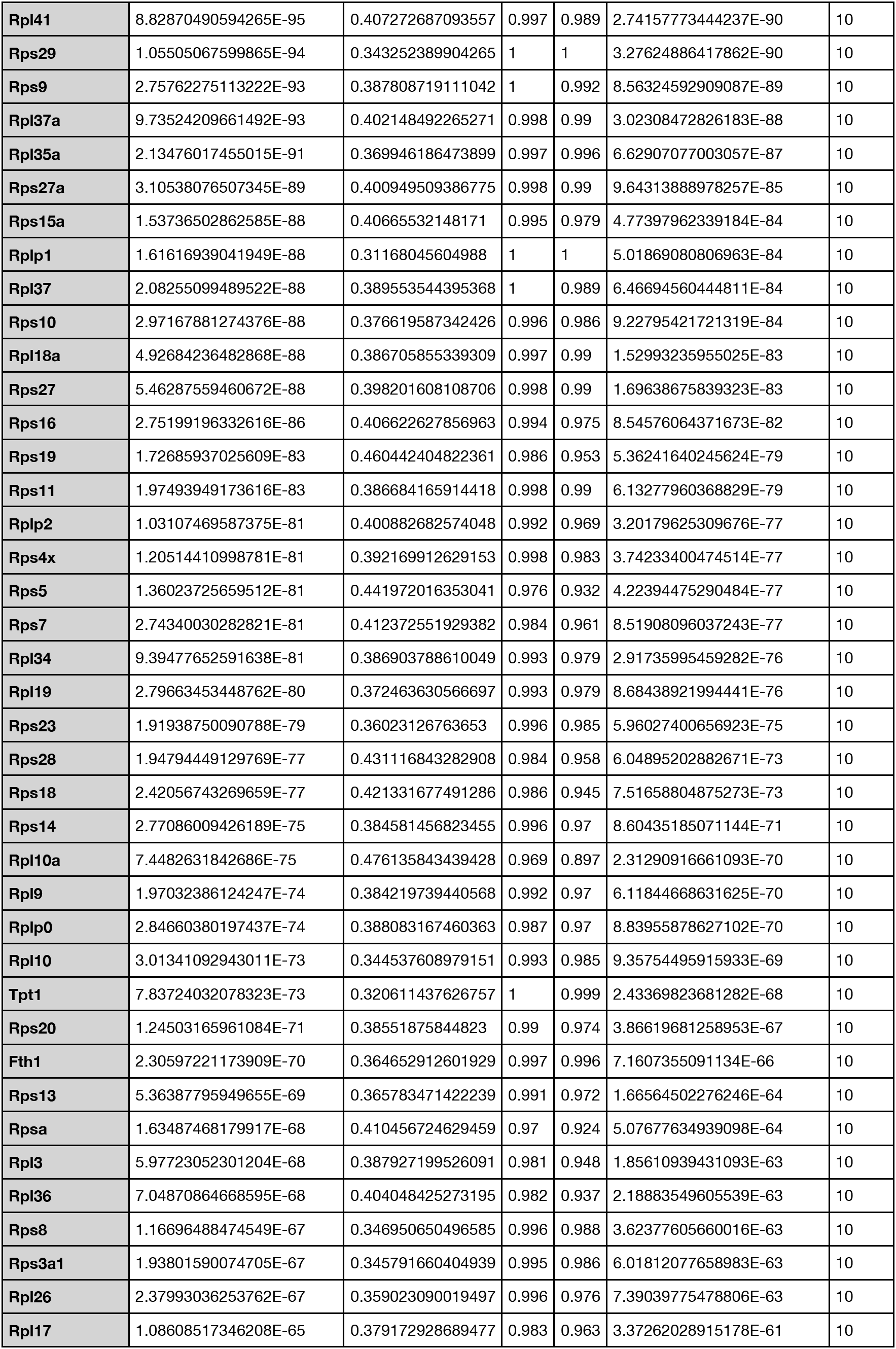
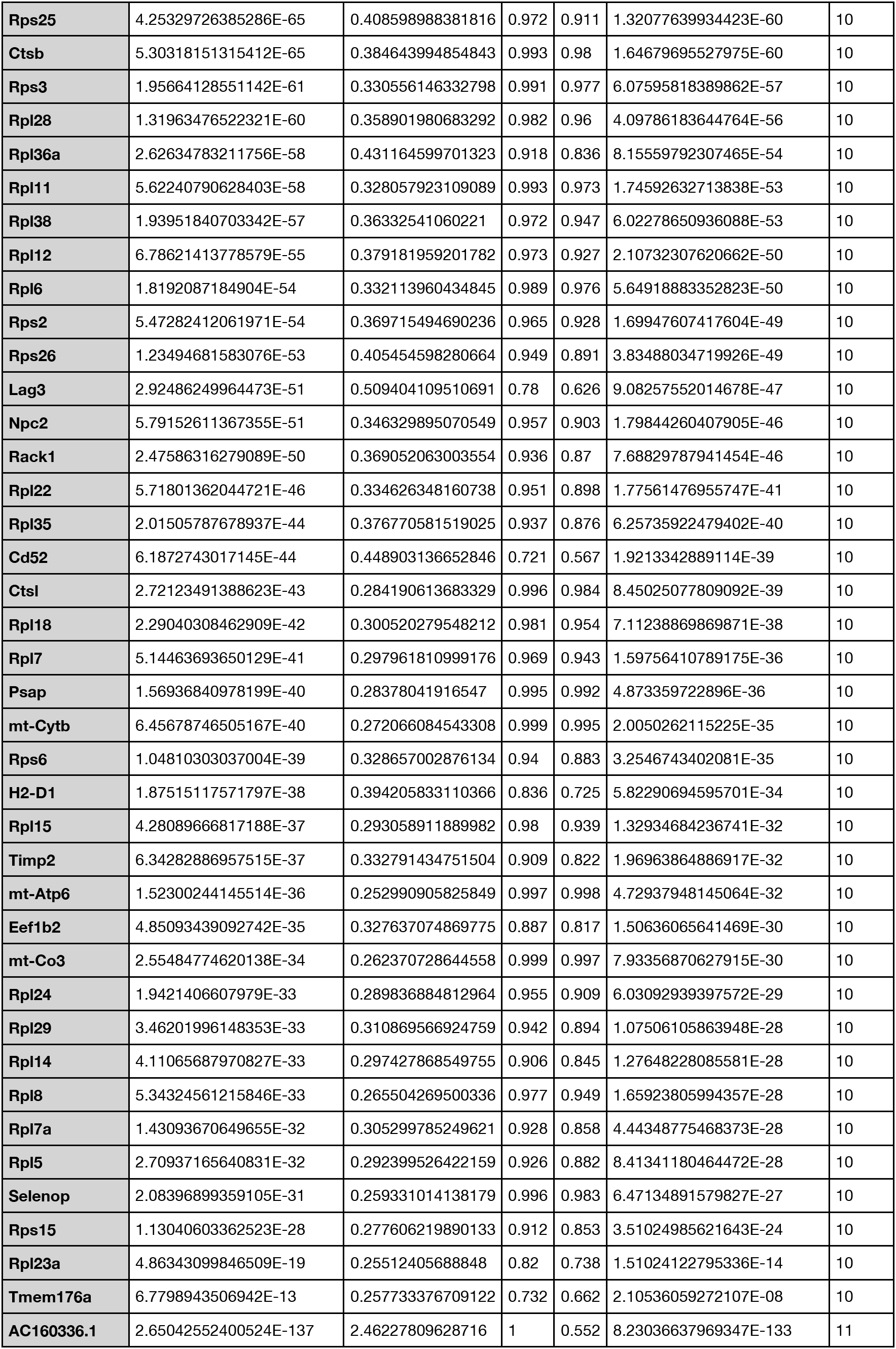
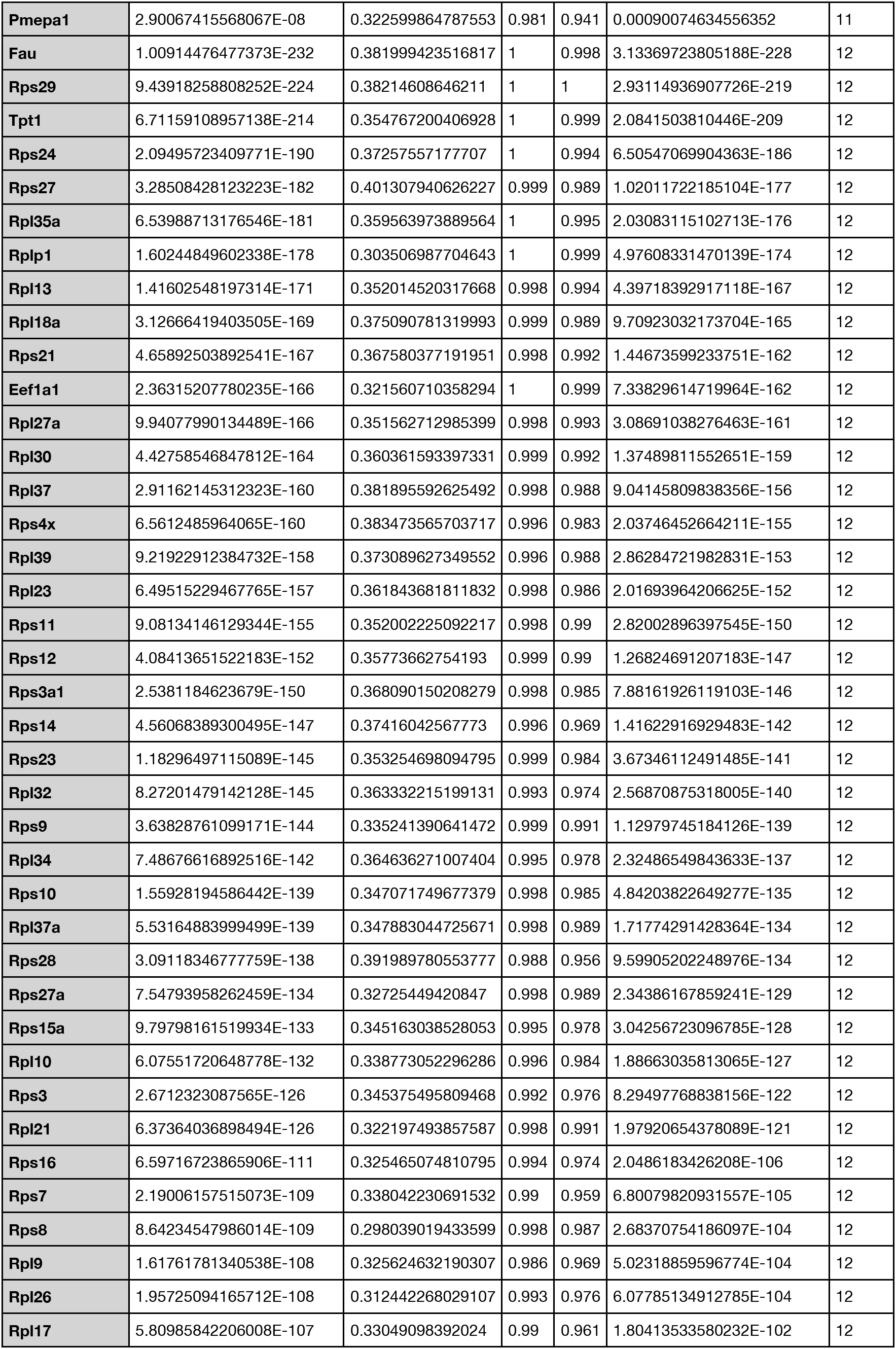
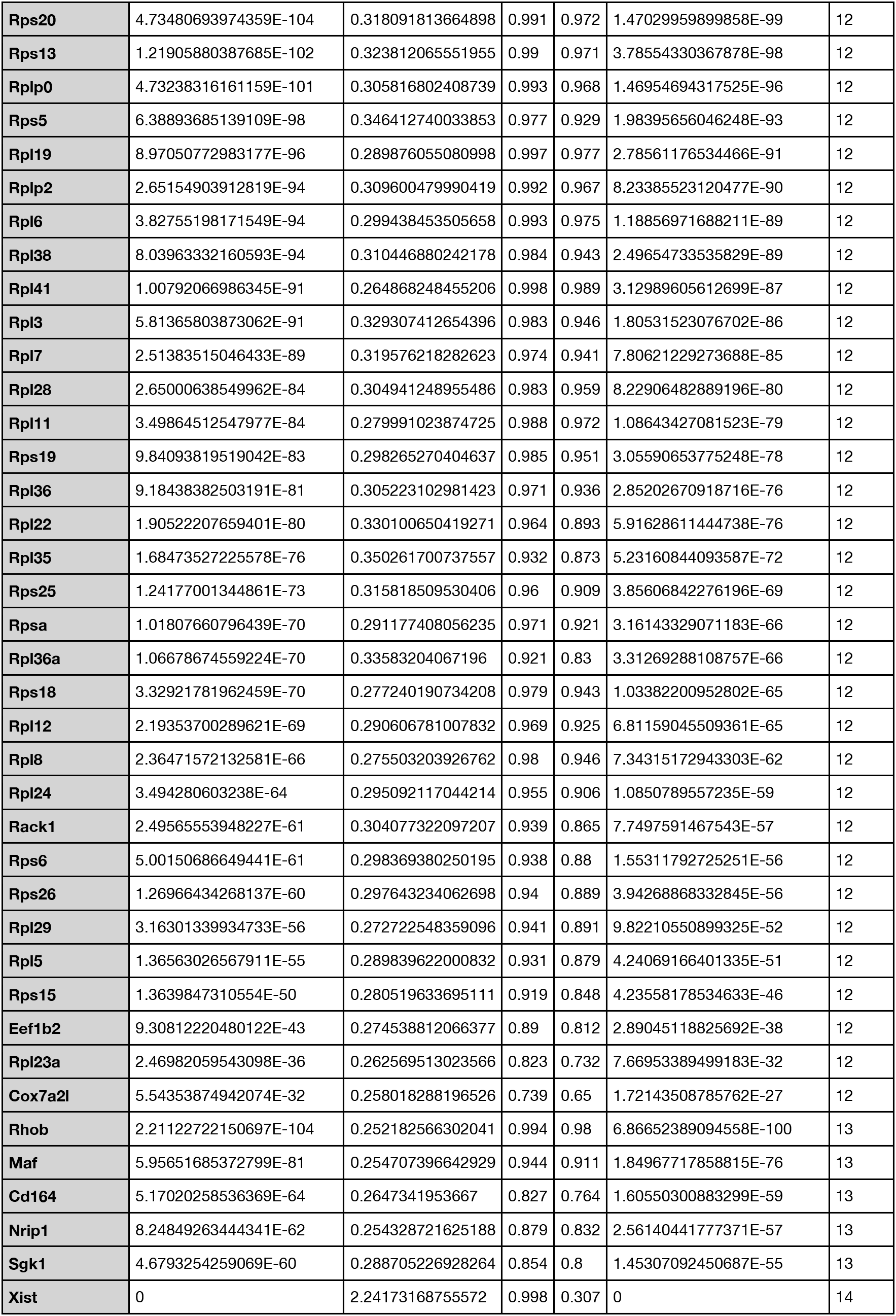
Upregulated in each cluster.

**Table 2:**
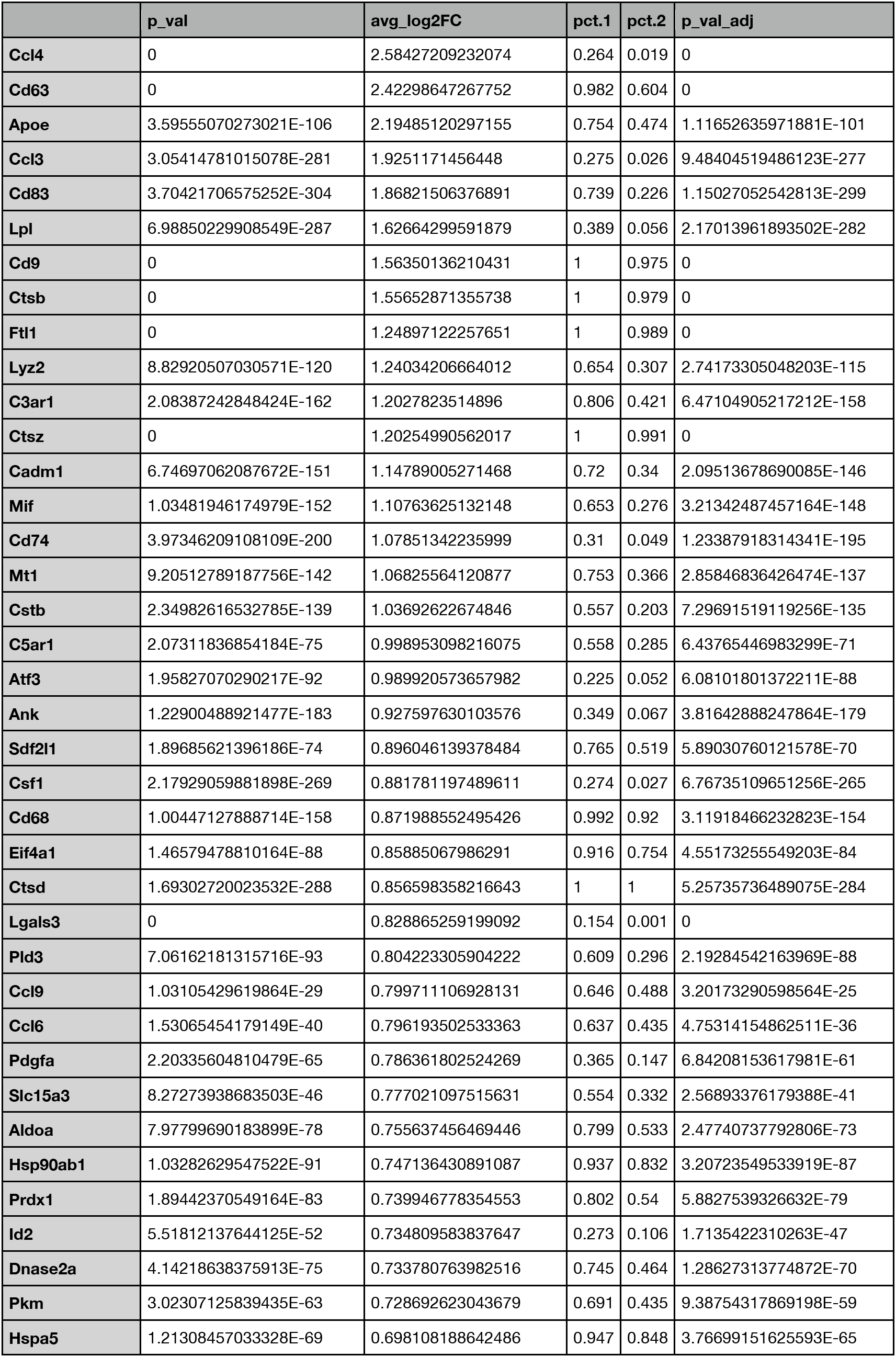
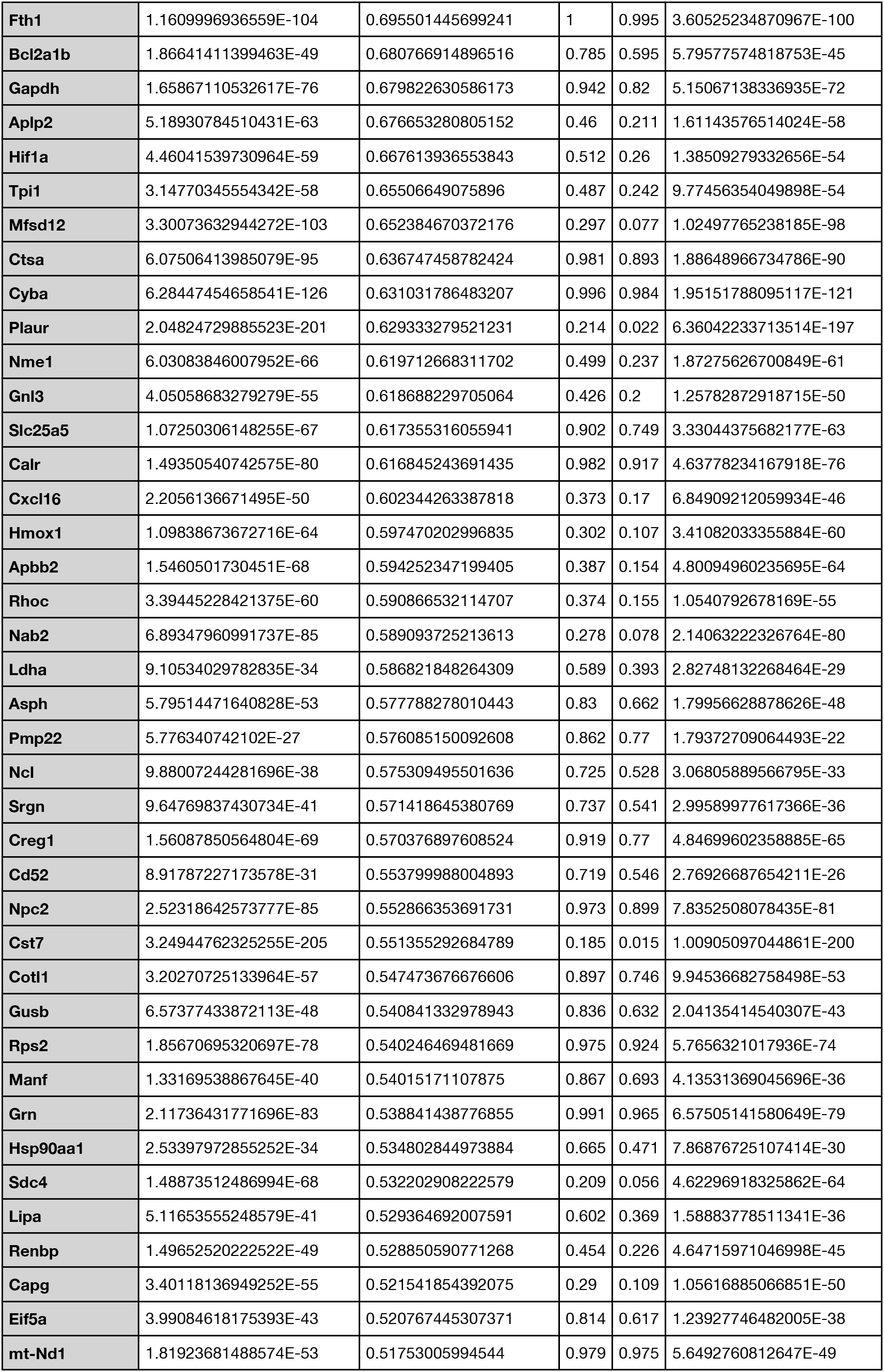
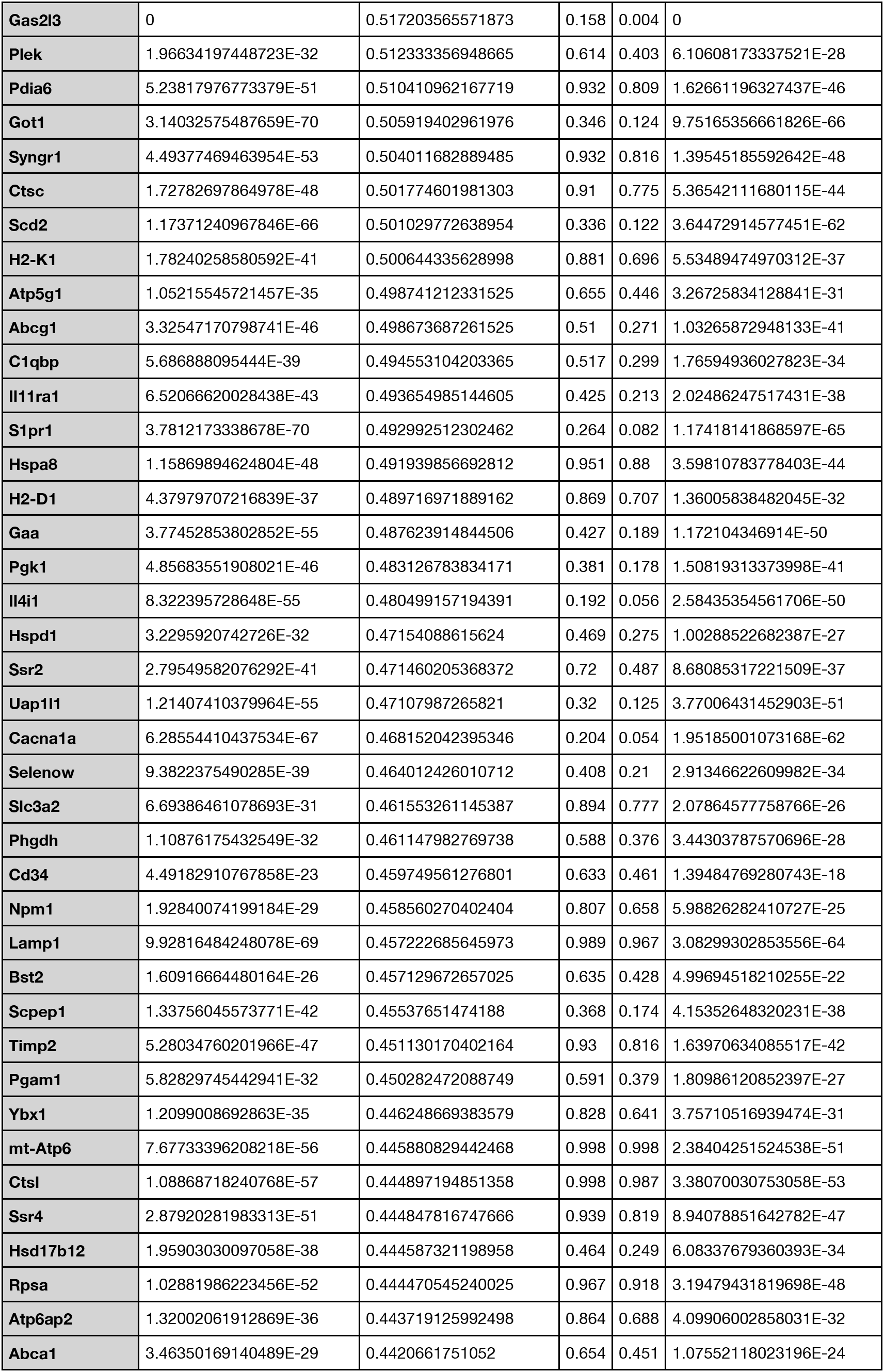
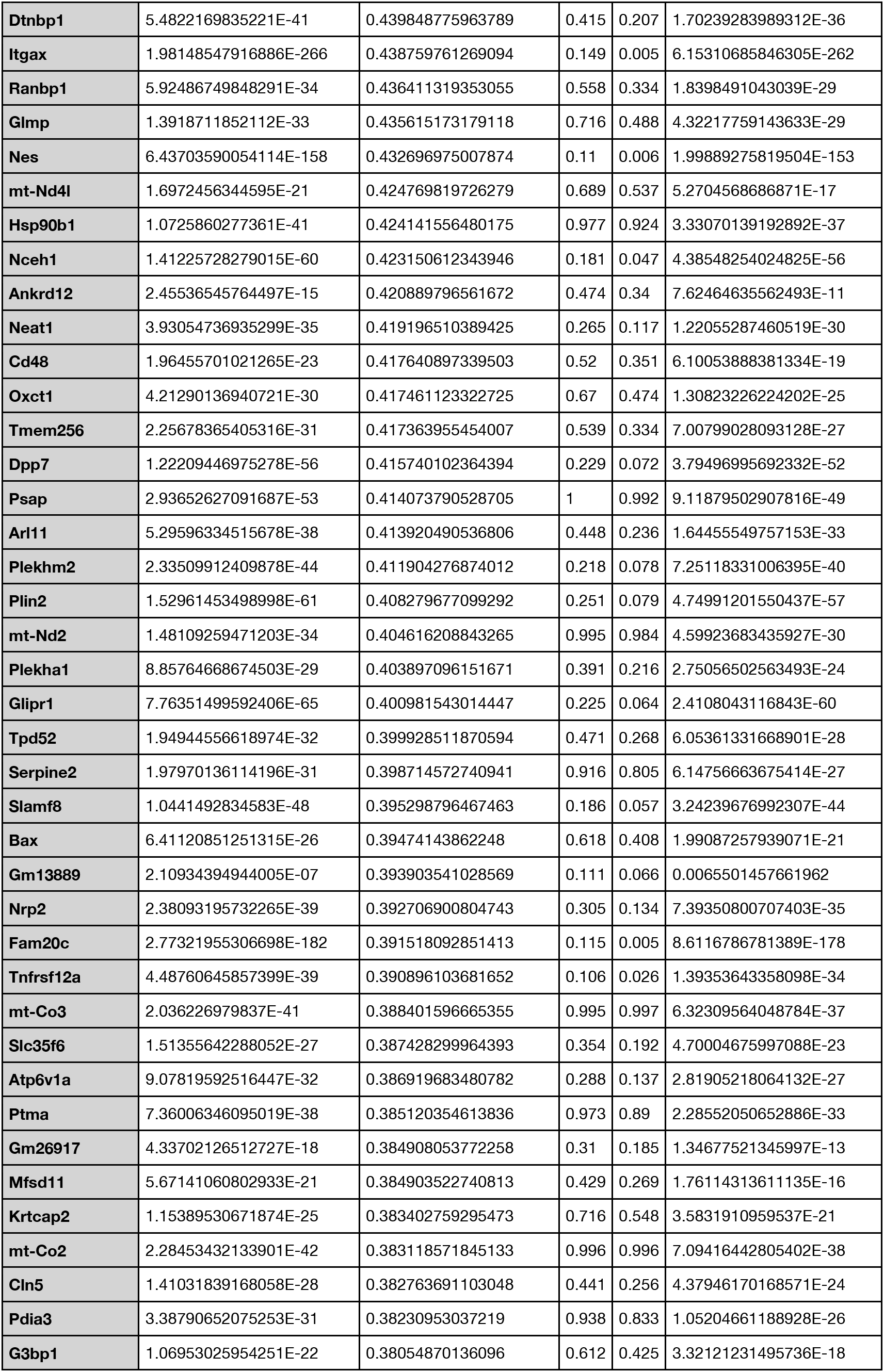
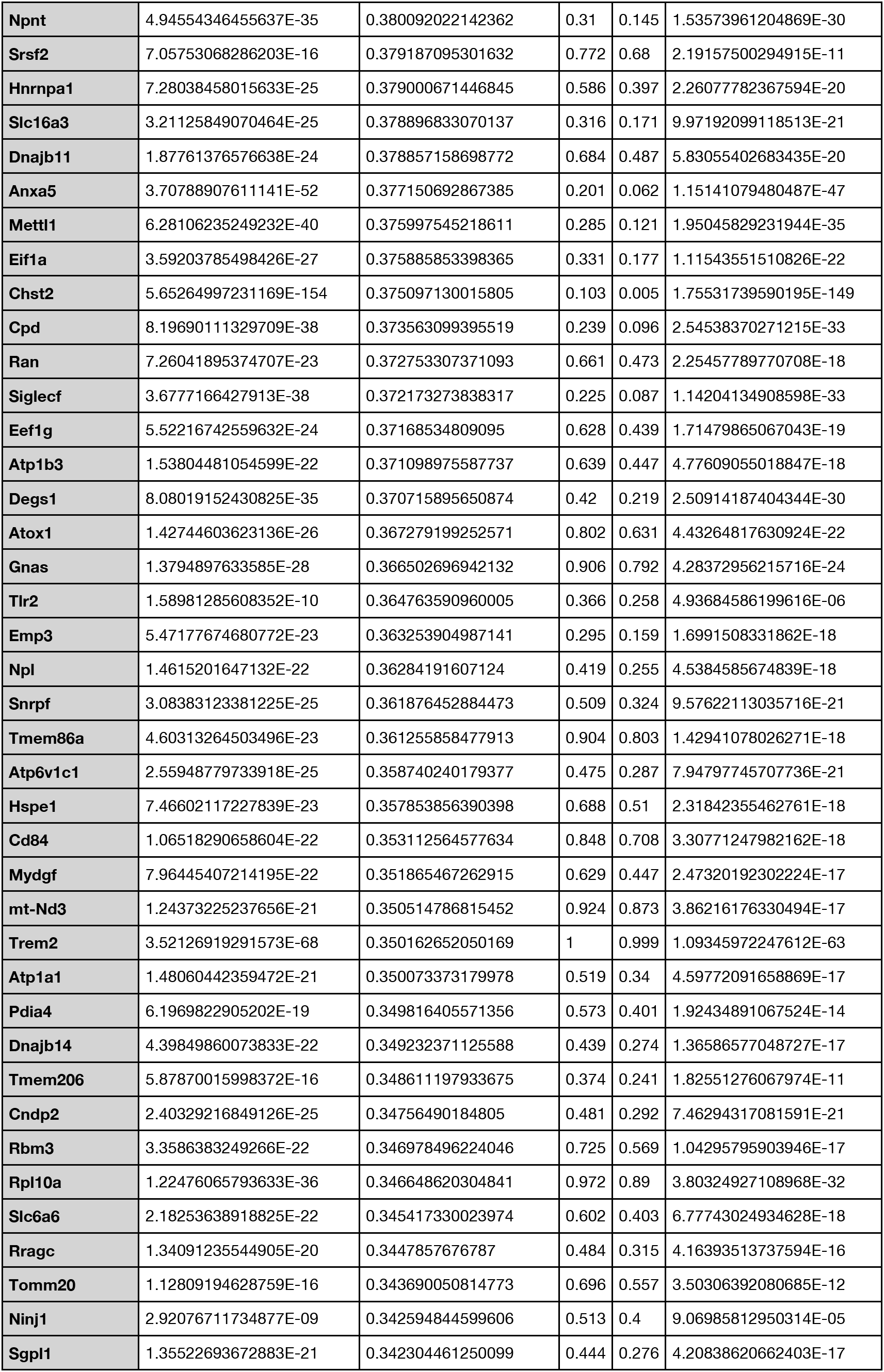
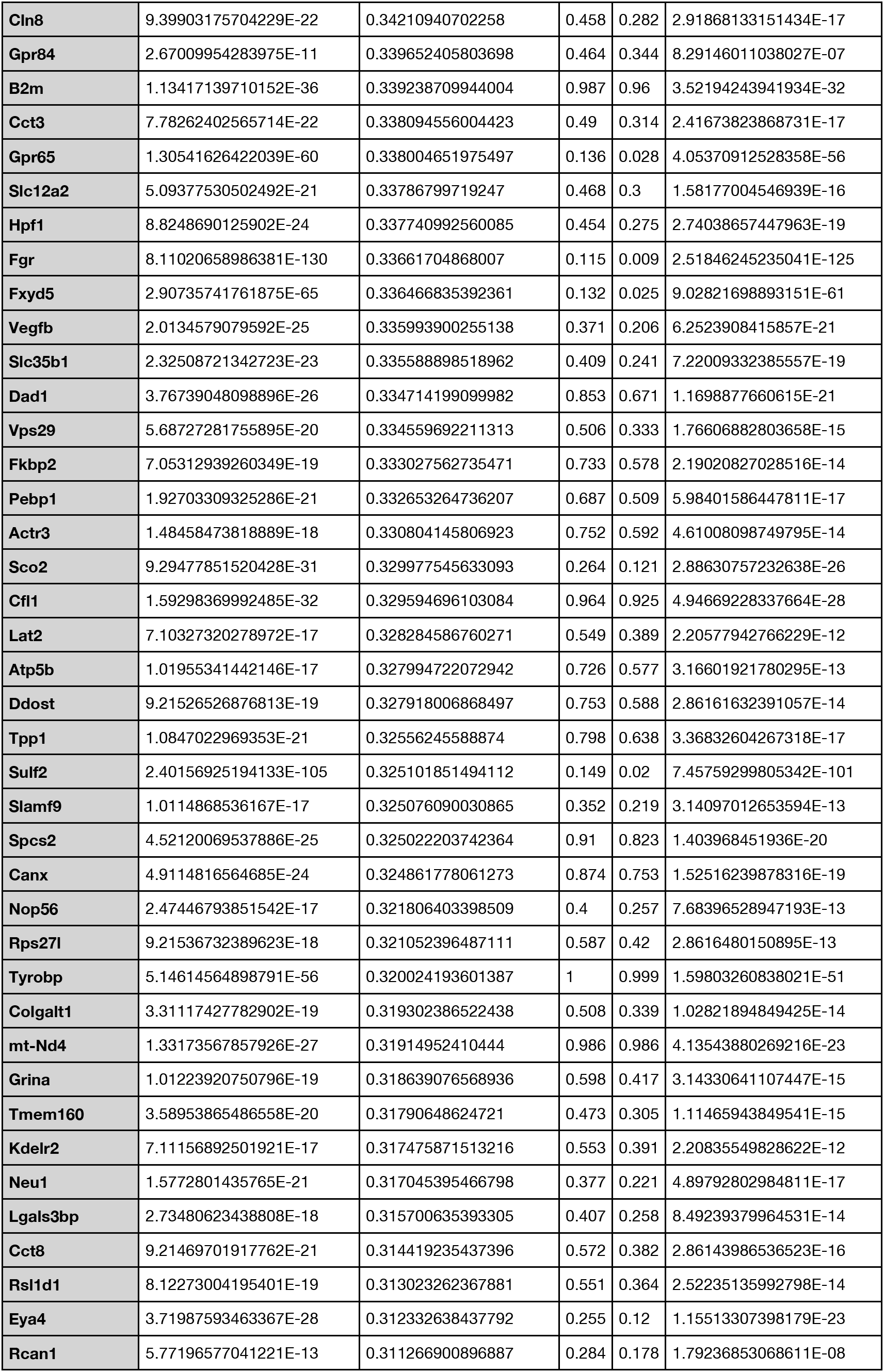
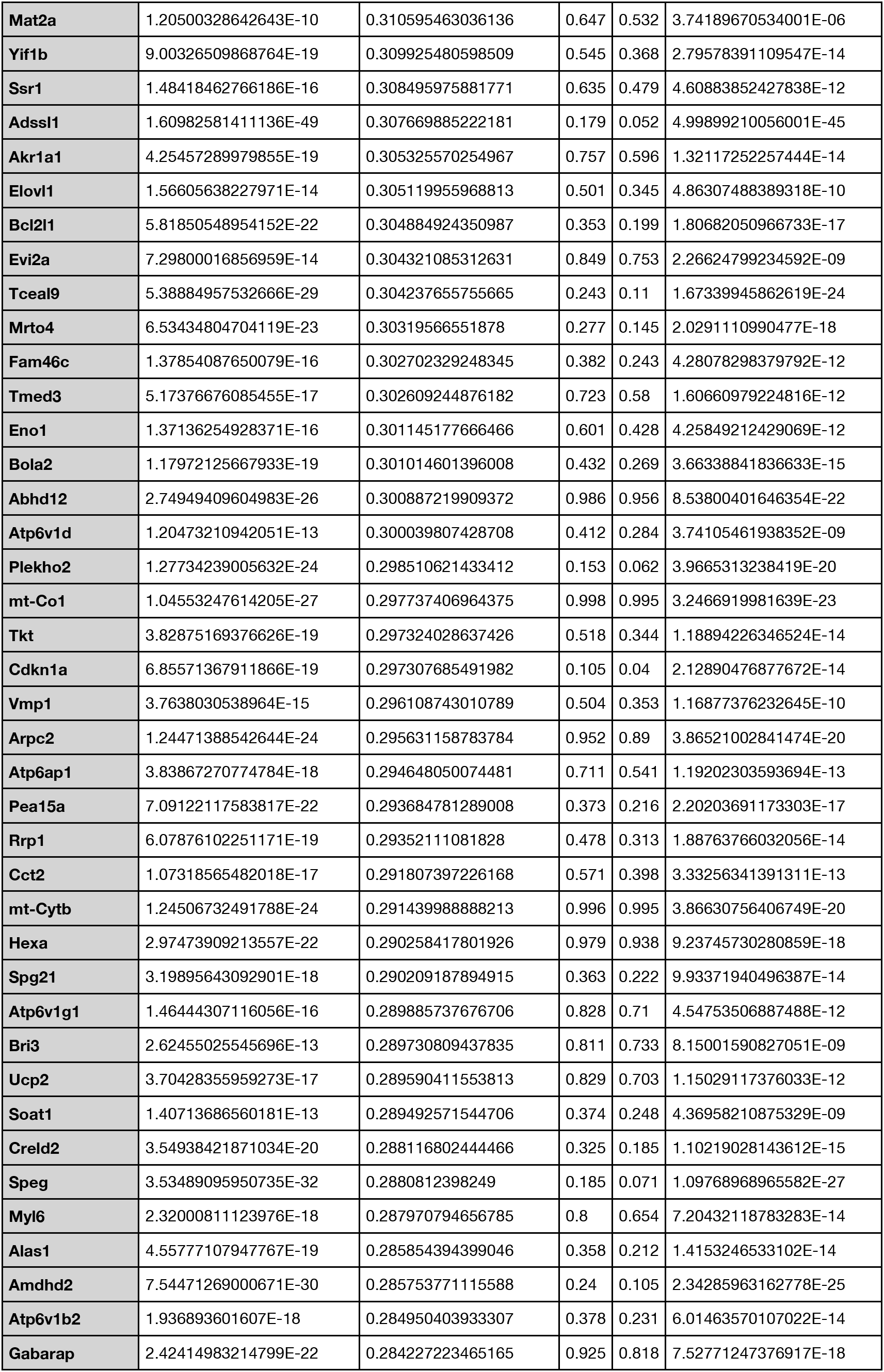
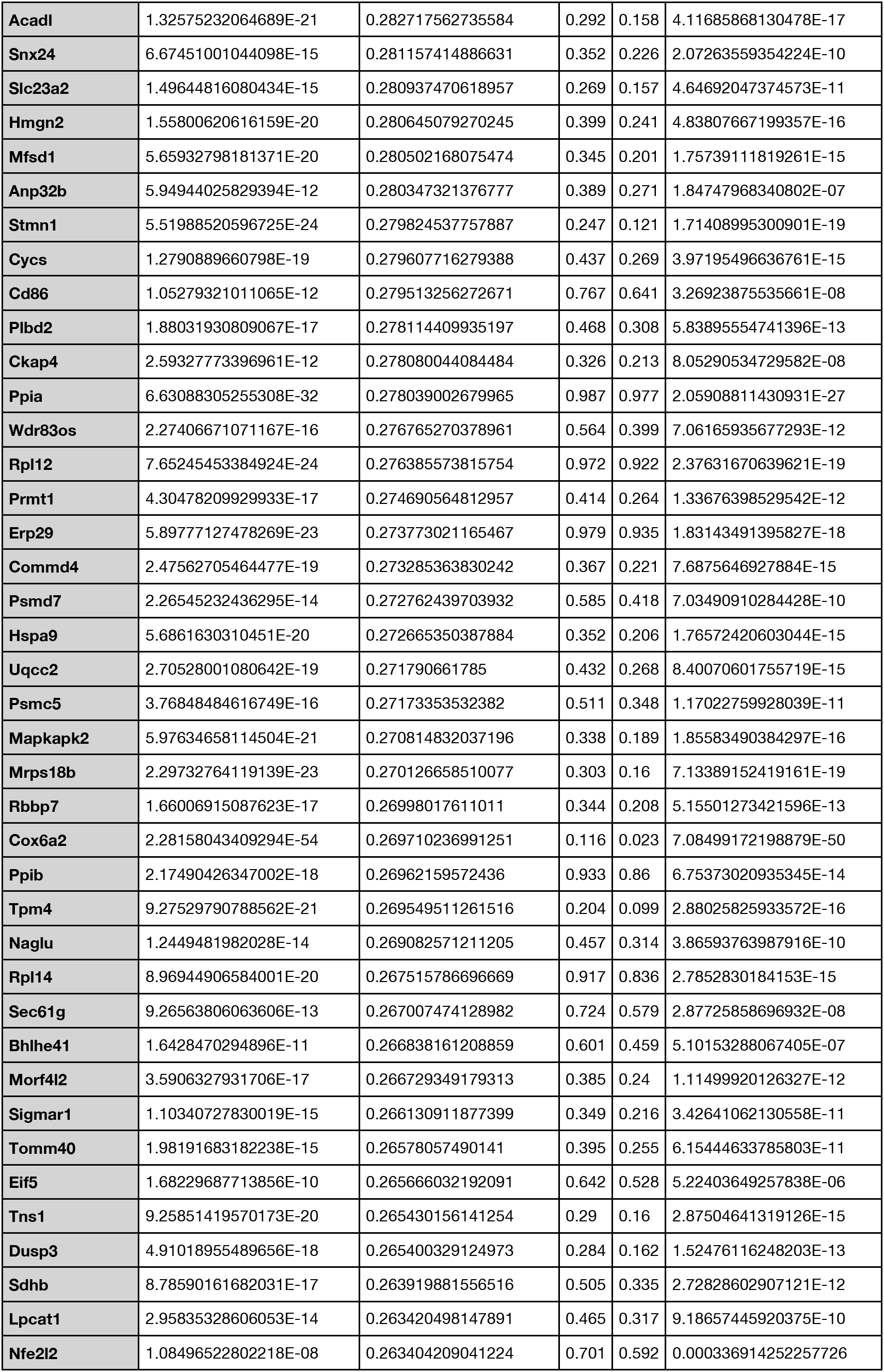
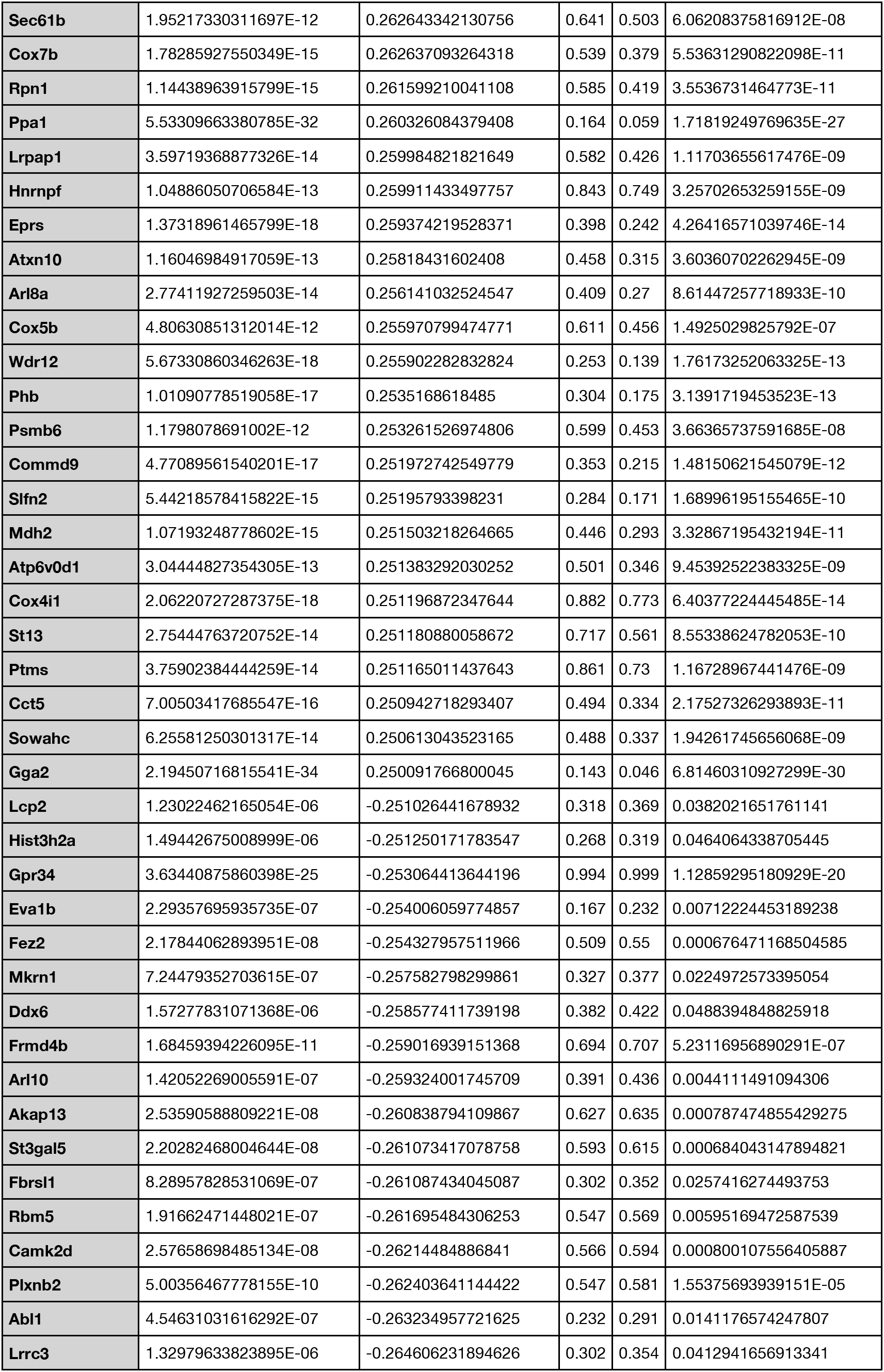
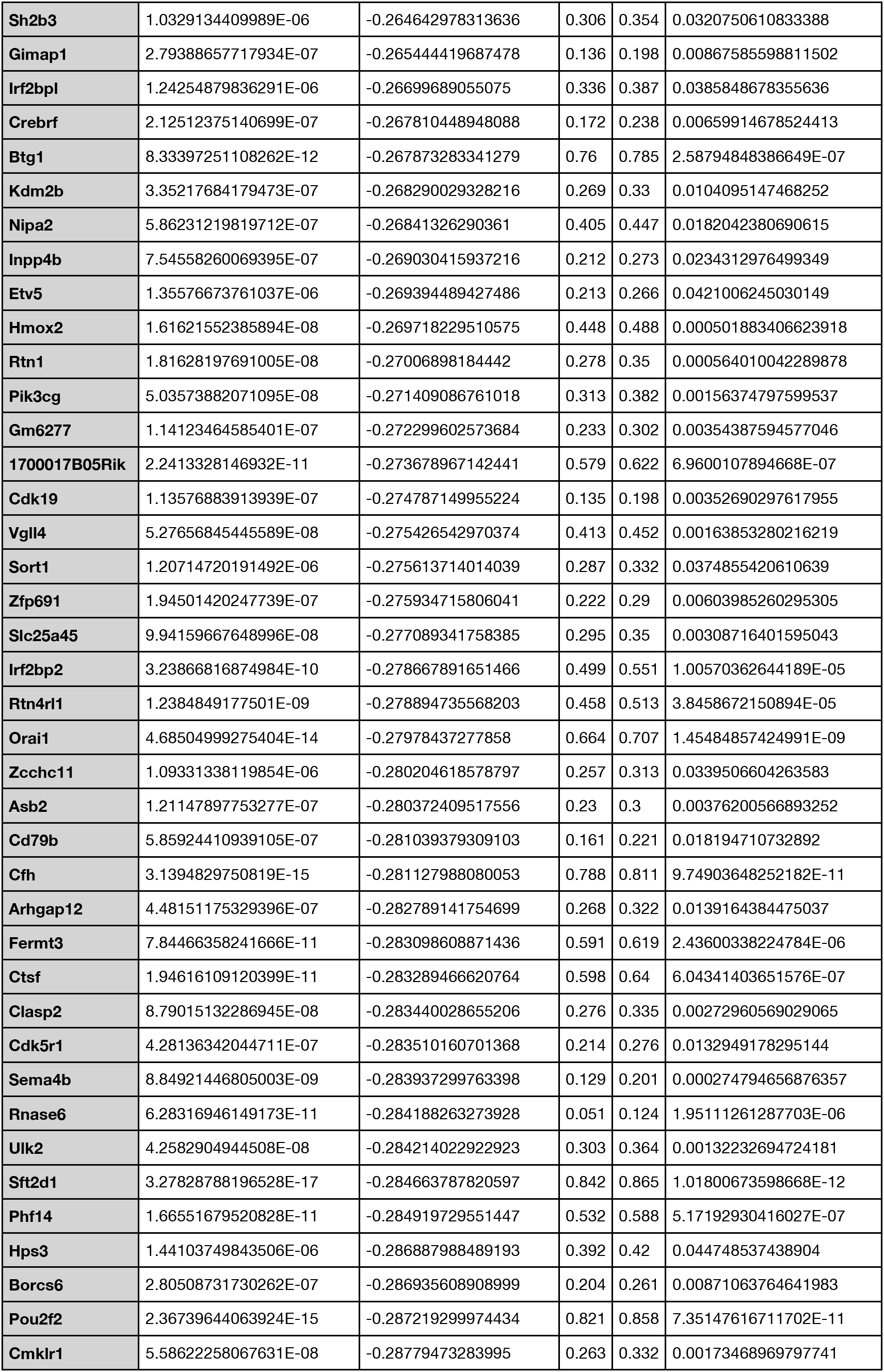
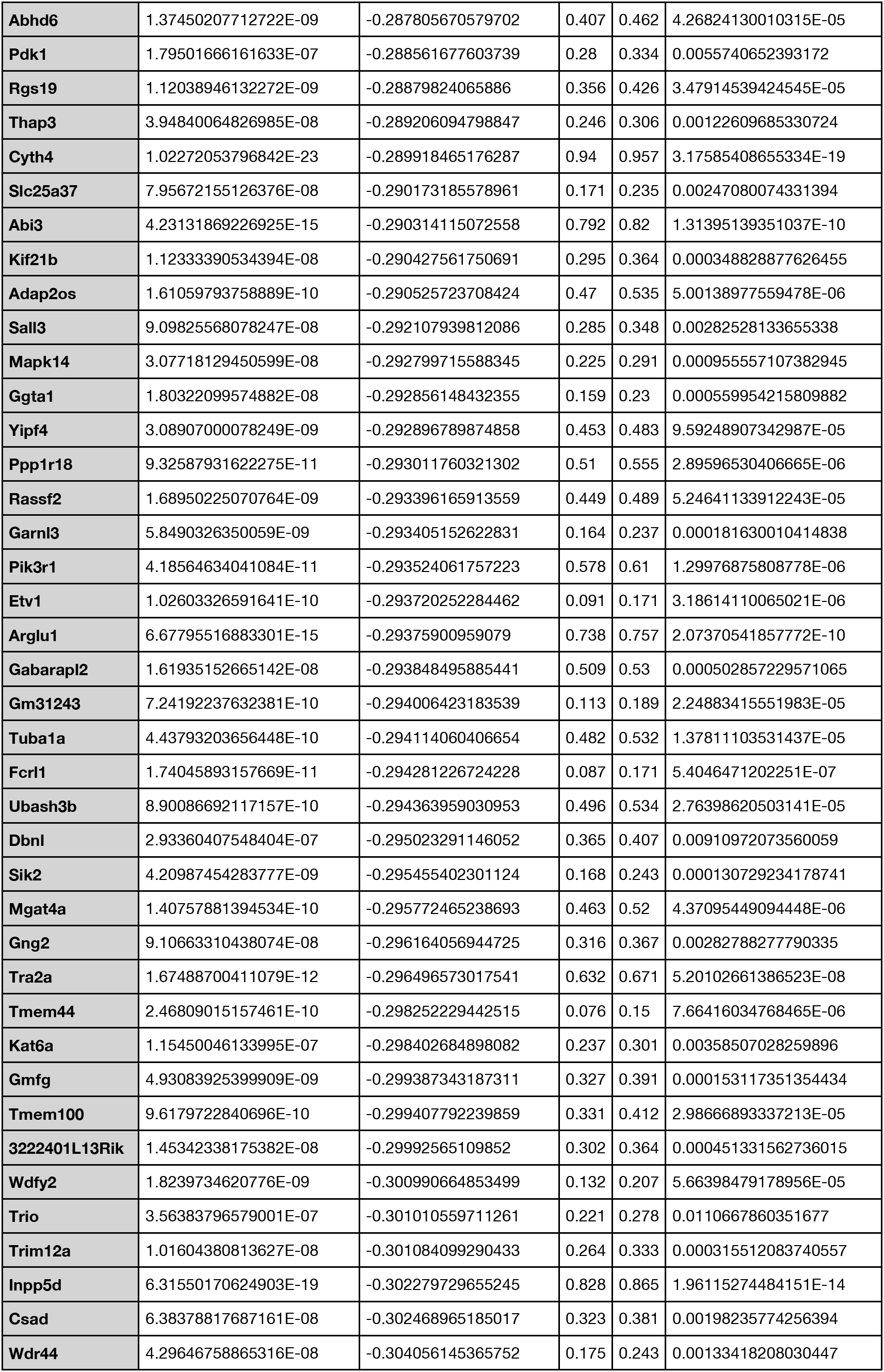
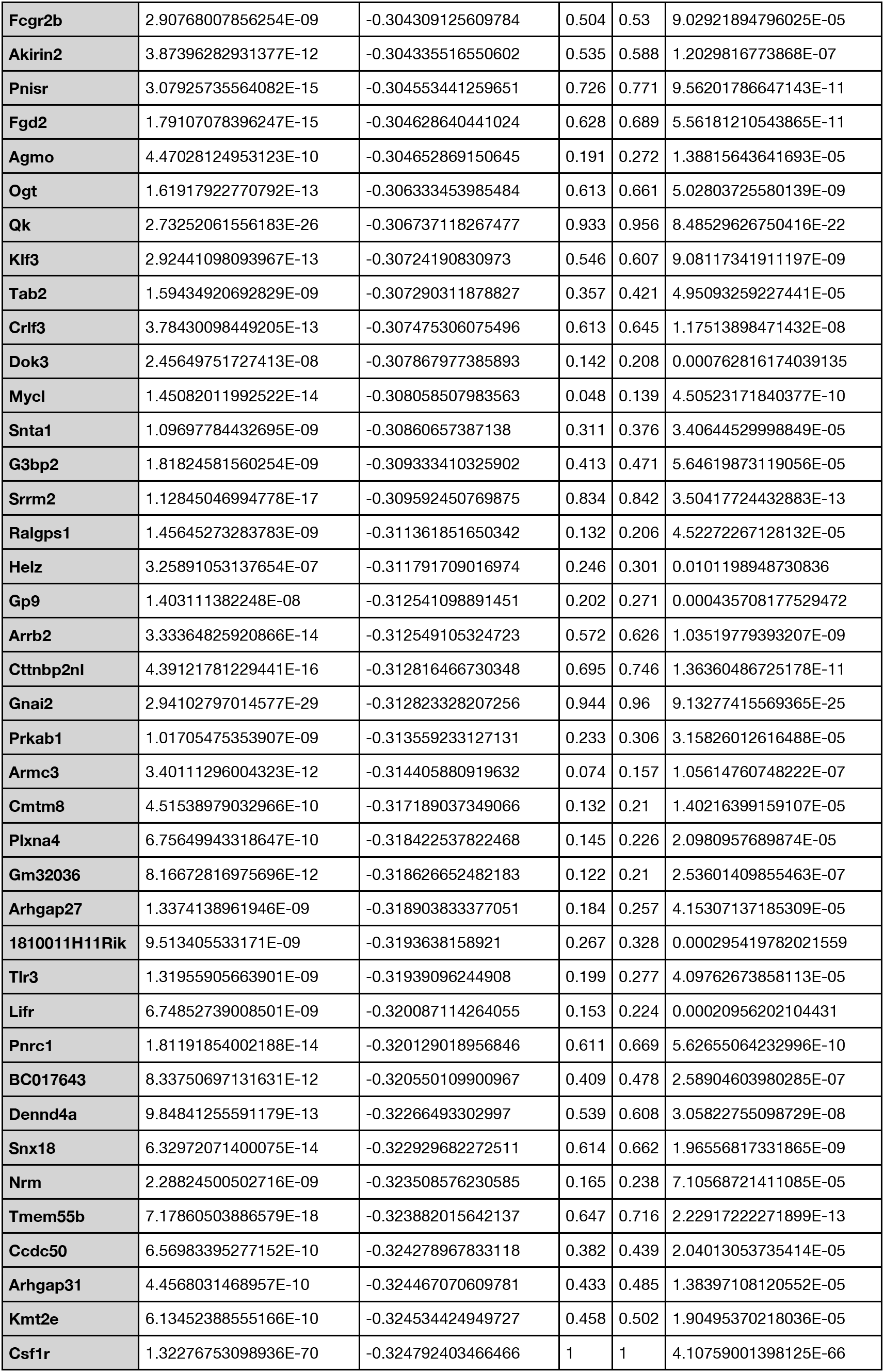
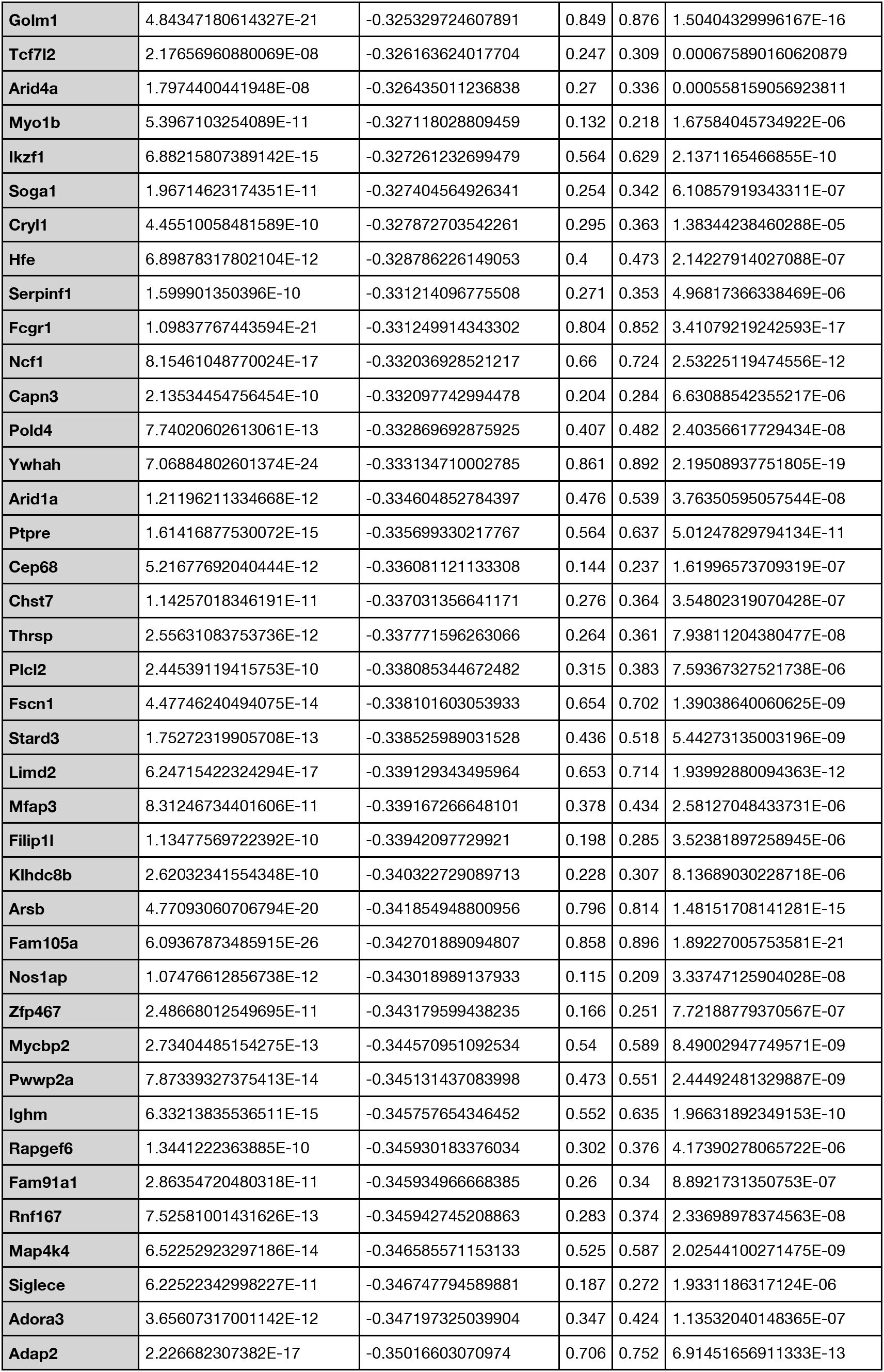
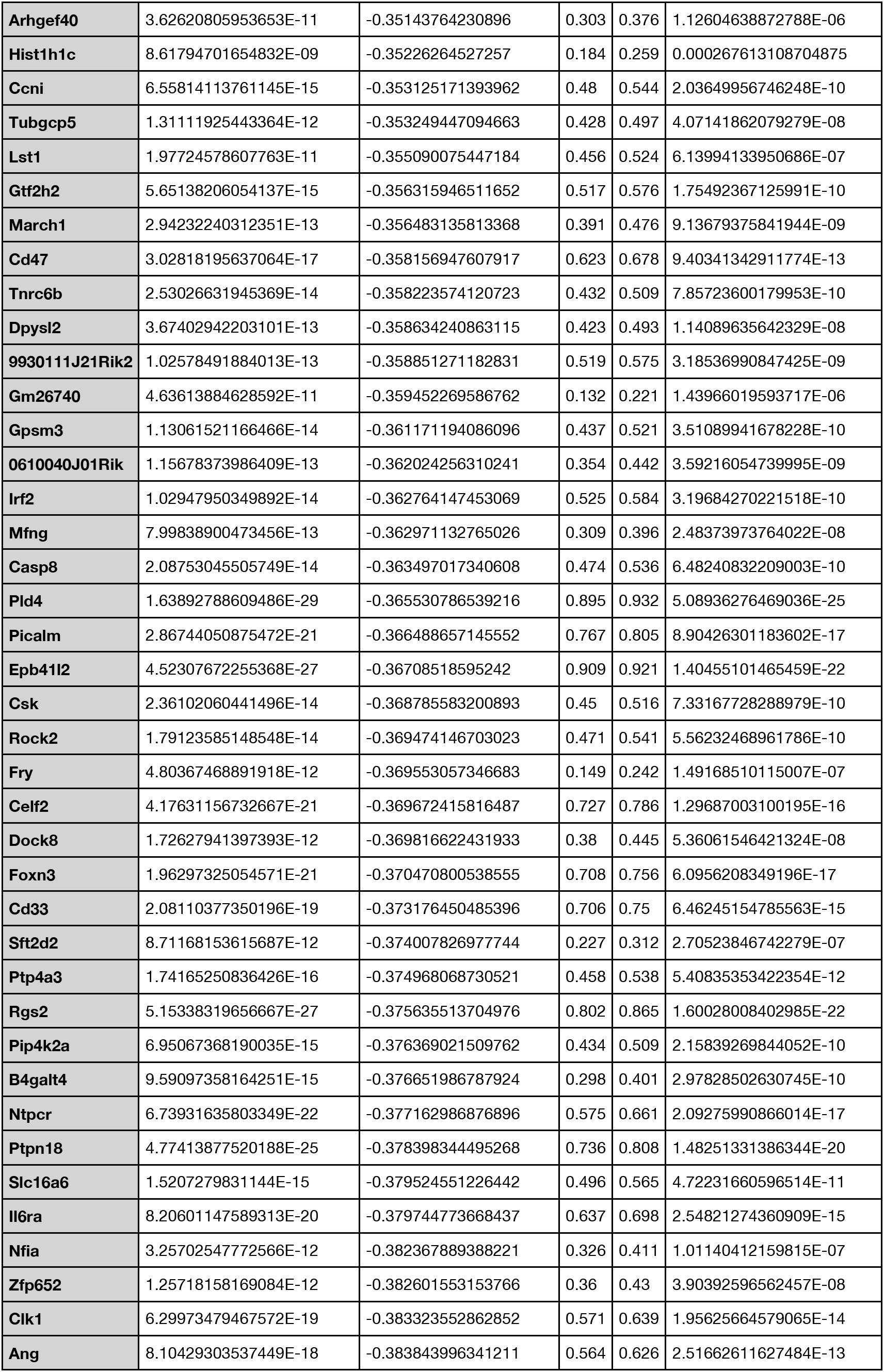
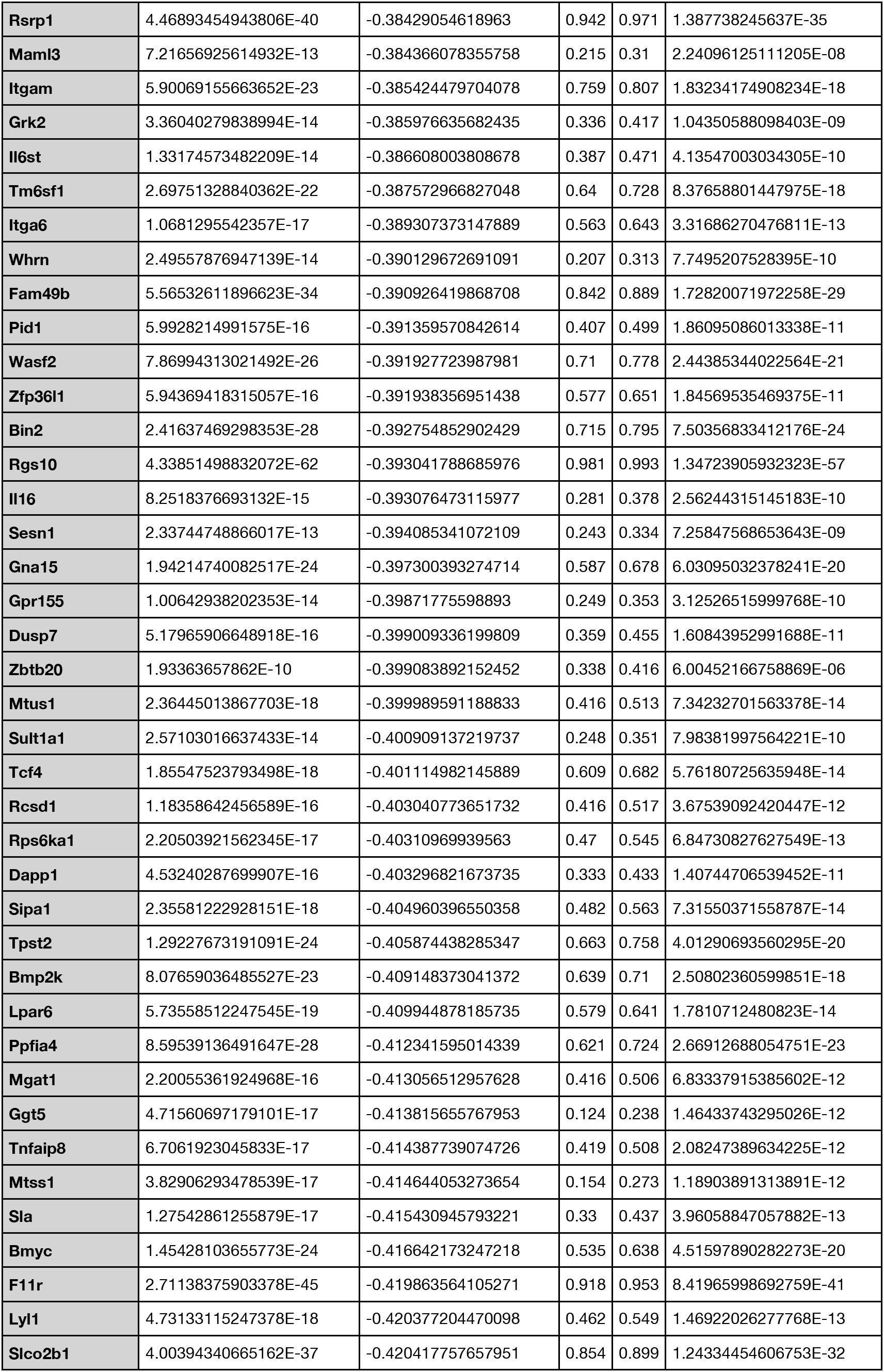
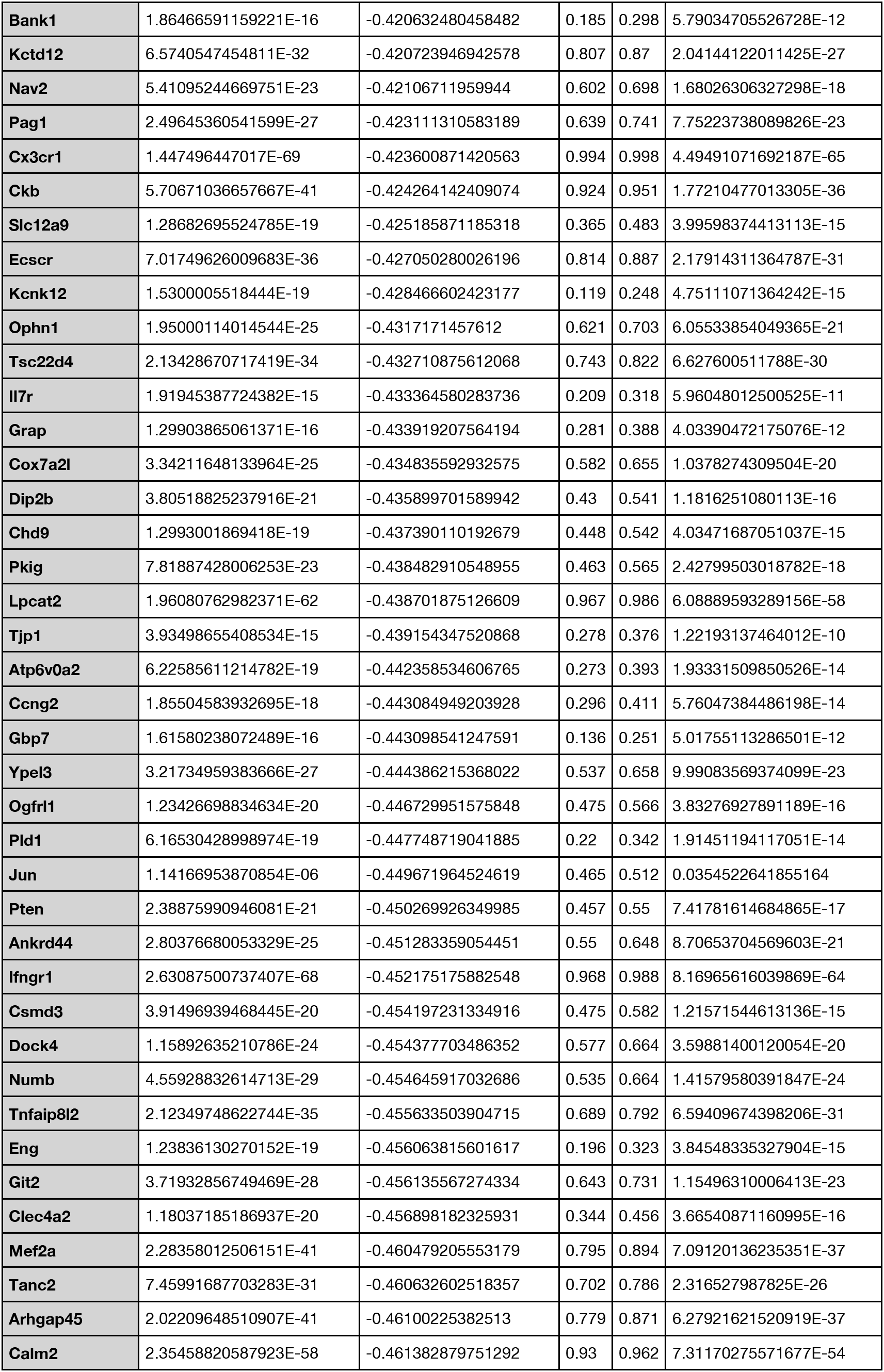
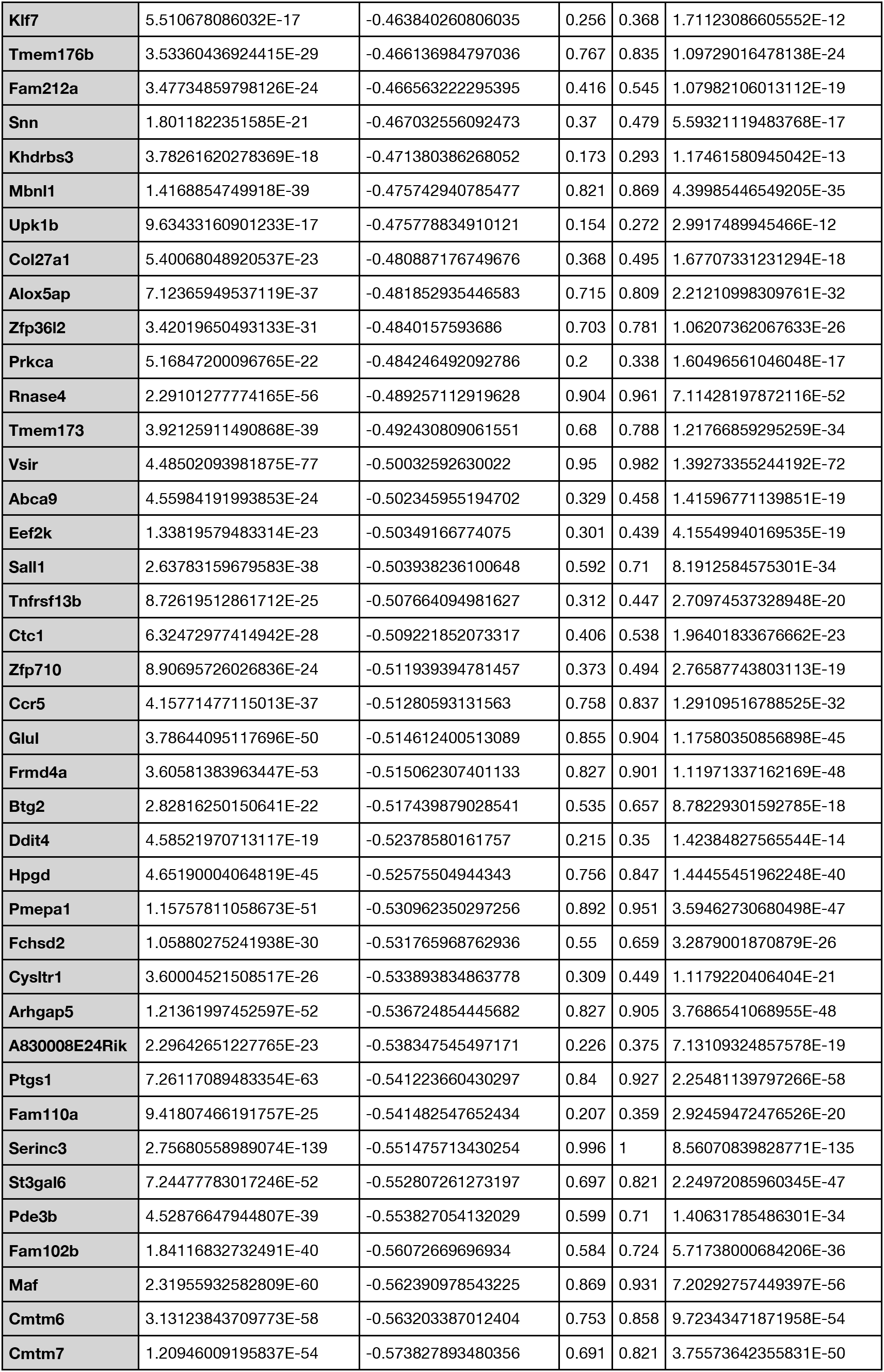
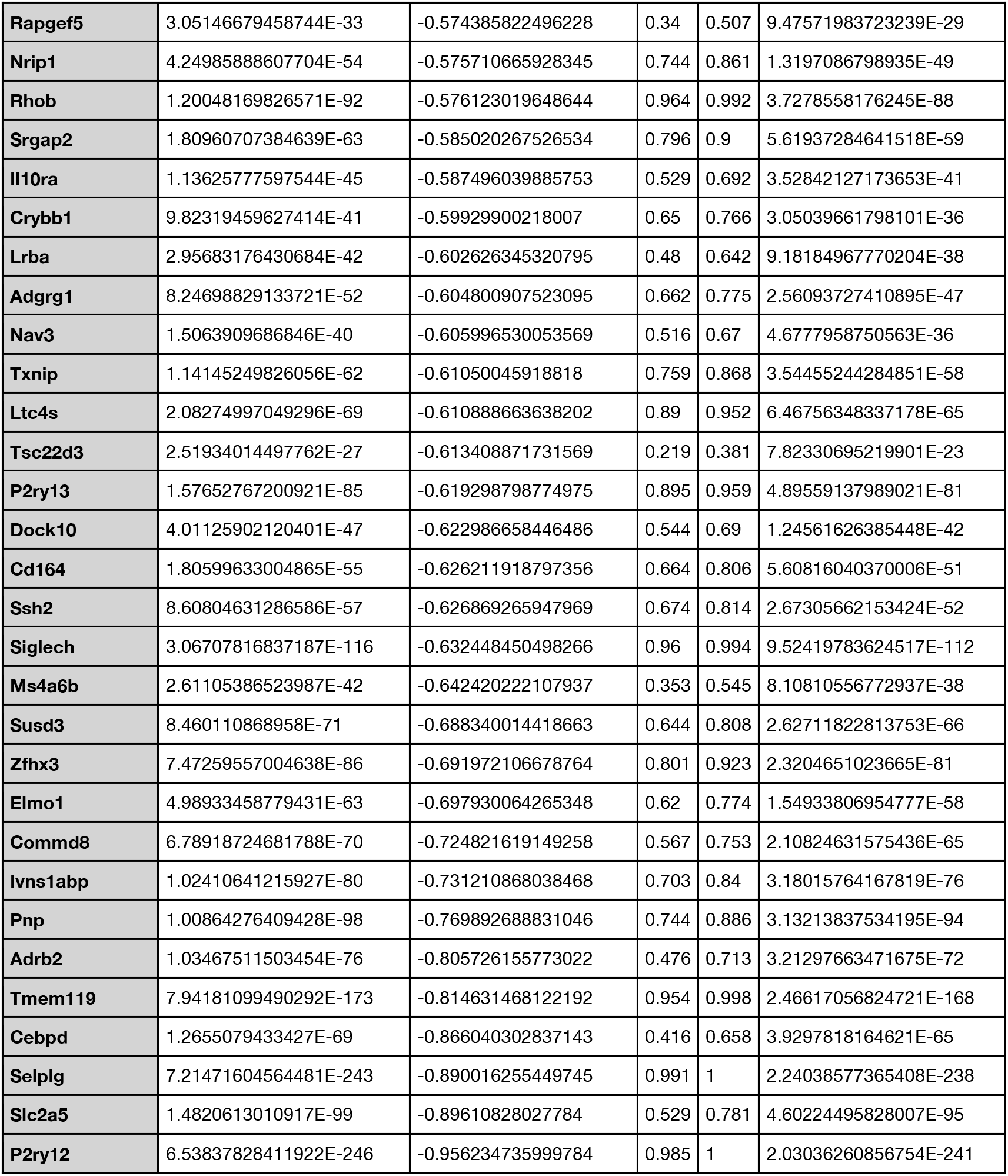
Top upregulated genes in SGZ cluster (8) versus homeostatic clusters (13 &14).

To assess whether these clusters truly distinguish cells based on biological differences, we next performed differential gene expression analysis between clusters (Table 1, Supplemental Figure 4). Due to the high enrichment of classical microglia-specific genes-*Tmem119, P2ry12, Selplg*-and low numbers of differentially expressed genes, we concluded that clusters 12,13,14, which represent about 72 percent of cells in the data set, reflect a homeostatic gene expression profile ^29, 30, 31^. We also inspected some key genes previously identified to be implicated in immune dynamics (Figure 1F) and then delved deeper into different transcriptomic clusters to determine to what extent these clusters reflect distinct populations ^10, 12, 29, 32^.

Clusters 4,5,6, and 11 did not meet the threshold for differential gene expression for gene set enrichment analysis. These clusters failed to show enrichment of at least ten nuclear genes. Cluster 4 shows high expression of mitochondrial genes. While these cells passed the threshold during processing, they may be indicative of cells that are lower quality. Cluster 5 was observed to express high levels of *Ccr1* and upregulate *Tmem176a*. These genes are found in border macrophages transitioning to a microglia-like state^12^. However, in our dataset these cells express microglia marker genes and no border macrophage genes such as *Mrc1*, *Ms4a7*, and *Pf4* and were detected. Cluster 6 expresses high levels of *Cdkn1a* and *Bax* suggesting that these cells may be undergoing apoptosis. Given that these clusters do not have many differentially expressed genes, they are unlikely to be distinct populations.

The remaining clusters express genes associated with either distinct functions or were characteristic of non-microglial populations. The presence of non-microglia macrophages has been reported in numerous studies^30, 32–35^. Cluster 2 appears to be a non-microglia macrophage population, based on, expression of genes associated with antigen presentation including *H2-Aa*, *H2-Eb1*, and *Cd74* in over half of the cells. These genes have been shown by previous findings to define adult choroid plexus macrophages^11^. The remaining cells in cluster 2 appear to have the same transcription profile associated with border macrophages or CNS-associated macrophages marked with expression of marker genes such as *Mrc1*, *Pf4*, and *Ms4a7* ^36^. These two groups thus likely comprise non-microglia myeloid lineage cells in the hippocampus.

In the resting brain, there is some degree of microglial turnover. Approximately one percent of cells in the total myeloid population in the hippocampus express genes undergoing cell cycle transition. These cells, enriched in Cluster 1, are expressing *Mki67*, *Top2a*, *H2afz*, known to be found in proliferating microglia^11^. Cluster 3 expresses genes known to be implicated in interferon response. Many of these genes, such as *Rtp4*, *Ifit3*, *Ifit2*, *Ifitm3*, *Oasl2,* have also been implicated in the aging transcriptome^12^. By contrast, cluster 9 comprises cells that exhibit an activation profile, potentially as an artifact of the isolation protocol, as evidenced by the upregulation of immediate early genes such as *Fos*, *Egr1*, and *Jun;* these genes have been shown to correlate specifically with activation due to homogenization and other isolation-associated experimental steps ^11^. Cluster 10 exhibits increased expression of genes associated with ribosomal subunits and also contains the highest *ApoE* expression. This group of cells undergoing high metabolic activity has been identified by another group but not characterized, to our knowledge^37^. Cluster 8 exhibits expression of genes associated with the Damage-associated microglial (DAM) phenotype, namely *Cd9*, *Cd63*, *Cst7*^8^ and more detailed analysis localizes this novel cluster to the dentate gyrus subgranular zone (Figures 3 and 4).

### Cd68 expression localizes to the subgranular zone of the dentate gyrus

In order to test whether microglia associated with the neurogenic niche are represented as a distinct transcriptomic cluster, we used our double reporter mice (Supplemental Figure 1) to visualize both neural progenitor cells and myeloid lineage cells. The neural progenitor pool clearly demarcates the SGZ from the rest of the dentate gyrus/hippocampus (Figure 2A). Somas from these cells line the interior (medial area) of the dentate gyrus. Processes of these cells protrude through the granule cell layer in the dentate gyrus to the molecular layer. Microglia in this region break the tile pattern they normally have in the adult cortex where processes of adjacent microglia seldomly coincide (Figure 2B & 2E). Such high density, or clustering, of microglia is often associated with increased microglial activity, particularly in clearing apoptotic cells^38, 39^. Additionally, neural stem cell processes are highly wrapped around microglial processes. Microglia are observed in very close apposition to neural progenitors in this region (Figure 2D). Cd68, (macrosialin) is found on the surface of lysosomes and increased Cd68 expression is detected in the SGZ ^40^. Cd68 puncta show higher colocalization in cells within the SGZ as well as processes from these cells (Figure 2C,D,G) as compared to microglia in the cortex or elsewhere in the dentate gyrus (Figure 2E & F). We analyzed the hippocampal clusters to see whether any transcriptomic cluster expresses elevated levels of Cd68 and observe that cluster 8 has significantly higher expression of Cd68 when compared to all other clusters (log2FC = 0.87; p-val <0.001) (Figure 2H).

**Figure 2:**
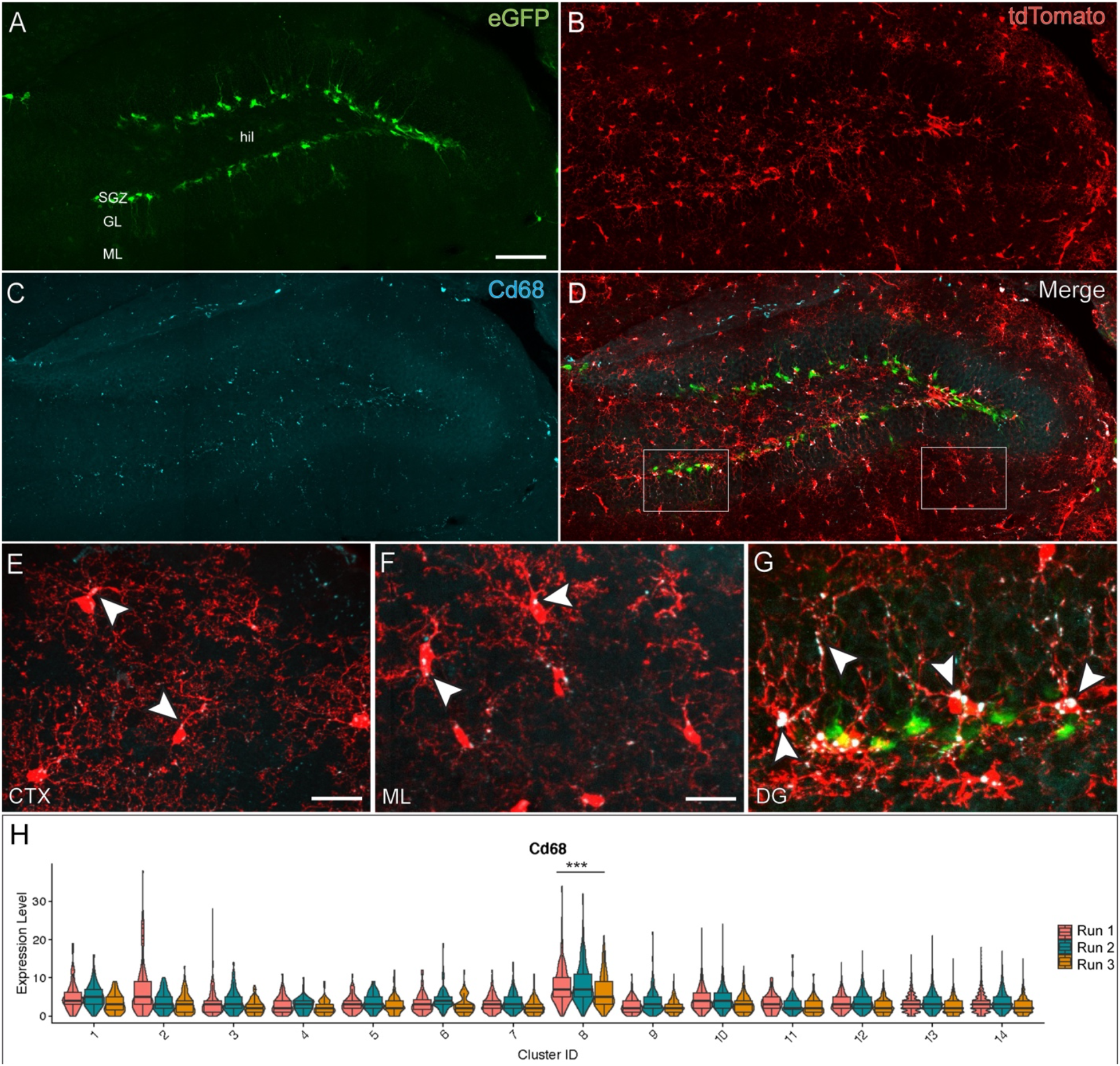
Increased Cd68 is localized to subgranular zone of the dentate gyrus and transcriptomic cluster 8. (A) Distribution of Nestin-eGFP expressing neural progenitors located in the SGZ. hil = hilus, GL = granule cell layer, ML = molecular layer. (B) Distribution of myeloid lineage cells expressing tdTomato in the dentate gyrus. (C) CD68^+^ lysosomal content staining in the dentate gyrus. (D) Merge image at showing colocalization of Cd68+ lysosomes in myeloid cells in apposition to Nestin-GFP cells. CD68^+^ lysosomal puncta in CTX (E) and ML (F) vs SGZ (G) with arrow heads to highlight CD68/tdTomato colocalization. Scale bars A&B = 40 µm; D-G: 100 µm; E= 25 µm F&G=: 20 µm. (H) Violin Plot with superimposed boxplots to show *Cd68* transcript counts and median values across clusters and runs.

### Transcriptomic analysis of the subgranular zone cluster demonstrates unique expression profile

After identifying Cluster 8 as the putative SGZ cluster, we tested for other genes differentially expressed by this cluster. We cross-referenced the Allen Institute *In Situ* Hybridization atlas to confirm spatial patterning of genes enriched or downregulated in this cluster (Supplemental Figure 7)^41^. Some previous studies have shown many microglia-specific marker genes to be downregulated in the context of immune activation^8^. Similarly, the SGZ microglia also exhibit decreased expression of microglia marker genes such as *Tmem119, P2ry12, Selplg,* and *Siglech* (Figure 3A). Interestingly, other microglia-specific genes such as *Hexb, Fclrs, Olfml3* retain stable expression in this cluster (Figure 3B).

**Figure 3:**
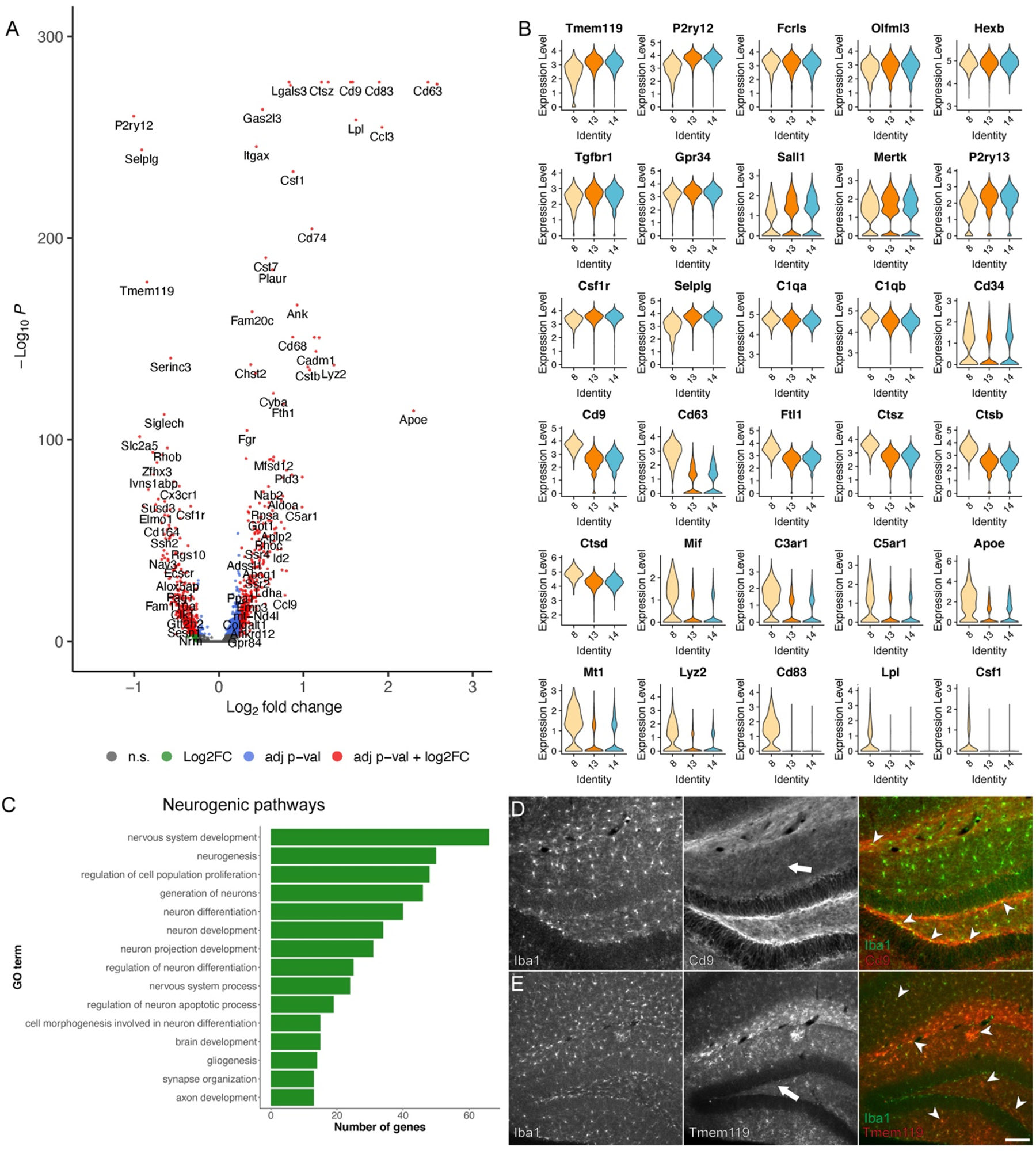
Phenotypic characteristics of SGZ microglia. (A) Volcano plot showing differentially expressed genes in SGZ microglia compared to other hippocampal microglia. Statistically significant genes (up or down-regulated) are represented by red dots (LFC > 0.25 and p-val < 10e-32). (B) Violin Plots showing comparison of expression profiles of key genes in SGZ cluster (cluster 8) vs homeostatic clusters (13 and 14). (C) Gene set enrichment analysis of upregulated genes in cluster 8 showing select ontology related to neurogenesis. (D) Immunohistochemistry staining for Cd9 colocalized with Iba1 in the dentate gyrus. Decreased immunoreactivity to Cd9 as indicated with an arrow in molecular layer (middle) with increased Iba1 colocalization in SGZ as indicated by arrowheads (right panel). (E) Immunohistochemistry staining for Tmem119 colocalized with Iba1 in the dentate gyrus. Decreased Tmem119 immunoreactivity in SGZ (arrow in middle panel) and sparse colocalization with Iba1 in SGZ indicated by arrowheads (right panel). Scale for D and E = 80 µm.

Cluster 8 shows an upregulation of genes associated with lysosomal function such as *Ctsz*, *Ctsb*, and *Ctsd* (Figure 3B). It is well established that complement receptors *C3ar1* and *C5ar1,* which are typically not found in resident microglia are expressed in the SGZ, and that complement cascade pathways are necessary for normal neuronal development and synaptic pruning ^42–44^. Therefore, it is unsurprising that we see them upregulated in cluster 8.

Many of these upregulated genes are implicated in nervous system development (Figure 3C), further suggesting that this cluster of cells correlates to microglia spatially aligned to the neurogenic nice, the SGZ of the dentate gyrus in the hippocampus. To validate whether these transcriptomic differences align with protein levels, we stained for candidate marker genes enriched in cluster 8. There is increased immunoreactivity to Cd9 in the SGZ (Figure 3D, Supplemental Figure 6). This is, however, not just localized to microglia. Conversely, *Tmem119* immunoreactivity is decreased in the SGZ (Figure 3E). We also observed decreased immunoreactivity of Iba1+ cell processes in the granular layer of the dentate gyrus, similar to reports in the subventricular zone where this other brain neurogenic region also demonstrates lower Iba1 expression.

We next utilized confocal imaging to test whether neural stem/progenitor cells express Cd9 using our double reporter mice. We found little to no colocalization of GFP with Cd9 staining (Supplemental Figure 6). We note increased staining around vessels, suggesting that Cd9 is enriched in vascularized regions such as the neurogenic niche also often referred to as the neurovascular niche. Lastly, we referred to the Allen Brain Atlas in situ hybridization database to confirm spatial localization of genes for which we could not find suitable antibodies. We note high specificity of *Cd63* in the SGZ, providing further evidence that this transcriptomic cluster is specific to the neurogenic niche of the hippocampus (Supplemental Figure 7).

### SGZ microglia display morphology and gene expression profiles consistent with a phagocytic phenotype

Microglial morphology and distribution are well established methods to compare activation states of immune cells ^15, 38, 39^. We first directly compared cells specifically in the sub granular zone with cells in the cortex. We and others have noted deviation in SGZ microglia from the tiled distribution of microglia found elsewhere (such as the cortex) (Figure 4A, 4B) in the homeostatic brain. To characterize morphometric traits, we utilized Sholl analysis to compare ramification of myeloid cells derived from the cortex versus the sub granular zone of the hippocampus. We find that cells with their cell bodies located in the sub granular zone are less ramified than cells in the cortex (Figure 4C), which is consistent with a phagocytic phenotype ^45, 46^. These results are in accordance with transcriptome analyses that suggest microglia display an alternative phenotype in the neurogenic niche ^25^.

**Figure 4:**
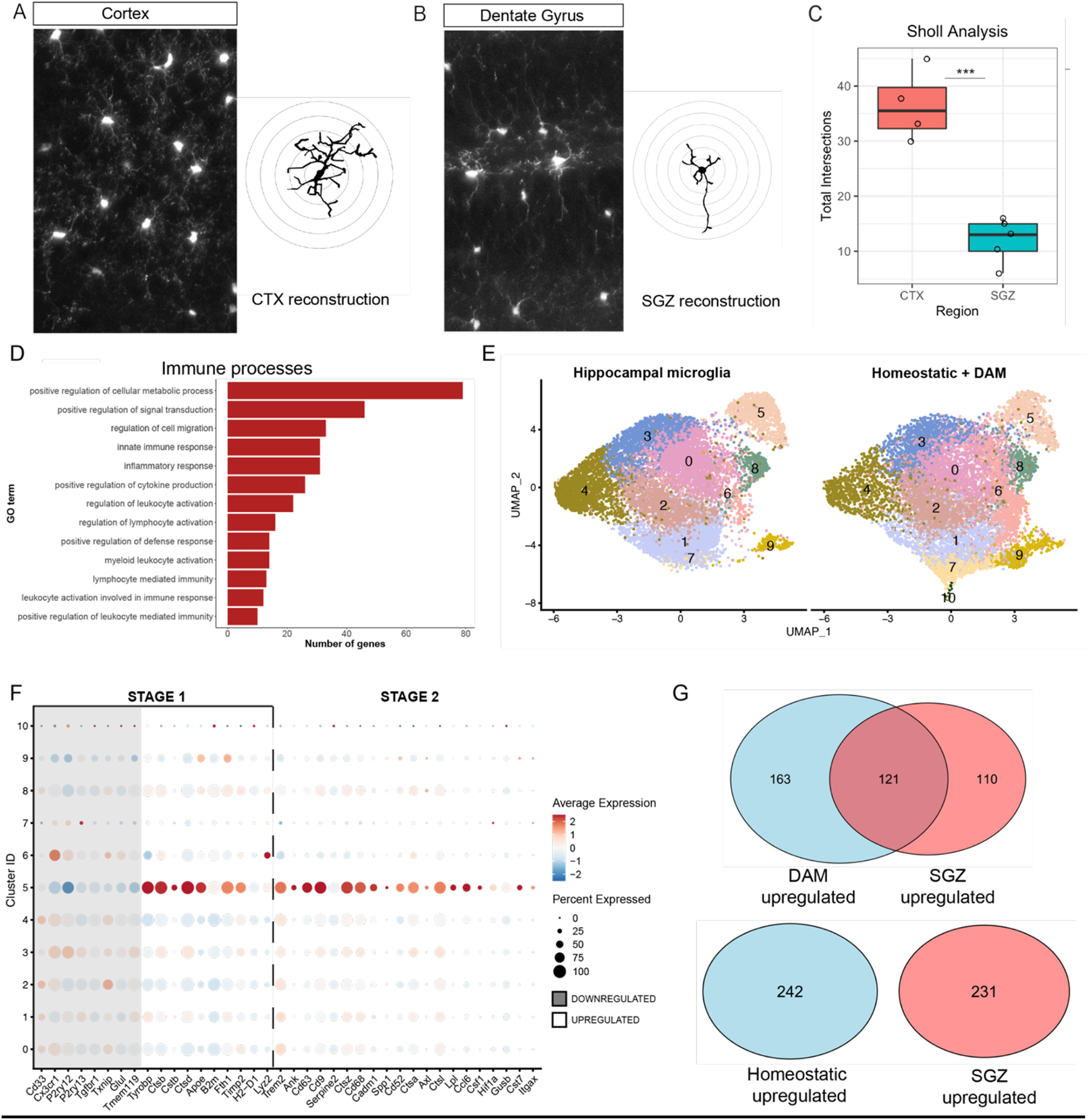
Subgranular zone (SGZ) microglia display a reactive phenotype. Representative projections of 3D z-stacked images of myeloid lineage cells labelled with tamoxifen induced tdTomato (*Cx3cr1^CreErt^*^2^*^/+^; Rosa26^loxp-tdTomato/+^*) in the cortex, CTX (A) and dentate gyrus (DG), with representative manual tracing of processes (right in A and B, respectively). (C) Sholl analysis to display morphometric differences between cell ramification of cells from cortex vs SGZ. p-val = 0.000227. (D) Gene set enrichment analysis of upregulated genes in cluster 4 showing select ontology related to immune activation. (E) UMAP plots showing integration of hippocampal myeloid dataset from this study (left) with microglial transcriptomic data adapted from ^8^ (right). (F) Dot Plot of key genes in integrated dataset (see E) known to be down regulated or upregulated in Stage-1 TREM2-independent activation versus Step-2 Trem2-dependent activation in DAM microglia. Shading corresponds to genes known to be downregulated in DAM profile (G) Venn diagram illustrating overlap of upregulated genes in SGZ clusters with Disease Associated Microglia (DAM) microglia (top) and overlap of upregulated genes in SGZ clusters with homeostatic microglia from ^8^ (bottom).

Using the cluster-8 specific gene list, we next examined the gene set enrichment analysis applying the Kolmogorov-Smirnov test through the topGO package for annotation of terms related to immune activation. We set a stringent false discovery rate (adjusted p-value less than 0.05) and found several gene ontology (GO) terms related to immune function (Figure 4D, Supplemental Table 2). We closely examined whether genes upregulated in the SGZ cluster overlapped with known activation profiles. We compared the SGZ transcriptome profile to that of Disease Associated Microglia (DAM) described in previous studies and found that microglia exclusively from the SGZ cluster with DAM (Figure 4E, Supplemental Figure 8) ^8^. We identified the DAM cluster by plotting key genes previously identified as Trem2-independent and Trem2-dependent (Figure 4F). We found genes upregulated primarily in the SGZ cluster with some expressed in the macrophage cluster. We noted extensive overlap between cells labelled as DAM in the Keren-Shaul dataset with the SGZ cluster in our study, with over half the genes enriched in the SGZ also enriched in DAM (Figure 4G). By contrast, we found no overlap between genes enriched in homeostatic cells from the Keren-Shaul dataset with those upregulated in the SGZ cluster (Figure 4G).

## Discussion

As the involvement of immune cells in CNS development and health are examined in greater depth, the role of innate immune cells reveals diverse function depending on cell type and context. Different cell types are tuned to recognize and respond to specific cues. Some populations present in the brain express different sets of genes which may explain mechanisms involving aberrant immune activation and inflammation leading to neurodegenerative diseases coupled with cognitive decline. Immune cells are undoubtedly an important layer of defense for the central nervous system, while dysregulation of the neuroimmune axis can disrupt healthy neural and cognitive functions. This makes characterizing the different players in the immune system key to uncovering pathological development and recovery in neurological diseases. By sampling a large number of hippocampal cells, we provide evidence of heterogeneity in the adult hippocampus, which may help explain specialized roles for microglia in mitigating disease within discrete brain areas such as the hippocampal neurogenic niche.

We have uncovered a novel context in which immune cell heterogeneity explains a specialized function. The subgranular zone of the hippocampus is a select site in the adult mammalian brain where neuronal development persists through adulthood. Thus, we have characterized a key component of this neurogenic niche, a specialized subpopulation of microglia with a distinct transcriptomic signature correlating to function within this niche. This population of cells comprises less than five percent of total immune cells in the hippocampus, highlighting why it may have been missed in studies that examine a smaller number of cells in the hippocampus.

Previous studies suggest that immune dynamics alter the neurogenic niche and immune input in this region appears highly specialized ^13, 14, 25^. In the SGZ, cells labelled with tdTomato (resulting from fractalkine Cre recombinase reporter) show stark morphological differences when compared to cells from non-neurogenic regions. Similar to microglia in the adult subventricular zone, these cells exhibit altered immunoreactivity to some common markers such as Tmem119 and Iba1. These cells, do however, express Cd68+ puncta, which are generally found only in low levels by microglia that are ramified or surveilling. It has been long known that microglia show spatial patterning in morphology and density across the brain^38^. These morphological differences often correlate with differential phagocytic activity, proliferative potential, and immunoreactivity as different regions of the brain have different needs.

In this study, we characterize the transcriptomic profile of these cells relative to other myeloid cells in the hippocampus at the level of single cell resolution. We identify a greater number of differentially expressed genes by subgranular zone microglia, and separate confounding results from CNS-associated macrophages. This approach allows us to computationally separate the immune population associated with the neurogenic niche of the hippocampal formation with great precision.

We first noted downregulation of several microglia-specific marker genes in the healthy, adult subgranular zone; strikingly similar to cases of disease and injury ^47, 48^. Due to the lower levels of expression of these marker genes, these cells may not be captured by conventional flow cytometry panels, requiring judicious use of marker genes depending on spatial, temporal, and associated disease context. We classify myeloid cells in the SGZ as microglia as they still retain expression, albeit to a lesser degree, of some microglia-specific genes while maintaining robust expression of other marker genes such as *Hexb* and *Olfm3*. This further corroborates findings which suggest not all microglia-specific markers retain consistent expression and comprehensively label microglia in the brain^47^.

Next, we compared genes upregulated in the SGZ to known microglial phenotypes. Studies have reported that microglia described as alternatively activated M2 microglia in the macrophage polarization scheme promote neurogenesis ^49–51^. Microglia in the dentate gyrus have also been reported to express a handful of genes in M2 microglia^13, 25^. Our results indicate that this profile does not accurately capture the SGZ transcriptome. Instead, we observe great overlap of genes enriched in our putative SGZ cluster with those upregulated in disease-associated microglia (DAM), which are involved in plaque clearance^8, 52^. This DAM signature is translationally relevant and potentially highly conserved, as it has been observed in postmortem human tissue from patients with AD and MS^9, 10^.

Many of these DAM genes are also found in proliferation-associated macrophages (PAM) found in developing white matter ^8, 12^. These are primarily genes which are associated with lysosomal function. In PAM these genes reflect phagocytosis of oligodendrocyte progenitors, important in myelination, and are no longer present in the mature, adult brain. The presence of these genes in the SGZ presumably reflects engulfment of excess neural progenitors that are undergoing apoptosis ^16, 53^. This points to a broader role of DAM genes extending beyond disease, particularly since phagocytic microglia have been shown to support neuronal development in the adult hippocampus^16^.

During early postnatal development, developing neurons are pruned through a CD11B-DAP12-dependent mechanism^54^. Interestingly, our SGZ also shows upregulation of *Dap12* (also known as *Tyrobp*), which is activated downstream of the DAM-specific receptor TREM2. Our data suggest that processes similar to those found in embryonic development of the nervous system as well as in PAMs during postnatal development persist throughout adulthood in the SGZ. This population specifically may be targeted in diseases in which neurogenesis is perturbed or needs to be altered^18, 24^. Notably, gene networks associated with phagocytic microglia are involved in neurogenic function *in vivo* within the neurogenic niche^16^.

We also ruled out whether transcriptomic differences in SGZ microglia could explain how sex-specific differences may contribute to altered risk for neurodevelopment diseases, particularly those affecting hippocampal function and neurogenesis ^55–58^. In the healthy adult murine hippocampus, there appear to be no sex-related differences in the transcription of immune-related genes Additionally, no significant differences in genes related to immune function are found specifically in SGZ microglia from male and female mice. Thus, differences in immune function that occur may come into play later in life or are due to post-transcriptional regulation or at the level of protein interactions.

Adult hippocampal neurogenesis is sensitive to various external or environmental factors. As such, the immune cells in this niche are tuned to varying inputs and may be responsible for relaying the state of the outside world to progenitors. Important questions that remain to be answered are whether this specialized microglial phenotype is indicative of a distinct ontogeny, or whether it arises as a response to the environmental cues provided by the niche. Furthermore, whether this signature describes an activated population or a reactive state is not known and of great interest ^59^. This is important to uncover because it can elucidate if alterations in adult neurogenesis in pathological contexts result from differential properties of immune cells in the SGZ. Given that these cells express many genes found in the DAM gene signature, the role of these genes specifically in the context of adult neurogenesis need to be examined.

While some genes expressed by microglia have been characterized in various developmental stages and disease models, many of the genes enriched by subgranular zone microglia have unknown functions. Our findings point to a need to separate the SGZ population and examine it separately in disease models. Extracting SGZ-specific differences is a necessary component of understanding hippocampal physiology, particularly from an immune perspective.

Further understanding the mechanisms behind how this population progresses and the trajectory it takes will be fundamentally important, particularly in the context of aging and disease ^57, 60^ While we highlight one specific subset of cells in this paper, the potential contributions of other populations cannot be excluded from regulation of the neurogenic niche or in disease development and progression. Our data provide a reference point for comparing immune alterations in the hippocampus and its neurogenic niche in the context of disease or other pathology.

The authors declare no competing financial interests or other conflict of interest.

## Acknowledgements

Acknowledgements

This research was supported by National Institutes of Health/National Institute of Neurological Disorders and Stroke Grants R01-NS-095803 (SGK) and the Paul Allen Foundation (SGK), and National Institute of Aging Grant R01-AG-066831 (VM). This research was funded in part through the NIH/NCI Cancer Center Support Grant P30CA013696 and used the Genomics and High Throughput Screening Shared Resource. Research reported in this publication was performed in the CCTI Flow Cytometry Core, supported in part by the Office of the Director, National Institutes of Health under awards S10OD020056. The content is solely the responsibility of the authors and does not necessarily represent the official views of the National Institutes of Health. Images were collected and/or image processing and analysis for this work was performed in the Confocal and Specialized Microscopy Shared Resource of the Herbert Irving Comprehensive Cancer Center at Columbia University, supported by NIH grant #P30 CA013696 (National Cancer Institute).

## Author Contributions

SK, EMB, and SC conceptualized this project. SC performed all experiments. NV assisted with tissue homogenization and microglia isolation. VM helped design the analysis framework. SC and PG processed and analyzed data. PG performed Harmony integration for comparison with DAM dataset. SC generated figures and wrote manuscript. SK,PG, EMB, and VM revised manuscript.

## References

1. Moser, E. I., Moser, M. B. & McNaughton, B. L. Spatial representation in the hippocampal formation: a history. Nat Neurosci 20, 1448–1464, doi:10.1038/nn.4653 (2017).

2. Eichenbaum, H. & Cohen, N. J. Can we reconcile the declarative memory and spatial navigation views on hippocampal function? Neuron 83, 764–770, doi:10.1016/j.neuron.2014.07.032 (2014).

3. Burgess, N., Maguire, E. A. & O’Keefe, J. The human hippocampus and spatial and episodic memory. Neuron 35, 625–641, doi:10.1016/s0896-6273(02)00830-9 (2002).

4. Efthymiou, A. G. & Goate, A. M. Late onset Alzheimer’s disease genetics implicates microglial pathways in disease risk. Mol Neurodegener 12, 43, doi:10.1186/s13024-017-0184-x (2017).

5. Pluvinage, J. V. & Wyss-Coray, T. Systemic factors as mediators of brain homeostasis, ageing and neurodegeneration. Nat Rev Neurosci 21, 93–102, doi:10.1038/s41583-019-0255-9 (2020).

6. Wyss-Coray, T. & Mucke, L. Ibuprofen, inflammation and Alzheimer disease. Nat Med 6, 973–974, doi:10.1038/79661 (2000).

7. Mosher, K. I. & Wyss-Coray, T. Microglial dysfunction in brain aging and Alzheimer’s disease. Biochem Pharmacol 88, 594–604, doi:10.1016/j.bcp.2014.01.008 (2014).

8. Keren-Shaul, H. et al. A Unique Microglia Type Associated with Restricting Development of Alzheimer’s Disease. Cell 169, 1276–1290 e1217, doi:10.1016/j.cell.2017.05.018 (2017).

9. Olah, M. et al. Single cell RNA sequencing of human microglia uncovers a subset associated with Alzheimer’s disease. Nat Commun 11, 6129, doi:10.1038/s41467-020-19737-2 (2020).

10. Masuda, T. et al. Spatial and temporal heterogeneity of mouse and human microglia at single-cell resolution. Nature 566, 388–392, doi:10.1038/s41586-019-0924-x (2019).

11. Li, Q. et al. Developmental Heterogeneity of Microglia and Brain Myeloid Cells Revealed by Deep Single-Cell RNA Sequencing. Neuron 101, 207–223 e210, doi:10.1016/j.neuron.2018.12.006 (2019).

12. Hammond, T. R. et al. Single-Cell RNA Sequencing of Microglia throughout the Mouse Lifespan and in the Injured Brain Reveals Complex Cell-State Changes. Immunity 50, 253–271 e256, doi:10.1016/j.immuni.2018.11.004 (2019).

13. Kreisel, T., Wolf, B., Keshet, E. & Licht, T. Unique role for dentate gyrus microglia in neuroblast survival and in VEGF-induced activation. Glia 67, 594–618, doi:10.1002/glia.23505 (2019).

14. Marshall, G. P., 2nd, Deleyrolle, L. P., Reynolds, B. A., Steindler, D. A. & Laywell, >E. D. Microglia from neurogenic and non-neurogenic regions display differential proliferative potential and neuroblast support. Front Cell Neurosci 8, 180, doi:10.3389/fncel.2014.00180 (2014).

15. Ribeiro Xavier A. L., Kress, B. T., Goldman, S.A. Lacerda de Menezes, J. R. & Nedergaard, M. A Distinct Population of Microglia Supports Adult Neurogenesis in the Subventricular Zone. J Neurosci 35, 11848–11861, doi:10.1523/JNEUROSCI.1217-15.2015 (2015).

16. Diaz-Aparicio, I. et al. Microglia Actively Remodel Adult Hippocampal Neurogenesis through the Phagocytosis Secretome. J Neurosci 40, 1453–1482, doi:10.1523/JNEUROSCI.0993-19.2019 (2020).

17. Chintamen, S., Imessadouene, F. & Kernie, S. G. Immune Regulation of Adult Neurogenic Niches in Health and Disease. Front Cell Neurosci 14, 571071, doi:10.3389/fncel.2020.571071 (2020).

18. Willis, E. F. et al. Repopulating Microglia Promote Brain Repair in an IL-6-Dependent Manner. Cell 180, 833–846 e816, doi:10.1016/j.cell.2020.02.013 (2020).

19. Blaiss, C. A. et al. Temporally specified genetic ablation of neurogenesis impairs cognitive recovery after traumatic brain injury. J Neurosci 31, 4906–4916, doi:10.1523/JNEUROSCI.5265-10.2011 (2011).

20. Scopa, C. et al. Impaired adult neurogenesis is an early event in Alzheimer’s disease neurodegeneration, mediated by intracellular Abeta oligomers. Cell Death Differ 27, 934–948, doi:10.1038/s41418-019-0409-3 (2020).

21. Moreno-Jimenez, E. P. et al. Adult hippocampal neurogenesis is abundant in neurologically healthy subjects and drops sharply in patients with Alzheimer’s disease. Nat Med 25, 554–560, doi:10.1038/s41591-019-0375-9 (2019).

22. Dranovsky, A. & Hen, R. Hippocampal neurogenesis: regulation by stress and antidepressants. Biol Psychiatry 59, 1136–1143, doi:10.1016/j.biopsych.2006.03.082 (2006).

23. Choi, S. H. et al. Combined adult neurogenesis and BDNF mimic exercise effects on cognition in an Alzheimer’s mouse model. Science 361, doi:10.1126/science.aan8821 (2018).

24. Berdugo-Vega, G. et al. Increasing neurogenesis refines hippocampal activity rejuvenating navigational learning strategies and contextual memory throughout life. Nat Commun 11, 135, doi:10.1038/s41467-019-14026-z (2020).

25. Artegiani, B. et al. A Single-Cell RNA Sequencing Study Reveals Cellular and Molecular Dynamics of the Hippocampal Neurogenic Niche. Cell Rep 21, 3271–3284, doi:10.1016/j.celrep.2017.11.050 (2017).

26. Yu, T. S., Zhang, G., Liebl, D. J. & Kernie, S. G. Traumatic brain injury-induced hippocampal neurogenesis requires activation of early nestin-expressing progenitors. J Neurosci 28, 12901–12912, doi:10.1523/JNEUROSCI.4629-08.2008 (2008).

27. Bohlen, C. J., Bennett, F. C. & Bennett, M. L. Isolation and Culture of Microglia. Curr Protoc Immunol 125, e70, doi:10.1002/cpim.70 (2019).

28. Yona, S. et al. Fate mapping reveals origins and dynamics of monocytes and tissue macrophages under homeostasis. Immunity 38, 79–91, doi:10.1016/j.immuni.2012.12.001 (2013).

29. Butovsky, O. et al. Identification of a unique TGF-beta-dependent molecular and functional signature in microglia. Nat Neurosci 17, 131–143, doi:10.1038/nn.3599 (2014).

30. Kim, J. S. et al. A Binary Cre Transgenic Approach Dissects Microglia and CNS Border-Associated Macrophages. Immunity 54, 176–190 e177, doi:10.1016/j.immuni.2020.11.007 (2021).

31. Bennett, M. L. et al. New tools for studying microglia in the mouse and human CNS. Proc Natl Acad Sci U S A 113, E1738–1746, doi:10.1073/pnas.1525528113 (2016).

32. Van Hove, H. et al. A single-cell atlas of mouse brain macrophages reveals unique transcriptional identities shaped by ontogeny and tissue environment. Nat Neurosci 22, 1021–1035, doi:10.1038/s41593-019-0393-4 (2019).

33. Goldmann, T. et al. Origin, fate and dynamics of macrophages at central nervous system interfaces. Nat Immunol 17, 797–805, doi:10.1038/ni.3423 (2016).

34. Li, Q. & Barres, B. A. Microglia and macrophages in brain homeostasis and disease. Nat Rev Immunol 18, 225–242, doi:10.1038/nri.2017.125 (2018).

35. Prinz, M., Masuda, T., Wheeler, M. A. & Quintana, F. J. Microglia and Central Nervous System-Associated Macrophages-From Origin to Disease Modulation. Annu Rev Immunol 39, 251–277, doi:10.1146/annurev-immunol-093019-110159 (2021).

36. Mrdjen, D. et al. High-Dimensional Single-Cell Mapping of Central Nervous System Immune Cells Reveals Distinct Myeloid Subsets in Health, Aging, and Disease. Immunity 48, 599, doi:10.1016/j.immuni.2018.02.014 (2018).

37. Zhan, L. et al. A MAC2-positive progenitor-like microglial population is resistant to CSF1R inhibition in adult mouse brain. Elife 9, doi:10.7554/eLife.51796 (2020).

38. Lawson, L. J., Perry, V. H., Dri, P. & Gordon, S. Heterogeneity in the distribution and morphology of microglia in the normal adult mouse brain. Neuroscience 39, 151–170, doi:10.1016/0306-4522(90)90229-w (1990).

39. Ayata, P. et al. Epigenetic regulation of brain region-specific microglia clearance activity. Nat Neurosci 21, 1049–1060, doi:10.1038/s41593-018-0192-3 (2018).

40. Chistiakov, D. A., Killingsworth, M. C., Myasoedova, V. A., Orekhov, A. N. & Bobryshev, Y.V. CD68/macrosialin: not just a histochemical marker. Lab Invest 97, 4–13, doi:10.1038/labinvest.2016.116 (2017).

41. Lein, E. S. et al. Genome-wide atlas of gene expression in the adult mouse brain. Nature 445, 168–176, doi:10.1038/nature05453 (2007).

42. Stevens, B. et al. The classical complement cascade mediates CNS synapse elimination. Cell 131, 1164–1178, doi:10.1016/j.cell.2007.10.036 (2007).

43. Schafer, D. P. et al. Microglia sculpt postnatal neural circuits in an activity and complement-dependent manner. Neuron 74, 691–705, doi:10.1016/j.neuron.2012.03.026 (2012).

44. Rahpeymai, Y. et al. Complement: a novel factor in basal and ischemia-induced neurogenesis. EMBO J 25, 1364–1374, doi:10.1038/sj.emboj.7601004 (2006).

45. Haynes, S. E. et al. The P2Y12 receptor regulates microglial activation by extracellular nucleotides. Nat Neurosci 9, 1512–1519, doi:10.1038/nn1805 (2006).

46. Stence, N., Waite, M. & Dailey, M. E. Dynamics of microglial activation: a confocal time-lapse analysis in hippocampal slices. Glia 33, 256–266 (2001).

47. Masuda, T. et al. Novel Hexb-based tools for studying microglia in the CNS. Nat Immunol 21, 802–815, doi:10.1038/s41590-020-0707-4 (2020).

48. van Wageningen, T. A. et al. Regulation of microglial TMEM119 and P2RY12 immunoreactivity in multiple sclerosis white and grey matter lesions is dependent on their inflammatory environment. Acta Neuropathol Commun 7, 206, doi:10.1186/s40478-019-0850-z (2019).

49. Yuan, J. et al. M2 microglia promotes neurogenesis and oligodendrogenesis from neural stem/progenitor cells via the PPARgamma signaling pathway. Oncotarget 8, 19855–19865, doi:10.18632/oncotarget.15774 (2017).

50. Yang, Y. et al. MiR-124 Enriched Exosomes Promoted the M2 Polarization of Microglia and Enhanced Hippocampus Neurogenesis After Traumatic Brain Injury by Inhibiting TLR4 Pathway. Neurochem Res 44, 811–828, doi:10.1007/s11064-018-02714-z (2019).

51. Choi, J. Y. et al. M2 Phenotype Microglia-derived Cytokine Stimulates Proliferation and Neuronal Differentiation of Endogenous Stem Cells in Ischemic Brain. Exp Neurobiol 26, 33–41, doi:10.5607/en.2017.26.1.33 (2017).

52. Krasemann, S. et al. The TREM2-APOE Pathway Drives the Transcriptional Phenotype of Dysfunctional Microglia in Neurodegenerative Diseases. Immunity 47, 566–581 e569, doi:10.1016/j.immuni.2017.08.008 (2017).

53. Sierra, A. et al. Microglia shape adult hippocampal neurogenesis through apoptosis-coupled phagocytosis. Cell Stem Cell 7, 483–495, doi:10.1016/j.stem.2010.08.014 (2010).

54. Wakselman, S. et al. Developmental neuronal death in hippocampus requires the microglial CD11b integrin and DAP12 immunoreceptor. J Neurosci 28, 8138–8143, doi:10.1523/JNEUROSCI.1006-08.2008 (2008).

55. Yagi, S. & Galea, L. A. M. Sex differences in hippocampal cognition and neurogenesis. Neuropsychopharmacology 44, 200–213, doi:10.1038/s41386-018-0208-4 (2019).

56. Yagi, S. et al. Sex Differences in Maturation and Attrition of Adult Neurogenesis in the Hippocampus. eNeuro 7, doi:10.1523/ENEURO.0468-19.2020 (2020).

57. Sala Frigerio, C. et al. The Major Risk Factors for Alzheimer’s Disease: Age, Sex, and Genes Modulate the Microglia Response to Abeta Plaques. Cell Rep 27, 1293–1306 e1296, doi:10.1016/j.celrep.2019.03.099 (2019).

58. Villa, A. et al. Sex-Specific Features of Microglia from Adult Mice. Cell Rep 23, 3501–3511, doi:10.1016/j.celrep.2018.05.048 (2018).

59. Bennett, M. L. & Viaene, A. N. What are activated and reactive glia and what is their role in neurodegeneration? Neurobiol Dis 148, 105172, doi:10.1016/j.nbd.2020.105172 (2021).

60. Eggen, B. J., Raj, D., Hanisch, U. K. & Boddeke, H. W. Microglial phenotype and adaptation. J Neuroimmune Pharmacol 8, 807–823, doi:10.1007/s11481-013-9490-4 (2013).

