## Supplemental Data for "Unique Microglial Transcriptomic Signature within the Hippocampal Neurogenic Niche"

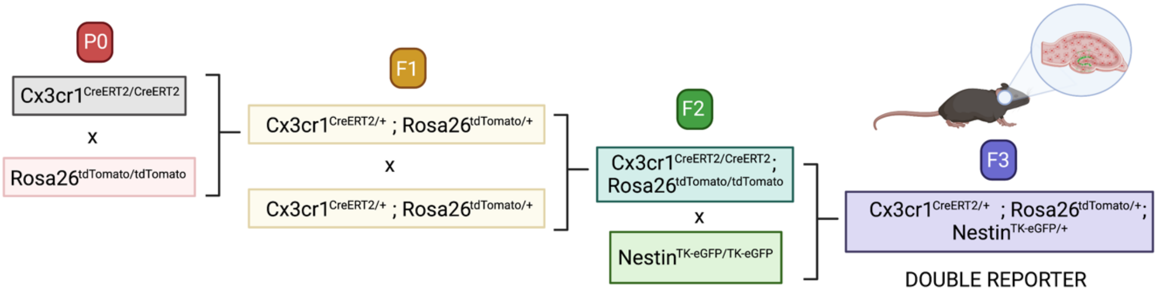

**Supplemental Figure 1**: Generation of double reporter mouse line. Schematic displaying breeding schemes and selection of breeders to generate mice expressing eGFP in Nestin+ neural progenitor cells and tdTomato in cells from fractalkine (Cx3Cr1) expressing cells and their daughter cells.

| | Cells pre-filter | Mean reads | Mean genes | Mean % mito genes | Cells post-filter | Mean reads post filter | Mean genes post filter | Mean % mito genes |
|---|---|---|---|---|---|---|---|---|
| Run 1 | 7255 | 4706 | 2014 | 3,47 | 6810 | 4885 | 2090 | 2.84 |
| Run 2 | 6895 | 3585 | 1621 | 2.49 | 5734 | 3935 | 1749 | 2.25 |
| Run 3 | 6226 | 3891 | 1792 | 3.09 | 5654 | 4109 | 1876 | 2.75 |

**Supplemental Table 1**: Quality control (QC) metrics pre and post- processing. Table displaying cells, mean transcript reads, mean number of unique genes, and the mean percentage of reads corresponding to mitochondrial genes in each sequencing sample.

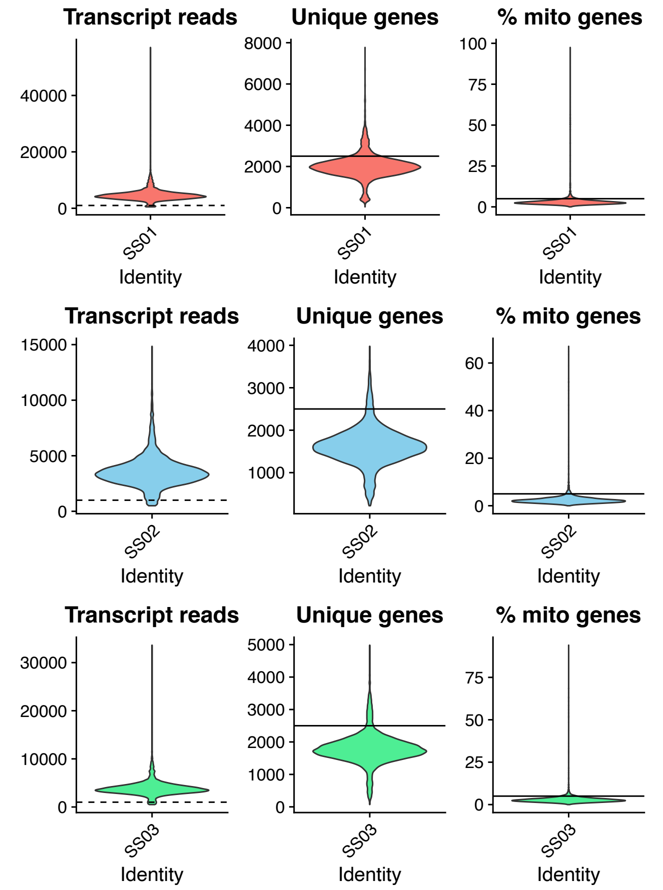

**Supplemental Figure 2**: Quality control (QC) metrics pre-processing. Violin Plots displaying transcript reads, number of unique genes, and the percentage of reads corresponding to mitochondrial genes in each sequencing sample. Dashed line corresponds to lower limit for filtering cells, solid lines represent maximum values for filtering.

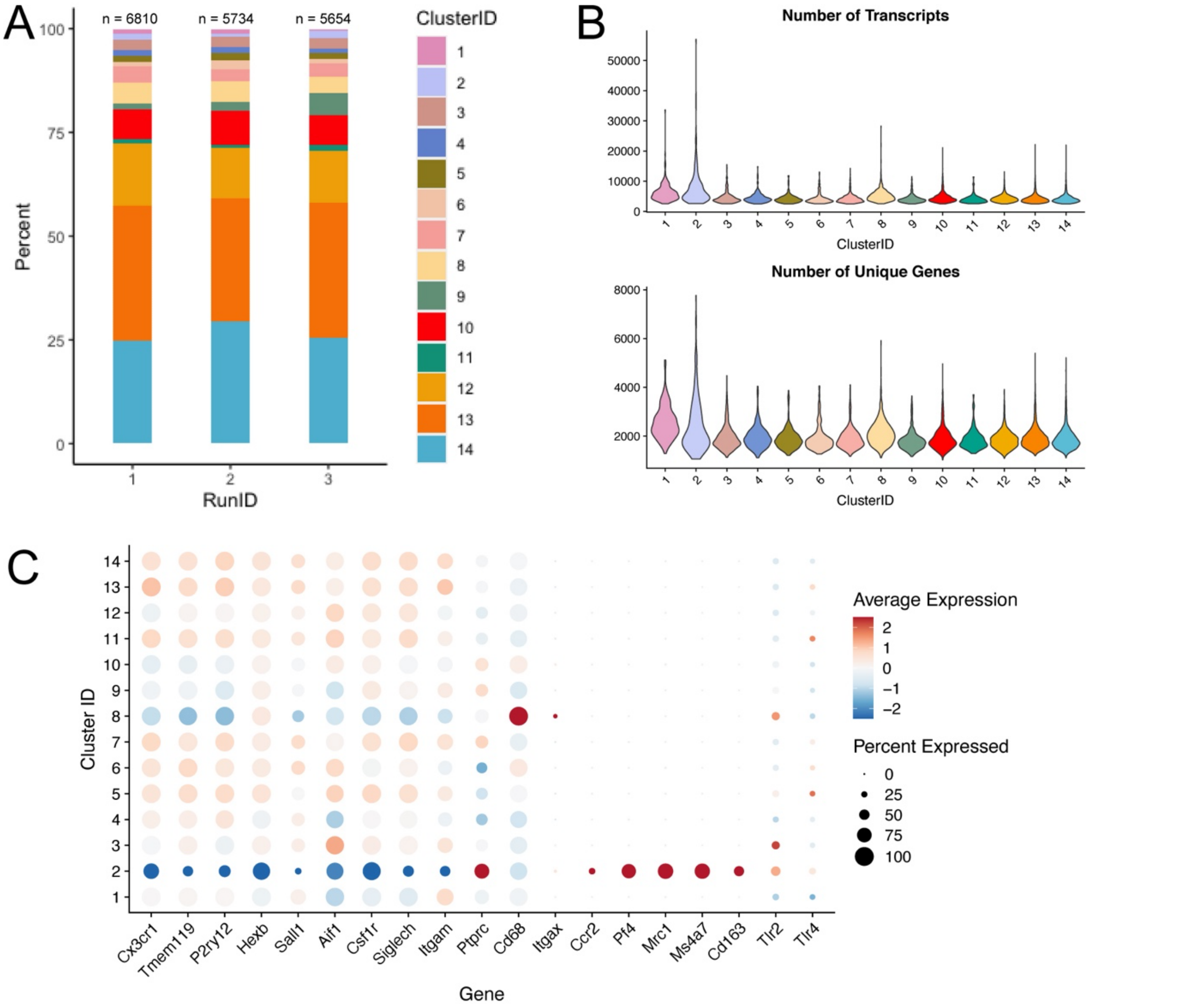

**<u>Supplemental Figure 3: Cluster analysis.</u>** Breakdown of subpopulation frequencies within clusters. (B) number of reads (top) and unique genes (bottom). (C) Dot Plot of myeloid lineage genes. (D) Breakdown of genes differentially expressed genes (highlighting sex-spcific clustering) in homeostatic clusters.

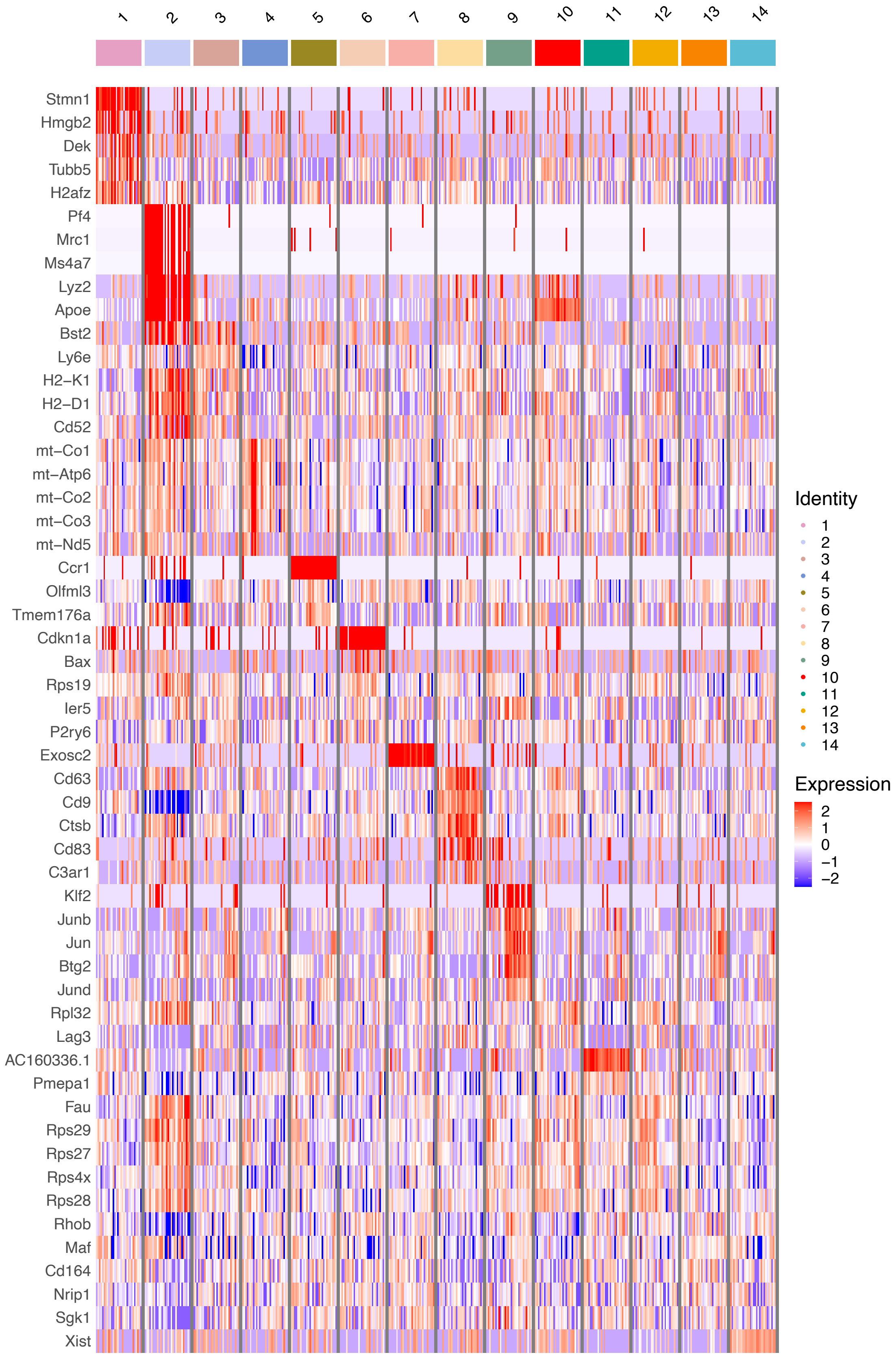

1
2
3
4
5
6
7
8
9
10
11
12
13
14
Stmn1
Hmgb2
Dek
Tubb5
H2afz
Pf4
Mrc1
Ms4a7
Lyz2
Apoe
Bst2
Ly6e
H2−K1
H2−D1
Cd52
mt−Co1
mt−Atp6
mt−Co2
mt−Co3
mt−Nd5
Ccr1
Olfml3
Tmem176a
Cdkn1a
Bax
Rps19
Ier5
P2ry6
Exosc2
Cd63
Cd9
Ctsb
Cd83
C3ar1
Klf2
Junb
Jun
Btg2
Jund
Rpl32
Lag3
AC160336.1
Pmepa1
Fau
Rps29
Rps27
Rps4x
Rps28
Rhob
Maf
Cd164
Nrip1
Sgk1
Xist
Identity
1
2
3
4
5
6
7
8
9
10
11
12
13
14
Expression
2
1
0
−1
−2

**<u>Supplemental Figure 4:</u>** Heat Map displaying scaled expression levels of top five or fewer differentially expressed genes based on set thresholds ($P_{adj}$ <0.001 and expressed in at least 70% of cells in cluster). Thirty cells were sampled for each cluster. Each vertical line represents scaled z-score value of gene expression across row.

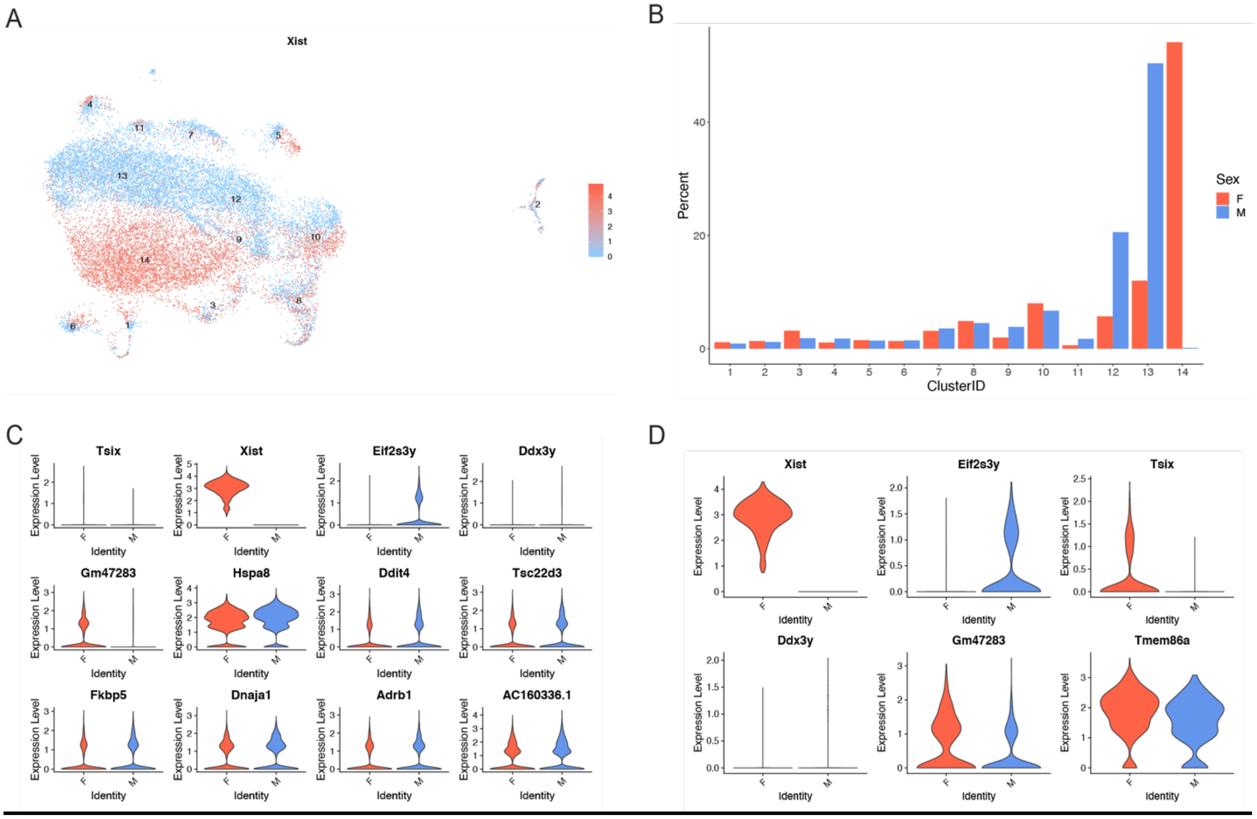

**Supplemental Figure 5:** Sex specific differences in hippocampal myeloid cells. (A) Feature plot displaying scaled expression of *Xist* projected onto UMAP. (B) Distribution of cells *Xist* $^{+}$ female cells versus *Xist* $^{-}$ male cells across clusters. (C) Violin Plots displaying genes differentially expressed between bulk female (F) and Male (M) cells. (D) Violin plots displaying genes differentially expressed between female and male cells in SGZ cluster 8.

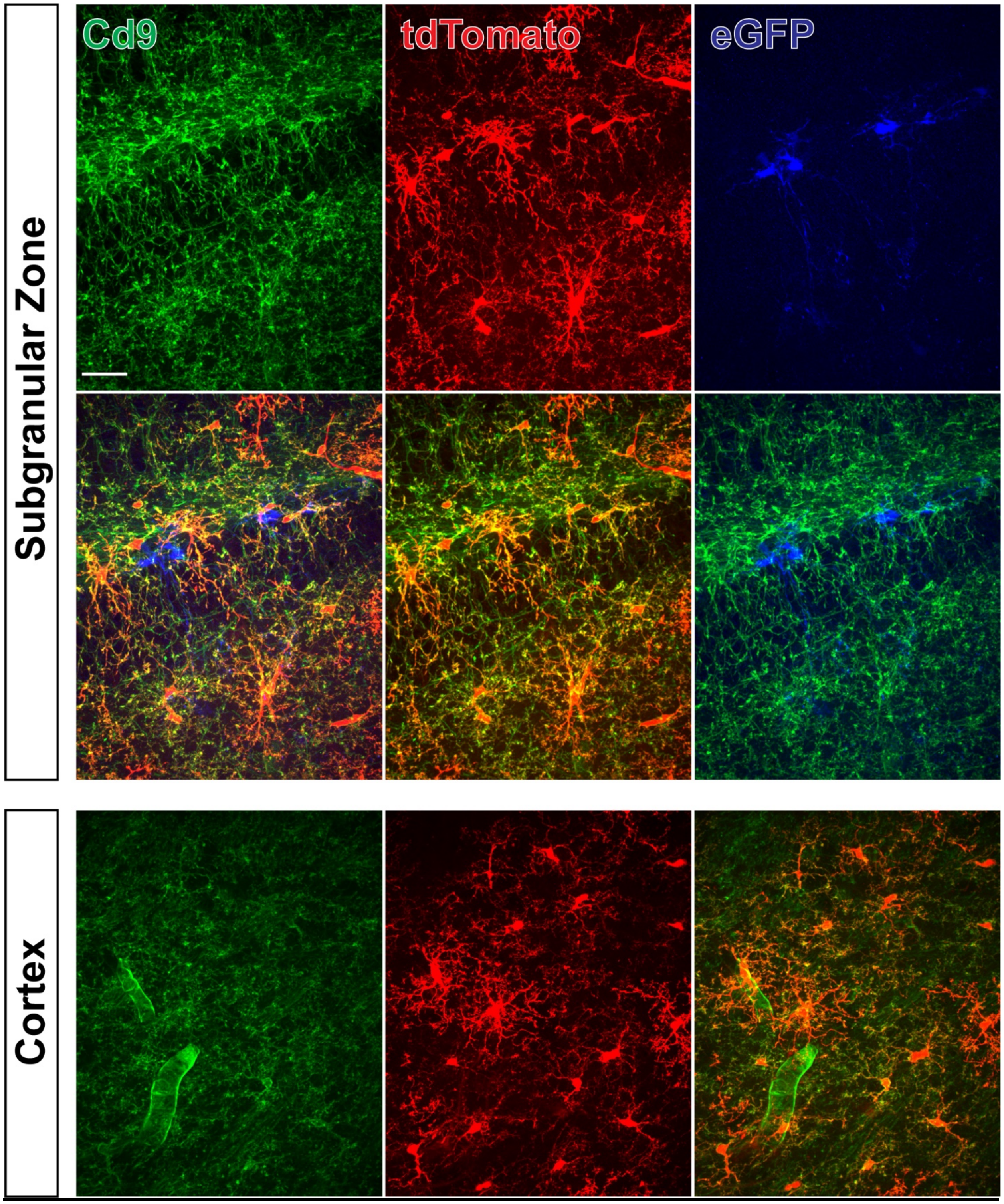

**Supplemental Figure 6: Cd9 immunoreactivity in dual reporter mice in.** Individual channels (top) show expression of Cd9, tdTomato+ myeloid cells and eGFP+ neural progenitor cells. Merge images (middle) show colocalization of Cd9 in both tdtomato and GFP positive cells (left middle). Diffuse immunoreactivity in cortex (bottom) Scale = 10 μm.

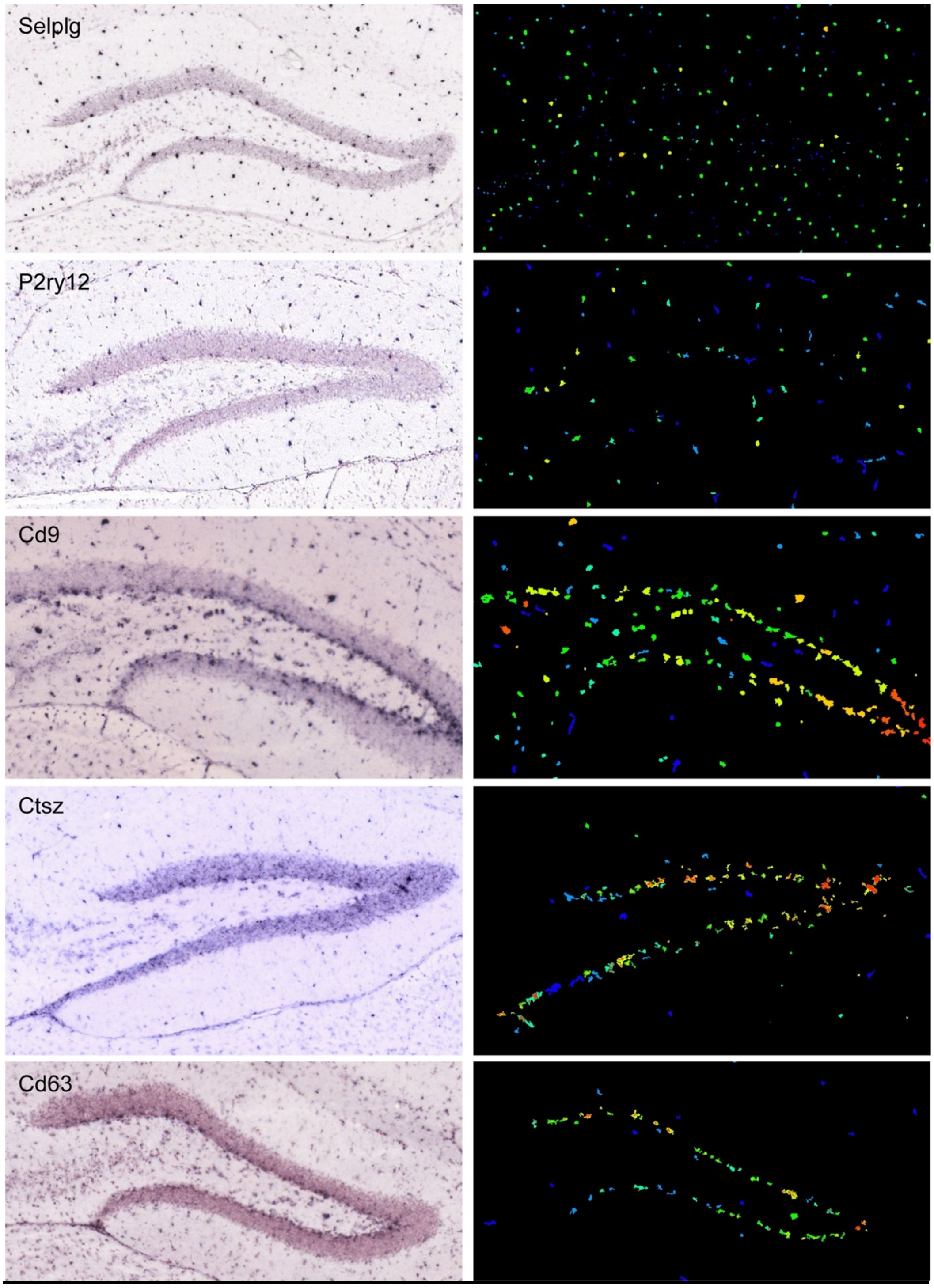

**<u>Supplemental Figure 7: Distribution of differentially expressed genes in Cluster 8 in the dentate gyrus.</u>** *In situ hybridization* images from the Allen Institute of genes downregulated (top two) and upregulated (bottom three).

**Supplemental Table 2: Gene ontology analysis of genes enriched in SGZ vs homeostatic clusters.**

| | GO.ID | Term | Annotated | Significant | Expected | Rank in classicFisher | classicFisher | classicKS | elimKS |
|---|---|---|---|---|---|---|---|---|---|
| 1 | GO:0010033 | response to organic substance | 92 | 92 | 92 | 2 | 1 | 0.00037 | 0.00037 |
| 2 | GO:0045087 | innate immune response | 28 | 28 | 28 | 3 | 1 | 0.00041 | 0.00041 |
| 3 | GO:0008284 | positive regulation of cell population proliferation | 31 | 31 | 31 | 4 | 1 | 0.00047 | 0.00047 |
| 4 | GO:0044283 | small molecule biosynthetic process | 21 | 21 | 21 | 5 | 1 | 0.00049 | 0.00049 |
| 5 | GO:0001819 | positive regulation of cytokine production | 24 | 24 | 24 | 6 | 1 | 0.00050 | 0.00050 |
| 6 | GO:0060429 | epithelium development | 21 | 21 | 21 | 7 | 1 | 0.00068 | 0.00068 |
| 7 | GO:0030335 | positive regulation of cell migration | 27 | 27 | 27 | 8 | 1 | 5.7E-06 | 0.00071 |
| 8 | GO:0048732 | gland development | 14 | 14 | 14 | 9 | 1 | 0.00118 | 0.00118 |
| 9 | GO:0000902 | cell morphogenesis | 26 | 26 | 26 | 10 | 1 | 0.00148 | 0.00148 |
| 10 | GO:0031349 | positive regulation of defense response | 11 | 11 | 11 | 11 | 1 | 0.00157 | 0.00157 |
| 11 | GO:0050727 | regulation of inflammatory response | 14 | 14 | 14 | 12 | 1 | 0.00204 | 0.00204 |
| 12 | GO:0050920 | regulation of chemotaxis | 12 | 12 | 12 | 13 | 1 | 0.00208 | 0.00208 |
| 13 | GO:0002694 | regulation of leukocyte activation | 19 | 19 | 19 | 14 | 1 | 0.00215 | 0.00215 |
| 14 | GO:0002687 | positive regulation of leukocyte migration | 10 | 10 | 10 | 15 | 1 | 0.00239 | 0.00239 |
| 15 | GO:0007166 | cell surface receptor signaling pathway | 57 | 57 | 57 | 16 | 1 | 0.00243 | 0.00243 |
| 16 | GO:0030595 | leukocyte chemotaxis | 17 | 17 | 17 | 17 | 1 | 0.00262 | 0.00262 |
| 17 | GO:0016485 | protein processing | 11 | 11 | 11 | 18 | 1 | 0.00298 | 0.00298 |
| 18 | GO:0050866 | negative regulation of cell activation | 15 | 15 | 15 | 19 | 1 | 0.00317 | 0.00317 |
| 19 | GO:0006954 | inflammatory response | 30 | 30 | 30 | 20 | 1 | 2.7E-05 | 0.00350 |
| 20 | GO:0065008 | regulation of biological quality | 126 | 126 | 126 | 21 | 1 | 0.00184 | 0.00355 |
| 21 | GO:0043523 | regulation of neuron apoptotic process | 19 | 19 | 19 | 22 | 1 | 0.00379 | 0.00379 |
| 22 | GO:0009967 | positive regulation of signal transduction | 45 | 45 | 45 | 23 | 1 | 0.00385 | 0.00385 |
| 23 | GO:0009605 | response to external stimulus | 77 | 77 | 77 | 24 | 1 | 1.1E-05 | 0.00391 |
| 24 | GO:0022603 | regulation of anatomical structure morphogenesis | 36 | 36 | 36 | 25 | 1 | 0.00075 | 0.00445 |
| 25 | GO:0048729 | tissue morphogenesis | 12 | 12 | 12 | 26 | 1 | 0.00538 | 0.00538 |

| | | | | | | | | | |
|---|---|---|---|---|---|---|---|---|---|
| **26** | GO:2000026 | regulation of multicellular organismal development | 63 | 63 | 63 | 27 | 1 | 0.00205 | 0.00543 |
| **27** | GO:0035295 | tube development | 34 | 34 | 34 | 28 | 1 | 0.00109 | 0.00552 |
| **28** | GO:0043066 | negative regulation of apoptotic process | 39 | 39 | 39 | 29 | 1 | 0.00575 | 0.00575 |
| **29** | GO:0045861 | negative regulation of proteolysis | 10 | 10 | 10 | 30 | 1 | 0.00577 | 0.00577 |
| **30** | GO:0043065 | positive regulation of apoptotic process | 25 | 25 | 25 | 31 | 1 | 0.00587 | 0.00587 |
| **31** | GO:0097530 | granulocyte migration | 11 | 11 | 11 | 32 | 1 | 0.00612 | 0.00612 |
| **32** | GO:0045766 | positive regulation of angiogenesis | 12 | 12 | 12 | 33 | 1 | 0.00612 | 0.00612 |
| **33** | GO:0043085 | positive regulation of catalytic activity | 38 | 38 | 38 | 34 | 1 | 0.00118 | 0.00690 |
| **34** | GO:0032270 | positive regulation of cellular protein metabolic process | 48 | 48 | 48 | 35 | 1 | 0.00749 | 0.00749 |
| **35** | GO:0051235 | maintenance of location | 15 | 15 | 15 | 36 | 1 | 0.00765 | 0.00765 |
| **36** | GO:0006165 | nucleoside diphosphate phosphorylation | 10 | 10 | 10 | 37 | 1 | 0.00841 | 0.00841 |
| **37** | GO:0050867 | positive regulation of cell activation | 11 | 11 | 11 | 38 | 1 | 0.00864 | 0.00864 |
| **38** | GO:0002831 | regulation of response to biotic stimulus | 13 | 13 | 13 | 39 | 1 | 0.00882 | 0.00882 |
| **39** | GO:0051347 | positive regulation of transferase activity | 13 | 13 | 13 | 40 | 1 | 0.00897 | 0.00897 |
| **40** | GO:0042127 | regulation of cell population proliferation | 48 | 48 | 48 | 41 | 1 | 1.4E-05 | 0.00914 |
| **41** | GO:0006090 | pyruvate metabolic process | 10 | 10 | 10 | 42 | 1 | 0.00914 | 0.00914 |
| **42** | GO:0045595 | regulation of cell differentiation | 53 | 53 | 53 | 43 | 1 | 0.00919 | 0.00919 |
| **43** | GO:0022008 | neurogenesis | 50 | 50 | 50 | 44 | 1 | 0.00973 | 0.00973 |
| **44** | GO:0072503 | cellular divalent inorganic cation homeostasis | 16 | 16 | 16 | 45 | 1 | 0.01042 | 0.01042 |
| **45** | GO:0001503 | ossification | 12 | 12 | 12 | 46 | 1 | 0.01071 | 0.01071 |
| **46** | GO:0001906 | cell killing | 12 | 12 | 12 | 47 | 1 | 0.01135 | 0.01135 |
| **47** | GO:0002683 | negative regulation of immune system process | 19 | 19 | 19 | 48 | 1 | 0.01138 | 0.01138 |
| **48** | GO:0071621 | granulocyte chemotaxis | 10 | 10 | 10 | 49 | 1 | 0.01178 | 0.01178 |
| **49** | GO:1990266 | neutrophil migration | 10 | 10 | 10 | 50 | 1 | 0.01178 | 0.01178 |
| **50** | GO:0045597 | positive regulation of cell differentiation | 38 | 38 | 38 | 51 | 1 | 0.01242 | 0.01242 |
| **51** | GO:0051128 | regulation of cellular component organization | 73 | 73 | 73 | 52 | 1 | 0.01246 | 0.01246 |
| **52** | GO:0048646 | anatomical structure formation involved in morphogenesis | 31 | 31 | 31 | 53 | 1 | 0.00156 | 0.01250 |

| | | | | | | | | | |
|---|---|---|---|---|---|---|---|---|---|
| **53** | GO:0033674 | positive regulation of kinase activity | 11 | 11 | 11 | 54 | 1 | 0.01264 | 0.01264 |
| **54** | GO:0071310 | cellular response to organic substance | 76 | 76 | 76 | 55 | 1 | 0.01271 | 0.01271 |
| **55** | GO:0007399 | nervous system development | 66 | 66 | 66 | 56 | 1 | 0.00049 | 0.01299 |
| **56** | GO:0030030 | cell projection organization | 41 | 41 | 41 | 57 | 1 | 0.01318 | 0.01318 |
| **57** | GO:0002009 | morphogenesis of an epithelium | 10 | 10 | 10 | 58 | 1 | 0.01366 | 0.01366 |
| **58** | GO:0022610 | biological adhesion | 34 | 34 | 34 | 59 | 1 | 0.01372 | 0.01372 |
| **59** | GO:0051960 | regulation of nervous system development | 34 | 34 | 34 | 60 | 1 | 0.01372 | 0.01372 |
| **60** | GO:0009888 | tissue development | 43 | 43 | 43 | 61 | 1 | 6.8E-06 | 0.01461 |
| **61** | GO:0050678 | regulation of epithelial cell proliferation | 13 | 13 | 13 | 62 | 1 | 0.01473 | 0.01473 |
| **62** | GO:0051249 | regulation of lymphocyte activation | 13 | 13 | 13 | 63 | 1 | 0.01497 | 0.01497 |
| **63** | GO:0051094 | positive regulation of developmental process | 51 | 51 | 51 | 64 | 1 | 0.00374 | 0.01520 |
| **64** | GO:0002696 | positive regulation of leukocyte activation | 10 | 10 | 10 | 65 | 1 | 0.01600 | 0.01600 |
| **65** | GO:0031399 | regulation of protein modification process | 45 | 45 | 45 | 66 | 1 | 0.01652 | 0.01652 |
| **66** | GO:0048869 | cellular developmental process | 94 | 94 | 94 | 67 | 1 | 0.00032 | 0.01709 |
| **67** | GO:0051651 | maintenance of location in cell | 10 | 10 | 10 | 68 | 1 | 0.01740 | 0.01740 |
| **68** | GO:0002695 | negative regulation of leukocyte activation | 11 | 11 | 11 | 69 | 1 | 0.01750 | 0.01750 |
| **69** | GO:0007167 | enzyme linked receptor protein signaling pathway | 24 | 24 | 24 | 70 | 1 | 0.01762 | 0.01762 |
| **70** | GO:0042742 | defense response to bacterium | 10 | 10 | 10 | 71 | 1 | 0.01814 | 0.01814 |
| **71** | GO:0031175 | neuron projection development | 31 | 31 | 31 | 72 | 1 | 0.01845 | 0.01845 |
| **72** | GO:0002366 | leukocyte activation involved in immune response | 10 | 10 | 10 | 73 | 1 | 0.01847 | 0.01847 |
| **73** | GO:0042325 | regulation of phosphorylation | 44 | 44 | 44 | 74 | 1 | 0.01885 | 0.01885 |
| **74** | GO:0040012 | regulation of locomotion | 35 | 35 | 35 | 75 | 1 | 8.2E-06 | 0.01903 |
| **75** | GO:0010562 | positive regulation of phosphorus metabolic process | 34 | 34 | 34 | 76 | 1 | 0.01908 | 0.01908 |
| **76** | GO:0045937 | positive regulation of phosphate metabolic process | 34 | 34 | 34 | 77 | 1 | 0.01908 | 0.01908 |
| **77** | GO:0051962 | positive regulation of nervous system development | 23 | 23 | 23 | 78 | 1 | 0.01919 | 0.01919 |

| | | | | | | | | | |
|---|---|---|---|---|---|---|---|---|---|
| **78** | GO:0045859 | regulation of protein kinase activity | 17 | 17 | 17 | 79 | 1 | 0.01942 | 0.01942 |
| **79** | GO:0072507 | divalent inorganic cation homeostasis | 17 | 17 | 17 | 80 | 1 | 0.01942 | 0.01942 |
| **80** | GO:0030182 | neuron differentiation | 40 | 40 | 40 | 81 | 1 | 0.02021 | 0.02021 |
| **81** | GO:1901362 | organic cyclic compound biosynthetic process | 53 | 53 | 53 | 82 | 1 | 0.02023 | 0.02023 |
| **82** | GO:0050790 | regulation of catalytic activity | 57 | 57 | 57 | 83 | 1 | 0.00013 | 0.02059 |
| **83** | GO:0003013 | circulatory system process | 17 | 17 | 17 | 84 | 1 | 0.02080 | 0.02080 |
| **84** | GO:0008015 | blood circulation | 17 | 17 | 17 | 85 | 1 | 0.02080 | 0.02080 |
| **85** | GO:0030154 | cell differentiation | 91 | 91 | 91 | 86 | 1 | 0.00036 | 0.02083 |
| **86** | GO:0002521 | leukocyte differentiation | 18 | 18 | 18 | 87 | 1 | 0.02085 | 0.02085 |
| **87** | GO:0043549 | regulation of kinase activity | 18 | 18 | 18 | 88 | 1 | 0.02085 | 0.02085 |
| **88** | GO:0050808 | synapse organization | 13 | 13 | 13 | 89 | 1 | 0.02105 | 0.02105 |
| **89** | GO:0061564 | axon development | 13 | 13 | 13 | 90 | 1 | 0.02105 | 0.02105 |
| **90** | GO:0071345 | cellular response to cytokine stimulus | 30 | 30 | 30 | 91 | 1 | 0.02121 | 0.02121 |
| **91** | GO:0016052 | carbohydrate catabolic process | 14 | 14 | 14 | 92 | 1 | 0.02128 | 0.02128 |
| **92** | GO:0060627 | regulation of vesicle-mediated transport | 21 | 21 | 21 | 93 | 1 | 0.02137 | 0.02137 |
| **93** | GO:0009966 | regulation of signal transduction | 70 | 70 | 70 | 94 | 1 | 0.00031 | 0.02144 |
| **94** | GO:0007155 | cell adhesion | 33 | 33 | 33 | 95 | 1 | 0.02147 | 0.02147 |
| **95** | GO:0070887 | cellular response to chemical stimulus | 98 | 98 | 98 | 96 | 1 | 0.00264 | 0.02216 |
| **96** | GO:0031401 | positive regulation of protein modification process | 33 | 33 | 33 | 97 | 1 | 0.02219 | 0.02219 |
| **97** | GO:0048583 | regulation of response to stimulus | 96 | 96 | 96 | 98 | 1 | 3E-05 | 0.02314 |
| **98** | GO:0009617 | response to bacterium | 23 | 23 | 23 | 99 | 1 | 0.02316 | 0.02316 |
| **99** | GO:1901575 | organic substance catabolic process | 63 | 63 | 63 | 100 | 1 | 0.02410 | 0.02410 |
| **100** | GO:0002449 | lymphocyte mediated immunity | 10 | 10 | 10 | 101 | 1 | 0.02455 | 0.02455 |
| **101** | GO:0006874 | cellular calcium ion homeostasis | 15 | 15 | 15 | 102 | 1 | 0.02463 | 0.02463 |
| **102** | GO:0043524 | negative regulation of neuron apoptotic process | 15 | 15 | 15 | 103 | 1 | 0.02463 | 0.02463 |
| **103** | GO:0048468 | cell development | 57 | 57 | 57 | 104 | 1 | 0.02478 | 0.02478 |
| **104** | GO:0032102 | negative regulation of response to external stimulus | 13 | 13 | 13 | 105 | 1 | 0.02501 | 0.02501 |

| | | | | | | | | |
|---|---|---|---|---|---|---|---|---|
| **105** | GO:0051480 | regulation of cytosolic calcium ion concentration | 13 | 13 | 13 | 106 | 1 | 0.02539 | 0.02539 |
| **106** | GO:0045785 | positive regulation of cell adhesion | 15 | 15 | 15 | 107 | 1 | 0.02569 | 0.02569 |
| **107** | GO:0000904 | cell morphogenesis involved in differentiation | 18 | 18 | 18 | 108 | 1 | 0.02613 | 0.02613 |
| **108** | GO:0051174 | regulation of phosphorus metabolic process | 50 | 50 | 50 | 109 | 1 | 0.02622 | 0.02622 |
| **109** | GO:0051047 | positive regulation of secretion | 20 | 20 | 20 | 110 | 1 | 0.02634 | 0.02634 |
| **110** | GO:0120036 | plasma membrane bounded cell projection organization | 37 | 37 | 37 | 111 | 1 | 0.02695 | 0.02695 |
| **111** | GO:0019438 | aromatic compound biosynthetic process | 50 | 50 | 50 | 112 | 1 | 0.02708 | 0.02708 |
| **112** | GO:0032879 | regulation of localization | 84 | 84 | 84 | 113 | 1 | 0.00357 | 0.02717 |
| **113** | GO:0006163 | purine nucleotide metabolic process | 16 | 16 | 16 | 114 | 1 | 0.02733 | 0.02733 |
| **114** | GO:0009150 | purine ribonucleotide metabolic process | 16 | 16 | 16 | 115 | 1 | 0.02733 | 0.02733 |
| **115** | GO:0009259 | ribonucleotide metabolic process | 16 | 16 | 16 | 116 | 1 | 0.02733 | 0.02733 |
| **116** | GO:0072521 | purine-containing compound metabolic process | 16 | 16 | 16 | 117 | 1 | 0.02733 | 0.02733 |
| **117** | GO:0002697 | regulation of immune effector process | 17 | 17 | 17 | 118 | 1 | 0.02763 | 0.02763 |
| **118** | GO:0006950 | response to stress | 113 | 113 | 113 | 119 | 1 | 0.00233 | 0.02847 |
| **119** | GO:0006875 | cellular metal ion homeostasis | 31 | 31 | 31 | 120 | 1 | 0.02903 | 0.02903 |
| **120** | GO:0034654 | nucleobase-containing compound biosynthetic process | 48 | 48 | 48 | 121 | 1 | 0.03007 | 0.03007 |
| **121** | GO:0051336 | regulation of hydrolase activity | 35 | 35 | 35 | 122 | 1 | 0.03025 | 0.03025 |
| **122** | GO:0007420 | brain development | 15 | 15 | 15 | 123 | 1 | 0.03047 | 0.03047 |
| **123** | GO:0048667 | cell morphogenesis involved in neuron differentiation | 15 | 15 | 15 | 124 | 1 | 0.03160 | 0.03160 |
| **124** | GO:0048858 | cell projection morphogenesis | 15 | 15 | 15 | 125 | 1 | 0.03160 | 0.03160 |
| **125** | GO:0045664 | regulation of neuron differentiation | 25 | 25 | 25 | 126 | 1 | 0.03246 | 0.03246 |
| **126** | GO:0048699 | generation of neurons | 46 | 46 | 46 | 127 | 1 | 0.03267 | 0.03267 |
| **127** | GO:0019725 | cellular homeostasis | 46 | 46 | 46 | 128 | 1 | 0.03312 | 0.03312 |
| **128** | GO:0042327 | positive regulation of phosphorylation | 32 | 32 | 32 | 129 | 1 | 0.03317 | 0.03317 |
| **129** | GO:0030155 | regulation of cell adhesion | 22 | 22 | 22 | 130 | 1 | 0.03324 | 0.03324 |

| | | | | | | | | | |
|---|---|---|---|---|---|---|---|---|---|
| **130** | GO:0043412 | macromolecule modification | 70 | 70 | 70 | 131 | 1 | 0.03326 | 0.03326 |
| **131** | GO:0001932 | regulation of protein phosphorylation | 39 | 39 | 39 | 132 | 1 | 0.03333 | 0.03333 |
| **132** | GO:0042063 | gliogenesis | 14 | 14 | 14 | 133 | 1 | 0.03340 | 0.03340 |
| **133** | GO:0019220 | regulation of phosphate metabolic process | 49 | 49 | 49 | 134 | 1 | 0.03370 | 0.03370 |
| **134** | GO:0050673 | epithelial cell proliferation | 15 | 15 | 15 | 135 | 1 | 0.03397 | 0.03397 |
| **135** | GO:0006464 | cellular protein modification process | 67 | 67 | 67 | 136 | 1 | 0.03408 | 0.03408 |
| **136** | GO:0036211 | protein modification process | 67 | 67 | 67 | 137 | 1 | 0.03408 | 0.03408 |
| **137** | GO:0045860 | positive regulation of protein kinase activity | 10 | 10 | 10 | 138 | 1 | 0.03582 | 0.03582 |
| **138** | GO:0050877 | nervous system process | 24 | 24 | 24 | 139 | 1 | 0.03647 | 0.03647 |
| **139** | GO:0030334 | regulation of cell migration | 33 | 33 | 33 | 140 | 1 | 2E-06 | 0.03675 |
| **140** | GO:0065009 | regulation of molecular function | 72 | 72 | 72 | 141 | 1 | 0.00046 | 0.03678 |
| **141** | GO:0060284 | regulation of cell development | 36 | 36 | 36 | 142 | 1 | 0.03678 | 0.03678 |
| **142** | GO:0048666 | neuron development | 34 | 34 | 34 | 143 | 1 | 0.03726 | 0.03726 |
| **143** | GO:0010720 | positive regulation of cell development | 26 | 26 | 26 | 144 | 1 | 0.03775 | 0.03775 |
| **144** | GO:0010646 | regulation of cell communication | 84 | 84 | 84 | 145 | 1 | 0.00068 | 0.03777 |
| **145** | GO:0023051 | regulation of signaling | 84 | 84 | 84 | 146 | 1 | 0.00068 | 0.03777 |
| **146** | GO:1902105 | regulation of leukocyte differentiation | 11 | 11 | 11 | 147 | 1 | 0.03789 | 0.03789 |
| **147** | GO:2000145 | regulation of cell motility | 35 | 35 | 35 | 148 | 1 | 8.2E-06 | 0.03955 |
| **148** | GO:0008285 | negative regulation of cell population proliferation | 17 | 17 | 17 | 149 | 1 | 0.03979 | 0.03979 |
| **149** | GO:0002263 | cell activation involved in immune response | 11 | 11 | 11 | 150 | 1 | 0.03987 | 0.03987 |
| **150** | GO:0002573 | myeloid leukocyte differentiation | 10 | 10 | 10 | 151 | 1 | 0.04051 | 0.04051 |
| **151** | GO:0030278 | regulation of ossification | 10 | 10 | 10 | 152 | 1 | 0.04116 | 0.04116 |
| **152** | GO:0003006 | developmental process involved in reproduction | 18 | 18 | 18 | 153 | 1 | 0.04134 | 0.04134 |
| **153** | GO:0010959 | regulation of metal ion transport | 15 | 15 | 15 | 154 | 1 | 0.04290 | 0.04290 |
| **154** | GO:1901615 | organic hydroxy compound metabolic process | 20 | 20 | 20 | 155 | 1 | 0.04301 | 0.04301 |
| **155** | GO:0055074 | calcium ion homeostasis | 16 | 16 | 16 | 156 | 1 | 0.04311 | 0.04311 |
| **156** | GO:0051641 | cellular localization | 72 | 72 | 72 | 157 | 1 | 0.04432 | 0.04432 |
| **157** | GO:0018130 | heterocycle biosynthetic process | 49 | 49 | 49 | 158 | 1 | 0.04479 | 0.04479 |

| | | | | | | | | | |
|---|---|---|---|---|---|---|---|---|---|
| **158** | GO:0009719 | response to endogenous stimulus | 35 | 35 | 35 | 159 | 1 | 0.04537 | 0.04537 |
| **159** | GO:0005975 | carbohydrate metabolic process | 23 | 23 | 23 | 160 | 1 | 0.04537 | 0.04537 |
| **160** | GO:0001501 | skeletal system development | 16 | 16 | 16 | 161 | 1 | 0.04609 | 0.04609 |
| **161** | GO:0050767 | regulation of neurogenesis | 31 | 31 | 31 | 162 | 1 | 0.04642 | 0.04642 |
| **162** | GO:0051050 | positive regulation of transport | 38 | 38 | 38 | 163 | 1 | 0.04837 | 0.04837 |
| **163** | GO:0005996 | monosaccharide metabolic process | 10 | 10 | 10 | 164 | 1 | 0.04991 | 0.04991 |

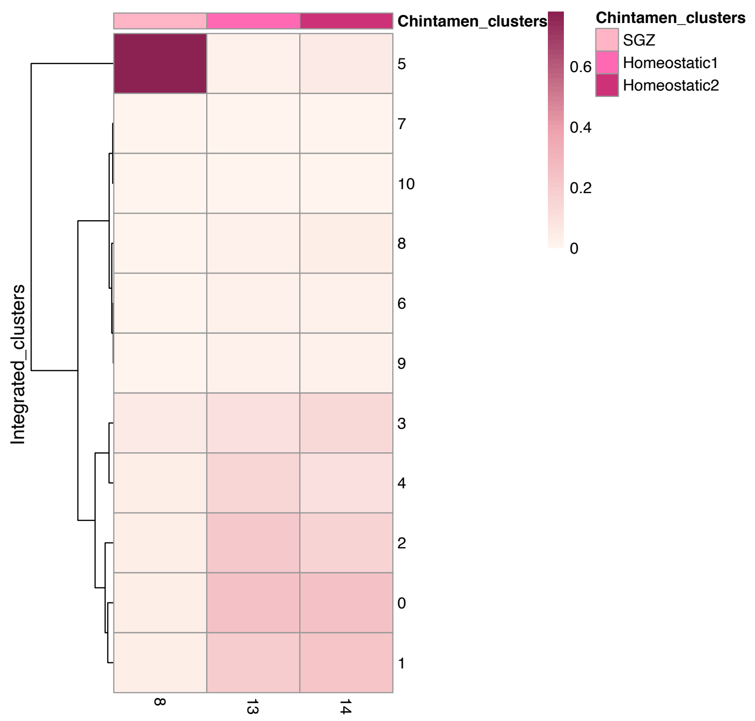

**Supplemental Figure 8: Correlation Matrix for original clusters versus integrated DAM dataset.** Plot showing relation between clusters 8,13,14 from this study and integrated dataset from Keren-Shaul et. al. 2017. SGZ cluster shows overlap with cluster 5 in the integrated dataset. Cluster 5.
